# A novel peptidoglycan deacetylase modulates daughter cell separation in *E. coli*

**DOI:** 10.1101/2025.02.18.638797

**Authors:** Victor Hernandez-Rocamora, Alessandra M. Martorana, Aitana Belloso, Daniel Ballesteros, Marta Zaccaria, Amilcar J. Perez, Bogdan I. Iorga, David Abia, Joe Gray, Eefjan Breukink, Jie Xiao, Manuel Pazos, Alessandra Polissi, Waldemar Vollmer

## Abstract

Peptidoglycan hydrolases facilitate bacterial cell wall growth by creating space for insertion of new material and allowing physical separation of daughter cells. In *Escherichia coli*, three peptidoglycan amidases, AmiA, AmiB and AmiC, cleave septal peptidoglycan during cell division. The LytM-domain proteins EnvC and NlpD activate these amidases either from inside the cell or the outer membrane: EnvC binds to the cytoplasmic membrane-anchored divisome components FtsEX, and NlpD and ActS are outer membrane lipoproteins. Here we report the identification of a novel periplasmic deacetylase called SddA that removes acetyl groups from denuded peptidoglycan glycan strands, the products of amidases. SddA is a substrate for the periplasmic protease Prc, suggesting regulation via protein degradation. The *sddA* gene is co-expressed with the gene encoding EnvC, linking SddA function to amidase activation. Consistent with this link, the deletion of *sddA* alleviates phenotypes associated with lack of amidase activation, while overexpression of *sddA* alleviates phenotypes related to a defective Tol-Pal system and causes cell chaining due to reduced septum peptidoglycan cleavage unless *envC* is co-expressed. We present a model according to which SddA modulates the activation of the septum-splitting amidases during cell division.

**AUTHOR SUMMARY:** Bacteria surround their cell membrane by the essential peptidoglycan (cell wall) layer to prevent bursting open due to their turgor. During cell division, bacteria produce a septum at midcell, which must be cleaved for daughter cells to separate. Here, we report the identification of a new enzyme, SddA, that modifies a particular type of peptidoglycan material, denuded glycan chains released during the splitting of septal peptidoglycan for daughter cell separation, in the Gram-negative *Escherichia coli*. We propose a model in which SddA modulates a switch in the septal peptidoglycan splitting, ensuring splitting is activated from the cell membrane in the early stages of cell division and from the outer membrane in the late stages.

## INTRODUCTION

The bacterial peptidoglycan (PG) sacculus provides support against the turgor and maintains the shape of the cell [1]. PG is made of chains of alternating *N*-acetylglucosamine (Glc*N*Ac) and *N*-acetylmuramic acid (Mur*N*Ac) residues with short peptides linked to Mur*N*Ac. Peptides from adjacent strands can be connected by crosslinks to form the PG network surrounding the cytoplasmic membrane [1]. Rod-shaped bacteria such as *E. coli* use membrane-spanning multiprotein complexes for sacculus growth: the divisome facilitates cell division at midcell and the elongasome side wall expansion [2, 3]. Both complexes coordinate the activities of various PG-synthesising and hydrolysing enzymes and PG biogenesis is coordinated with outer membrane (OM) synthesis and maintenance [2, 4, 5].

Cell division is a key process in the cell cycle in which PG synthesis and hydrolysis are coupled. The cytoplasmic tubulin homologue FtsZ forms treadmilling FtsZ polymers rotating around the division site [6–8], attached to the cytoplasmic membrane (CM) by FtsA and ZipA [9]. Early in cell division, PG synthesis is initiated by class A Penicillin-binding proteins (PBPs) [10, 11], regulated by SPOR-domain PG-binding proteins such as FtsN [11, 12]. Assembly of the complete divisome activates the major cell division-specific synthases FtsW-PBP3 [3]. PG synthases polymerize glycan strands from the PG precursor lipid II and cross-link the peptides [13]. PG hydrolases are needed for splitting the septal PG for the separation of the daughter cells. PG *N*-acetylmuramoyl-L-alanine amidases (amidases) remove the stem peptides from the glycan strands producing ’denuded’ strands. In *E. coli*, AmiA, AmiB and AmiC are the major septum-splitting enzymes responsible for daughter cell separation [14, 15]. They are periplasmic enzymes with structurally-related catalytic domains and regulated by LytM-domain-containing activators [16]. EnvC activates AmiA and AmiB and is itself activated by the cytoplasmic membrane-anchored FtsEX proteins, which are members of the divisome [17–20]. By contrast, the lipoproteins NlpD and ActS activate AmiC from the OM [21–23]. The regulation of the NlpD/AmiC complex is poorly understood and might involve the Tol-Pal system, but not DolP [24]. The transmembrane Tol-Pal system facilitates OM constriction during cell division and regulates, via the accessory protein CpoB, the activation of the transpeptidase (TPase) of the PG synthase PBP1B [4].

Knockout of either the amidases or their regulators, or members of the Tol-Pal system, produces chaining and OM-defects in *E. coli* [14, 16, 25].

Many bacteria modify the chemical structure of peptidoglycan to adapt to different growth conditions or to resist attacks by hydrolases released by the host or bacteriophages [1, 26]. One important modification is the deacetylation of the sugars in the glycan chain, which helps bacteria to resist the digestion of PG by lysozyme. The first PG *N*-acetylglucosamine deacetylase, PgdA, was identified in *Streptococcus pneumoniae* [27], where it contributes to virulence [28], and homologues are present in other Gram-positive bacteria, for example *Listeria monocytogenes* [29] or *Streptococcus suis* [30]. The PgdA catalytic domain contains a NodB-homology domain (PFAM PF01522) with a conserved triad of two histidine and one aspartic residue coordinating Zn^2+^ as metal cofactor [31]. PG deacetylases have been thought to be absent in Gram-negative bacteria. The ’atypical’ PG deacetylase suggested to be present in a Gram-negative bacterium, PgdA from *Helicobacter pylori* [32, 33], seems to lack a signal peptide for export into the periplasm, and deacetylated PG fragments were not observed in *H. pylori* [34–37]. Interestingly, a recent high-resolution mass spectrometry-based approach identified a small amount of deacetylated PG glycan fragments (Glc*N*-Mur*N*Ac, S1B Fig) [38] in *E. coli*, although *E. coli* does not appear to have a homologue of PgdA and deacetylated disaccharide peptides (muropeptides) were not released from *E. coli* PG by a muramidase [39, 40].

In this work, we report the discovery of a new enzyme in *E. coli* that can deacetylate denuded PG glycan strands produced by amidases. The deacetylase, which we named SddA (<u>S</u>eptal <u>d</u>enuded strand <u>d</u>eacetylase <u>A</u>), is genetically linked to the amidase activator EnvC and localizes to the cell division septum. SddA is distantly related to PgdA structurally as both contain catalytic domains belonging to the Glycoside hydrolase/deacetylase, beta/alpha-barrel superfamily (Interpro IPR011330). Our results suggest that SddA is involved in regulating amidase activity during cell division by affecting the activation of amidases from the OM or CM.

## RESULTS

### SddA is a denuded glycan strand deacetylase

We searched the *E. coli* genome for genes encoding for periplasmic proteins with homology to PG deacetylases. A fold-based search using *S. pneumoniae* PgdA as query and the *E. coli* proteome as target yielded 8 hits, of which three are proteins targeted for periplasmic export (PgaB, YadE and YibQ). These proteins contain a polysaccharide domain similar to that of PgdA. PgaB is an OM protein that deacetylates the extracellular poly-β-(1→6)-Glc*N*Ac (PGA) matrix for biofilm formation [41, 42]. YadE is a protein of unknown function with genetic links to genes involved in lipid A modification and synthesis [43]. We also identified YibQ, a protein containing a divergent polysaccharide deacetylase domain (protein family PF04748) that is related to but distinct to the deacetylase domain of PgdA (PF01522). Interestingly, the *yibQ* gene locates within the same operon as the PG hydrolase activator gene, *envC* (Fig 1A), and YibQ has a predicted signal peptide for export into the periplasm, making it a candidate for a PG deacetylase. To identify how conserved this genetic linkage is, we determined the distribution of genes encoding for YibQ homologues within the Annotree database which contains a representative set of consistently annotated bacterial genomes [44]. Genes encoding for YibQ homologues are present mainly within genomes of Gram-negative bacteria (Fig 1B), and most of these, including *E. coli yibQ*, are located on the same strand of the DNA (Fig 1C) and within two kilobases to genes encoding for EnvC homologues (Fig 1D). An AlphaFold model of YibQ predicts an N-terminal globular domain with conserved metal binding residues followed by a long C-terminal disordered region (Fig 1E and S2B Fig).

**Fig 1.**
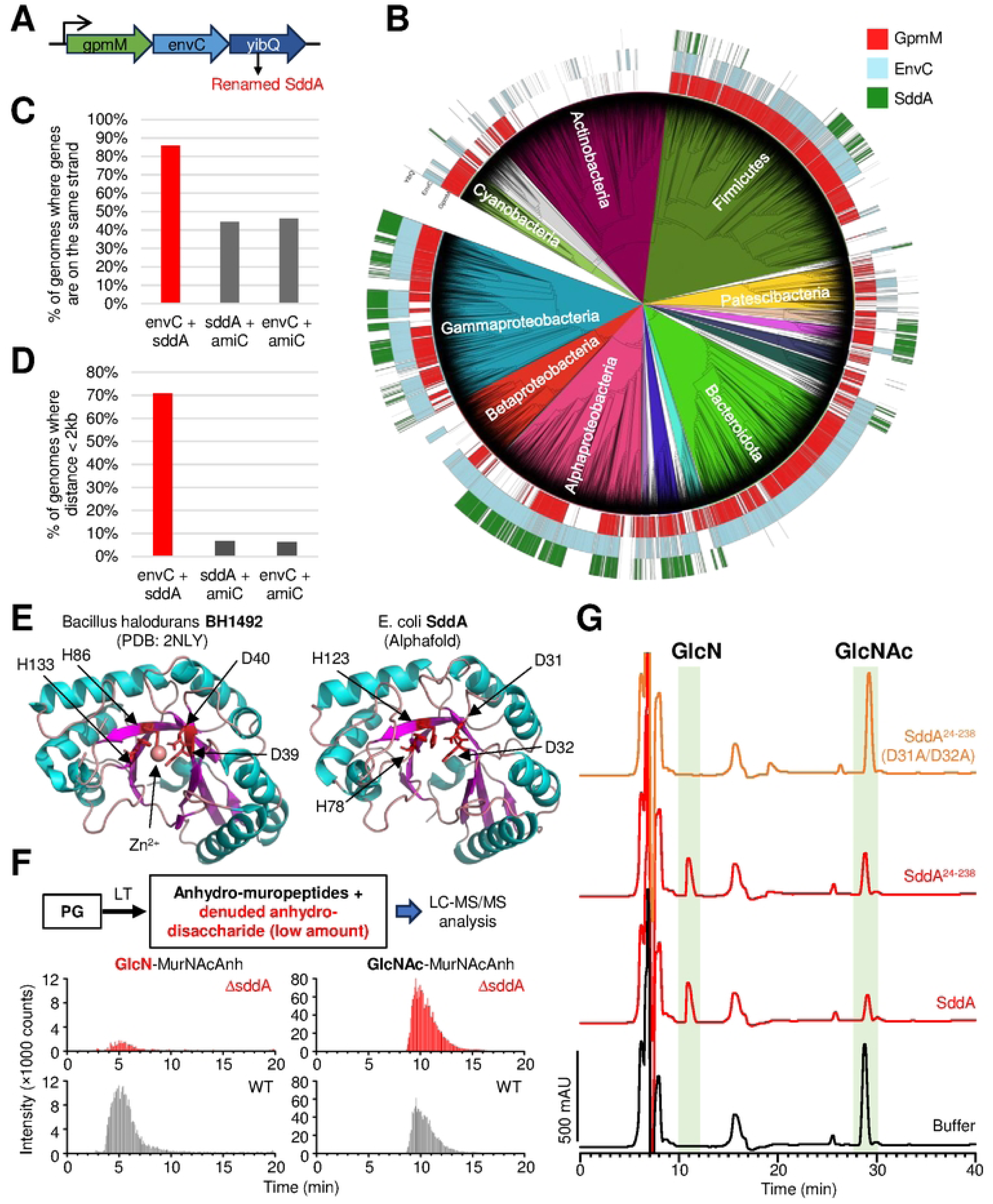
SddA is a denuded PG glycan strand deacetylase genetically linked to amidase activator EnvC. **(A)** The *E. coli* gene *yibQ* (*sddA*) is located downstream in the same operon as amidase activator *envC*. **(B)** Distribution of *gpmM*, *envC* and *sddA* homologues in the bacterial genome database Annotree. **(C)** When *sddA* and *envC* are present in the same genome in Annotree, they are most often present on the same strand. **(D)** When *sddA* and *envC* are present in the same genome in Annotree they are found most often at a distance of less than 2 kilobases. **(E)** Alphafold model of SddA (left), compared with the crystal structure of BH1492 from *Bacillus halodurans* (PDB 2NLY), which also contains a divergent deacetylase domain. The residues involved in Zn^2+^ coordination in the BH1492 structure and their homologues in SddA are indicated. **(F)** Analysis by LC/MS of the presence of Glc*N*-Mur*N*AcAnh and Glc*N*Ac-Mur*N*AcAnh in PG from *E. coli* BW25113 (WT) or *E. coli* BW25113Δ*sddA* (Δ*sddA*), released by MltA. Top, scheme of the procedure, bottom, chromatograms showing intensity of peaks eluting with m/z 437 and 479 corresponding to Glc*N*-Mur*N*AcAnh and Glc*N*Ac-Mur*N*AcAnh, respectively, for WT and Δ*sddA*. The identity of Glc*N*-Mur*N*AcAnh and GlcN*A*c-Mur*N*AcAnh MS peaks was verified by MS/MS fragmentation (Fig S1). In the scheme, LT stands for lytic transglycosylase. **(G)** Chromatograms of the analysis of denuded glycan strands treated first with buffer, SddA, SddA^24-238^ or SddA^24-238^ D31A/D32A, and then with the lytic transglycosylase MltA to produce anhydro-muropeptides. Labelled peaks were identified by mass spectrometry (see text) and correspond to Glc*N*-Mur*N*AcAnh (Glc*N*) and Glc*N*Ac-Mur*N*AcAnh (Glc*N*Ac).

To test if YibQ is the enzyme responsible for the presence of Glc*N*-Mur*N*Ac in *E. coli* PG, we purified PG from *E. coli* BW25113 and an isogenic *yibQ* deletion strain, digested it with the lytic transglycosylase MltA to generate the 1,6-anhydro-disaccharide subunits and analysed these by liquid-chromatography-mass spectrometry (LC-MS/MS) for the presence of Glc*N*-Mur*N*AcAnh and Glc*N*Ac-Mur*N*AcAnh (Fig 1F and S1AB Fig). Glc*N*-Mur*N*AcAnh was present in BW25113 and absent in BW25113 Δ*yibQ* (Fig 1F), suggesting that YibQ is responsible for deacetylating Glc*N*Ac residues in denuded strands. Based on these results and results shown below, we renamed YibQ to SddA (Septal Denuded strand Deacetylase A).

### SddA is a Glc*N*Ac deacetylase specific for denuded strands

Next, we wanted to determine whether the N-terminally oligohistidine-tagged version of SddA lacking the signal peptide has PG deacetylase activity. However, upon His-SddA overexpression, most of the produced protein was insoluble and the soluble cell fraction contained only a truncated version (S2 Fig). We purified the soluble, truncated His-SddA and used mass spectroscopy to identify the truncation sites by determining the molecular weights of the protein variants present in these preparations. This analysis showed that the truncation happened within the C-terminal flexible region and the protein still contained the whole globular catalytic domain (S2A Fig). We thus tested the activity of this version (named herein His-SddA). His-SddA was active against 4-nitrophenyl acetate, a common substrate used to test deacetylase activity, including that of PgdA [45] (S2C Fig), suggesting that the C-terminal flexible region is not required for activity. Therefore, we also expressed and purified a truncated version lacking the predicted C-terminal flexible domain entirely, named His- SddA^24-238^, which was completely soluble when overexpressed.

We next tested the activity of His-SddA against whole PG sacculi or muropeptides, which were obtained by digesting PG with a muramidase. No visible changes in the muropeptide profiles were observed in the presence of the His-SddA (S3 Fig), indicating that SddA is not active against PG or muropeptides. Considering the location of *sddA* next to *envC*, we next tested if His-SddA is active against denuded glycan strands. For this, we developed an assay in which radiolabelled, un-crosslinked PG strands were generated with a TPase-defective PBP1B variant and [^14^C]GlcNAc-labelled lipid II. These strands were then treated with the amidase AmiC in the presence of its activator NlpD to produce radiolabelled denuded glycan strands (S4A Fig). Incubation of these strands with His-SddA prevented their digestion with the muramidase cellosyl (S4B Fig). If the radiolabelled denuded glycan strands were (partially) pre-digested with cellosyl into short glycan strands, His-SddA produced new peaks with different elution times, indicating that SddA modified denuded glycan strands (S4C Fig). To determine the exact reaction performed by SddA, we prepared denuded glycan strands by complete digestion of purified PG with the amidase AmiD and incubated them with His-SddA. The product was digested with lytic transglycosylase MltA, which is able to completely digest denuded glycan strands [46], and analysed the products by HPLC and mass spectrometry (Fig 1G and S1 Fig). Treatment of denuded strands with His-SddA produced a decrease in Glc*N*Ac-Mur*N*AcAnh (neutral mass 478.10 amu) and yielded a new peak, identified by MS/MS as Glc*N*-Mur*N*AcAnh (neutral mass 436.05 amu) (Fig 1G). His-SddA^24-238^ generated the same product, confirming that the flexible C-terminal region in SddA is not required for activity. Finally, His-SddA was also able to deacetylate short Glc*N*Ac-Mur*N*Ac oligomers produced by partial digestion of denuded strands by cellosyl (S5 Fig). Overall, these results demonstrate that His-SddA deacetylates GlcNAc residues in denuded strands.

### Conserved Asp residues in SddA are required for activity

The Protein Data Bank (PDB) includes two structures of a PF04748 domain, i.e., the same family as the catalytic domain in SddA. In one of these, corresponding to BH1492 from *Bacillus halodurans*, a Zn^2+^ ion is coordinated by two Asp and two His residues (Fig 1E). These four residues are conserved in SddA: D31, D32, H78, H123 (S6 Fig). To test if these residues are important for SddA activity, we purified the SddA D31A/D32A version and assayed the activity assay against denuded glycan strands. SddA D31A/D32A showed no activity in this assay (Fig 1G), suggesting that the two hypothetical metal (Zn^2+^)- binding residues are important for SddA activity.

### Prc digests SddA *in vivo*

As the C-terminal flexible part in SddA was cleaved when expressing the protein in the cytoplasm, we asked whether this part of SddA could also be subject to cleavage in the periplasm by the tail-specific protease Prc, which degrades some PG-related proteins in the periplasm [47, 48]. To study this, we expressed SddA versions with an N- or C-terminal FLAG-tag and the native SddA signal peptide (FLAG-SddA or SddA-FLAG). These constructs were expressed from a low-copy number plasmid under control of an IPTG-inducible promoter (S7A Fig). We induced the expression of these constructs with IPTG in BW27783 (WT) and Δ*prc* cells and detected the expressed proteins using western blot and an anti-FLAG antibody (S7B-C Fig). Only one band was detected in western blots with either construct, with the approximate apparent molecular weight expected for the mature, full-length SddA (S7B Fig). In both WT and Δ*prc* backgrounds, SddA-FLAG had a lower signal than FLAG-SddA (S7B Fig), suggesting a lower expression or higher degradation. Alternatively, the anti-FLAG antibody might have a higher affinity for N-terminal FLAG tags. Importantly, both versions of FLAG-tagged SddA were detected at higher levels in the Δ*prc* background (S7B Fig), suggesting that Prc is able to digest SddA *in vivo*. An AlphaFold3 prediction shows that Prc could interact with SddA via the flexible C-terminal region, although additional studies would be required to identify the exact position(s) where the proteolysis may take place (S7D Fig).

### The localization of FtsN SPOR domain is not affected by *sddA* deletion

SPOR domains recognize PG denuded glycan strands allowing proteins bearing these domains to locate to division sites where these strands are produced by amidases [49, 50]. These domains are present in the *E. coli* proteins FtsN, DedD, DamX and RlpA. All of these except RlpA have been shown to interact with or activate PG synthases during cell division, and they contain a single transmembrane helix anchoring them to the cytoplasmic membrane (Fig 2A) [12, 49, 51, 52]. We wondered whether the deacetylation of denuded glycan strands could affect the binding of SPOR domain proteins, thus preventing their midcell localization or function. We first asked if the deletion of *sddA* has an effect in a Δ*dedD* background. Cells lacking *dedD* are more elongated and more sensitive to outer membrane (OM) stressors such as detergents [53]. A Δ*dedD* Δ*sddA* strain showed a reduced sensitivity to deoxycholate compared to Δ*dedD* but was still more sensitive than the WT (Fig 2B). In addition, Δ*dedD* Δ*sddA* cells had similar length as Δ*dedD* (Fig 2C), indicating that the deficient cell division phenotype was not restored by *sddA* deletion.

**Fig 2.**
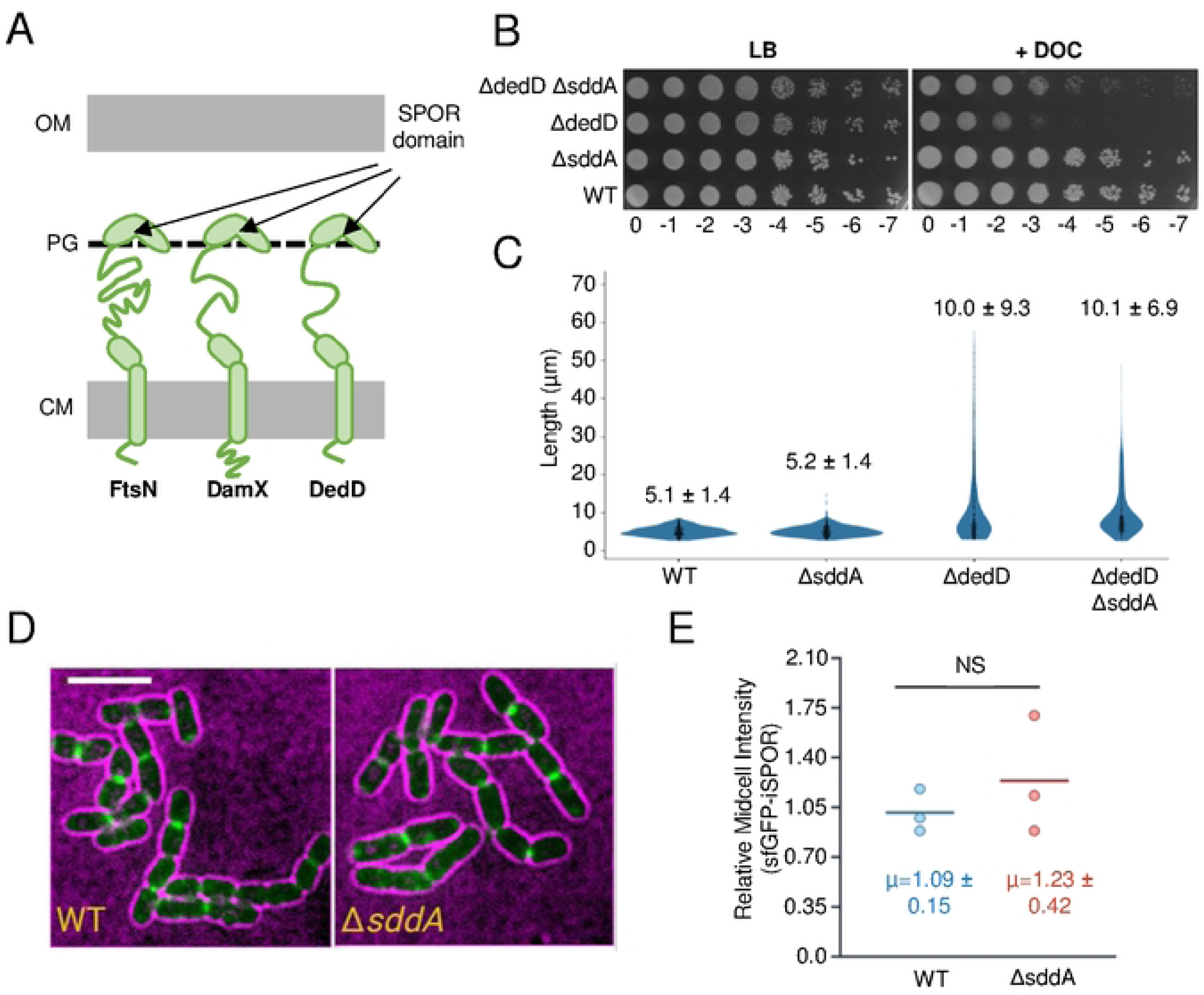
Effect of *sddA* deletion on SPOR domain proteins. **(A**) Scheme showing the three SPOR domain proteins anchored to the cytoplasmic membrane (CM) of *E. coli*, which bind denuded glycan strands at midcell and participate in cell division regulation. **(B)** Ten-fold serial dilutions spotted plate of BW25113 (WT), MP005 (Δ*sddA*), MP001 (Δ*dedD*) and MP006 (Δ*dedD* Δ*sddA*) strain cultures incubated at 30°C in LB-Lennox with or without 0.2% (w/v) sodium deoxycholate (DOC). **(C)** Violin plot showing cell length of the strains in **(B)**, grown at 30°C in LB-Lennox in exponential phase. The mean ± standard deviation (SD) is indicated (n>300). **(D)** Representative images of BW25113 (WT) or BW25113 Δ*sddA* (Δ*sddA*) bearing pCH-ss^dsbA^-sfGFP-iSPOR showing the localization of sfGFP-iSPOR (green) in cells (black cells with magenta background). Scale bar is 5 μm. **(E)** Quantification of average midcell intensity of sfGFP-iSPOR. Each data point is representative of the mean average value of 29-52 cells “relative midcell intensities” (normalized by the WT median per replicate, see methods). Written is the mean ± SD from three independent biological replicates. NS means not significant (nonparametric Mann-Whitney test).

To directly probe SPOR protein recruitment in the absence of SddA, we used fluorescence microscopy to localize the isolated SPOR domain of FtsN tagged with a fluorescent protein (sfGFP-iSPOR) in BW25113 or the isogenic strain lacking *sddA* (Fig 2D). The deacetylation of denuded glycan strands might alter the affinity of SPOR domains and, hence, their recruitment to midcell could be different in the absence of SddA. However, we did not observe a significant difference in midcell localization intensity of sfGFP-iSPOR between the WT and Δ*sddA* strains (Fig 2E), suggesting that either SPOR domains are not affected by deacetylation by SddA or that the activity of SddA at midcell is too low to affect SPOR localization. This result suggests SddA is not involved in the regulation of FtsN localization at midcell. Further studies are needed to explore whether SddA affects the localization and function of DedD, DamX, or RlpA.

### SddA function is linked to peptidoglycan amidases

In order to understand the role of SddA in the cell, we investigated phenotypes caused by deleting the *sddA* gene in *E. coli* BW25113. BW25113 Δ*sddA* grew with normal rate and morphology under standard laboratory conditions (S8 Fig), and the lack of *sddA* did not affect OM permeability of mutants lacking one or more amidase activators (S9 Fig). Mutants lacking amidases or amidase activators form chains and become sensitive to detergents such as deoxycholate [54]. We confirmed this phenotype in mutants that depend on either EnvC (Δ*nlpD* Δ*actS*) or NlpD (Δ*envC* Δ*actS*) (Fig 3) for amidase activation. Interestingly, deleting *sddA* in these backgrounds rendered cells less sensitive to deoxycholate, suggesting that SddA might inhibit amidases and removal of SddA enhances amidase activity (Fig 3B). In contrast to the strains depending solely on NlpD or EnvC for amidase activation, the deletion of *sddA* from the strain depending solely on ActS (Δ*envC* Δ*nlpD*) produced a considerable increase in lag phase, although growth rates during exponential phase were similar (S10 Fig). Both, Δ*envC* Δ*nlpD* and Δ*envC* Δ*nlpD* Δ*sddA* produced very long and sometimes twisted cell chains (S10 Fig). Overall, these results suggest that SddA can inhibit PG amidases in some genetic backgrounds. This modulation might be through direct interaction with the amidases and/or their activators, or indirectly through the deacetylated PG product of SddA.

**Fig 3.**
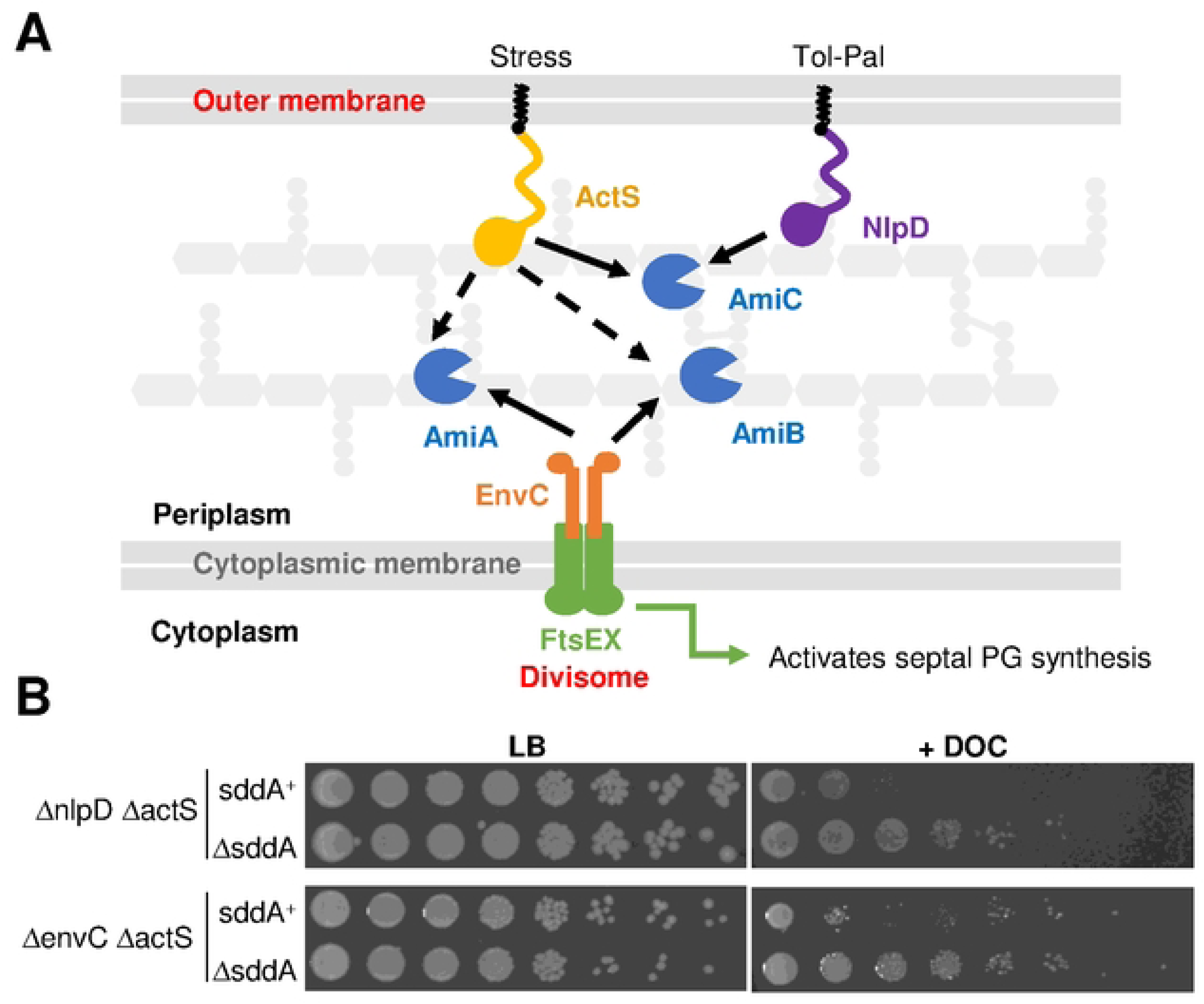
SddA deletion alters deoxycholate sensitivity of strains defective in multiple amidase activators. **(A)** Scheme depicting amidase activation in *E. coli*. AmiA and AmiB are activated by the FtsEX-EnvC complex, which is part of the divisome, from inside the cell. AmiC is activated by lipoproteins NlpD and ActS. ActS can also activate AmiA and AmiB *in vitro* (dashed arrows). **(B)** Ten- fold serial dilutions spotted plates of BW25113 Δ*nlpD* Δ*actS* and BW25113 Δ*nlpD* Δ*actS* Δ*sddA* (top), and BW25113 Δ*envC* Δ*actS*, and BW25113 Δ*envC* Δ*actS* Δ*ssdA* (bottom). Plates were incubated at 37°C. Top plates contain LB with or without 0.5% deoxycholate (DOC). Bottom plates contain LB- Lennox with or without 0.5% deoxycholate (DOC).

### SddA overexpression causes cell chaining and OM defects

As the deletion of *sddA* in a WT background caused no obvious growth phenotype, we tested the effects of SddA overexpression by inserting the full length *sddA* sequence, including its signal peptide, into an expression plasmid under control of IPTG-inducible *Ptac* promoter yielding pGS100-*sddA*. Addition of IPTG to BW25113 bearing pGS100-*sddA* but not the empty plasmid (pGS100) increased the sensitivity to vancomycin and SDS/EDTA (Fig 4A) and induced the formation of long cell chains (Fig 4B), phenocopying the deletion of amidases or their activators [54] and suggesting that PG amidases are impaired upon SddA overexpression. As *sddA* and *envC* are co-expressed in the same operon on the genome, we tested whether co-overexpression of both from a plasmid has the same effects as overexpression of the individual proteins. To test this, we expressed *envC* or both, *envC* and *sddA* from pGS100 in BW25113. IPTG addition to BW25113 cells with pGS100-*envC*-*sddA* or pGS100-*envC* produced no cell chains, in sharp contrast to cells with pGS100-*sddA* (Fig 4B). Co-expression of *nlpD* with *sddA* from pGS100 and pBAD24 plasmids, respectively, did not suppress either sensitivity to vancomycin and SDS/EDTA or the chaining phenotype (S11 Fig), suggesting that only EnvC can alleviate cell chaining caused by excessive SddA. In line with the above, deletion of *envC* or *ftsX* did not rescue the cell chaining caused by SddA overexpression (S12 Fig). Moreover, overexpression of SddA in cells lacking *envC* and its cognate amidases (*amiA* and *amiB*), or *nlpD* and its cognate amidase (*amiC*), aggravated the cell chaining phenotype (S13 Fig). These results indicate that neither amidase system (EnvC-AmiA/B or NlpD-AmiC) is functional upon overexpression of SddA. However, the overproduction of SddA in Δ*envC* Δ*amiA* Δ*amiB* cells produced a sicker phenotype than overproduction of SddA in Δ*nlpD* Δ*amiC* as judged by an extensive cell chaining and highly defective growth phenotype (S13 Fig). These results suggest that SddA inhibits AmiC more than AmiA/B and that the cell needs to balance the expression of SddA and EnvC and achieves this by encoding the corresponding genes in the same operon.

**Fig 4.**
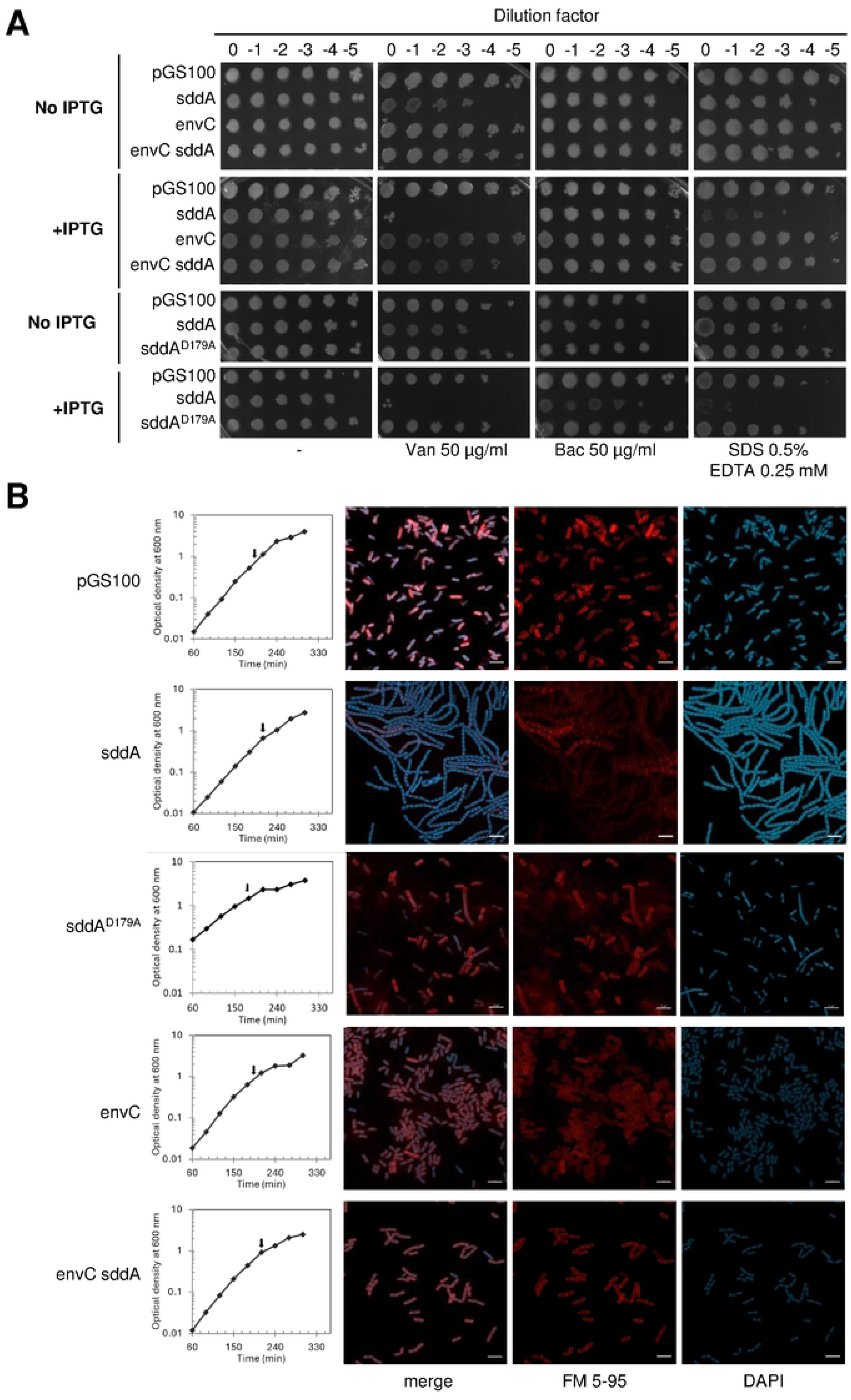
High SddA levels cause OM defects and cell chaining unless *envC* is overexpressed. **(A)** Overnight cultures of BW25113 harbouring pGS100 or pGS100 expressing *sddA, sddA^D179A^*, *envC or* both *envC* and *sddA* were serially diluted and spotted onto LB with 5%NaCl supplemented with 25 µg ml^-1^ chloramphenicol and antibiotics or chemicals at the indicated concentrations, with or without 0.5 mM IPTG. Plates were incubated at 37°C for 24 h. **(B)** BW25113 cells harbouring pGS100 or pGS100 expressing *sddA, sddA^D179A^, envC* or both *envC* and *sddA* were grown in LB-Lennox supplemented with 25 µg ml^-1^ chloramphenicol and 0.5 mM IPTG. Samples were collected at exponential phase (arrows), stained with FM5-95 (red, cell membrane) and DAPI (blue, nucleoid), immobilized and imaged by confocal fluorescence microscopy. Representative images are shown. Scale bar is 5 µm.

### SddA localizes at cell division sites

The cell chaining phenotype caused by overexpressing SddA without EnvC suggests that SddA can reduce the activation or activity of amidases at midcell, preventing daughter cell separation. If this inhibition relies on a direct interaction with septal amidases or their activators, SddA should localize at midcell. To determine the cellular localization, we expressed an SddA-sfGFP fusion containing the native SddA signal peptide. Western blot analysis of cell extracts showed no cleavage of the fluorescent tag, indicating that the SddA-sfGFP fusion is stable (S14 Fig). Overexpression of SddA-sfGFP resulted in cell chaining (as the overexpression of WT protein) and, interestingly, the fluorescence signal localised to the uncleaved septa (Fig 5). Next, we examined the effect of aztreonam on SddA localization. Aztreonam is a β-lactam which inhibits the septal transpeptidase PBP3 [55, 56] but still allows for pre-septal PG synthesis at future cell division sites by class A PBPs, which are recruited by the FtsZ membrane anchors FtsA and ZipA [11]. As expected, aztreonam induced cell filamentation in all strains tested, indicating an inhibition of cell division (S15 Fig). When expressed after aztreonam addition, SddA-sfGFP did not localize at future division sites, indicating that ongoing septum PG synthesis by PBP3 is required for SddA recruitment to midcell (S15 Fig). These results suggest that SddA interacts with one or more divisome proteins, or with septal PG, in a manner dependent on septal PG synthesis.

**Fig 5.**
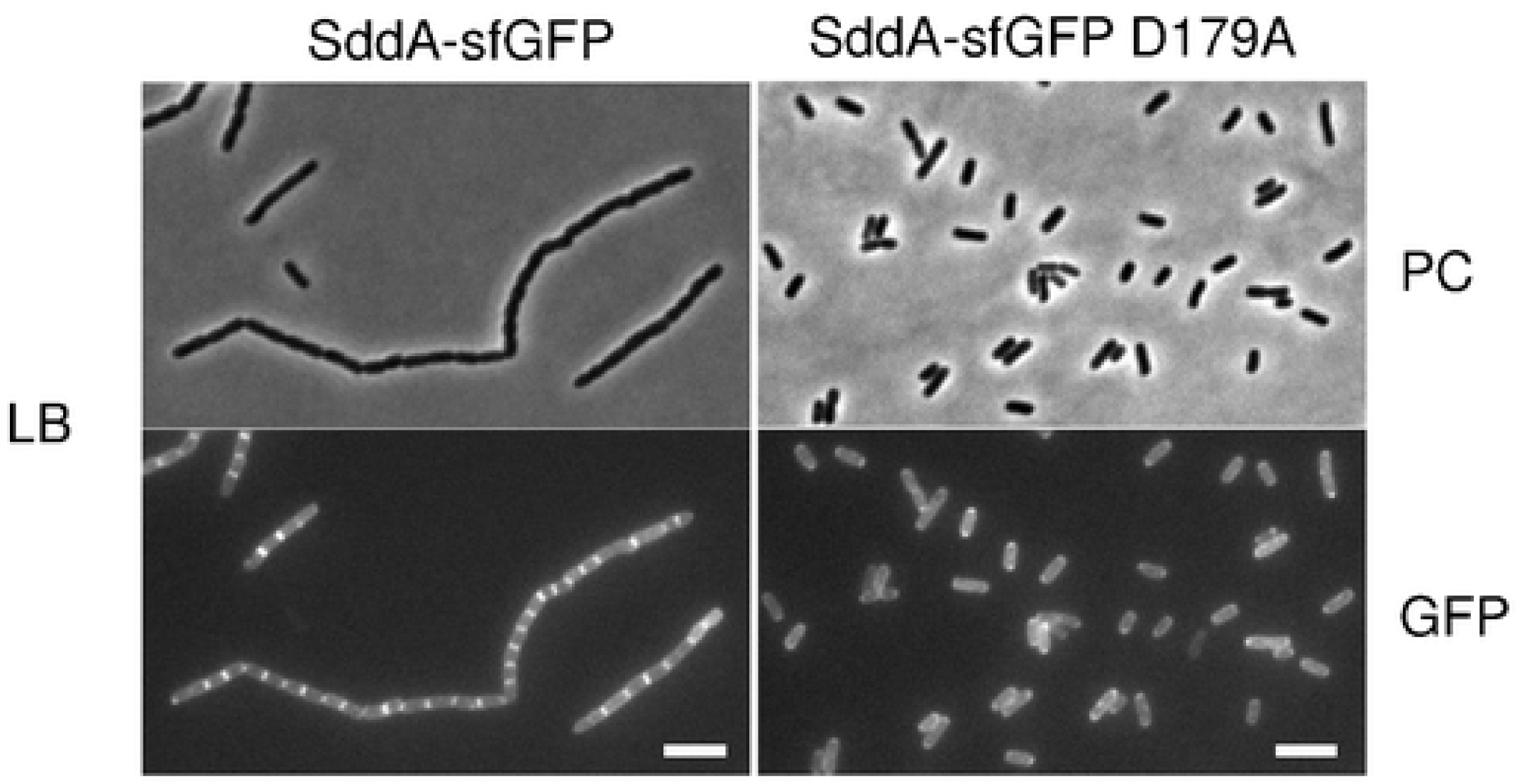
Sdd*A* localizes at the cell division site. BW25113 cells harbouring pGS100 encoding *sddA::sfGFP* or *sddA D179A::sfGFP* were grown in LB with 20 µg ml^-1^ chloramphenicol at 37°C and expression of the fluorescent constructs was induced with 0.5 mM IPTG for 140 min. Samples were imaged by phase contrast (PC) and fluorescence microscopy (GFP). Representative images are shown. Scale bar is 5 µm.

### SddA overexpression phenocopies the removal of septal amidase activators

The above results suggest that SddA localizes at midcell, where it can modulate the activity of EnvC- AmiA/B and/or NlpD-AmiC pairs. If this hypothesis is correct, SddA overexpression should mimic the effects of *envC* or *nlpD* deletion. We tested this hypothesis using *E. coli* mutants defective in the Tol-Pal system, which lyse upon the addition of sub-MIC concentrations of the PBP3-inhibitor aztreonam [57]. We confirmed this phenotype in both, a Δ*tolR* mutant and in cells harbouring TolR D23R, which is unable to harness the proton motive force for the correct functioning of Tol-Pal [58] (Fig 6). The Tol-Pal system coordinates PG synthesis and cell envelope constriction during cell division, suggesting that this phenotype could result from dysregulation of synthetic and hydrolytic PG activities. Additional inactivation of the amidase activators *envC* (or *nlpD*) prevented (or reduced) cell lysis (Fig 6), supporting the idea that amidase activity is misregulated in Tol-Pal*-*defective cells when PBP3 is inhibited by aztreonam. Similarly, the overproduction of SddA (either native or fused to sfGFP) prevented lysis of Δ*tolR* cells in the presence of the PBP3 inhibitor (Fig 6). This further supports the hypothesis that SddA inhibits amidases or their regulators at midcell.

**Fig 6.**
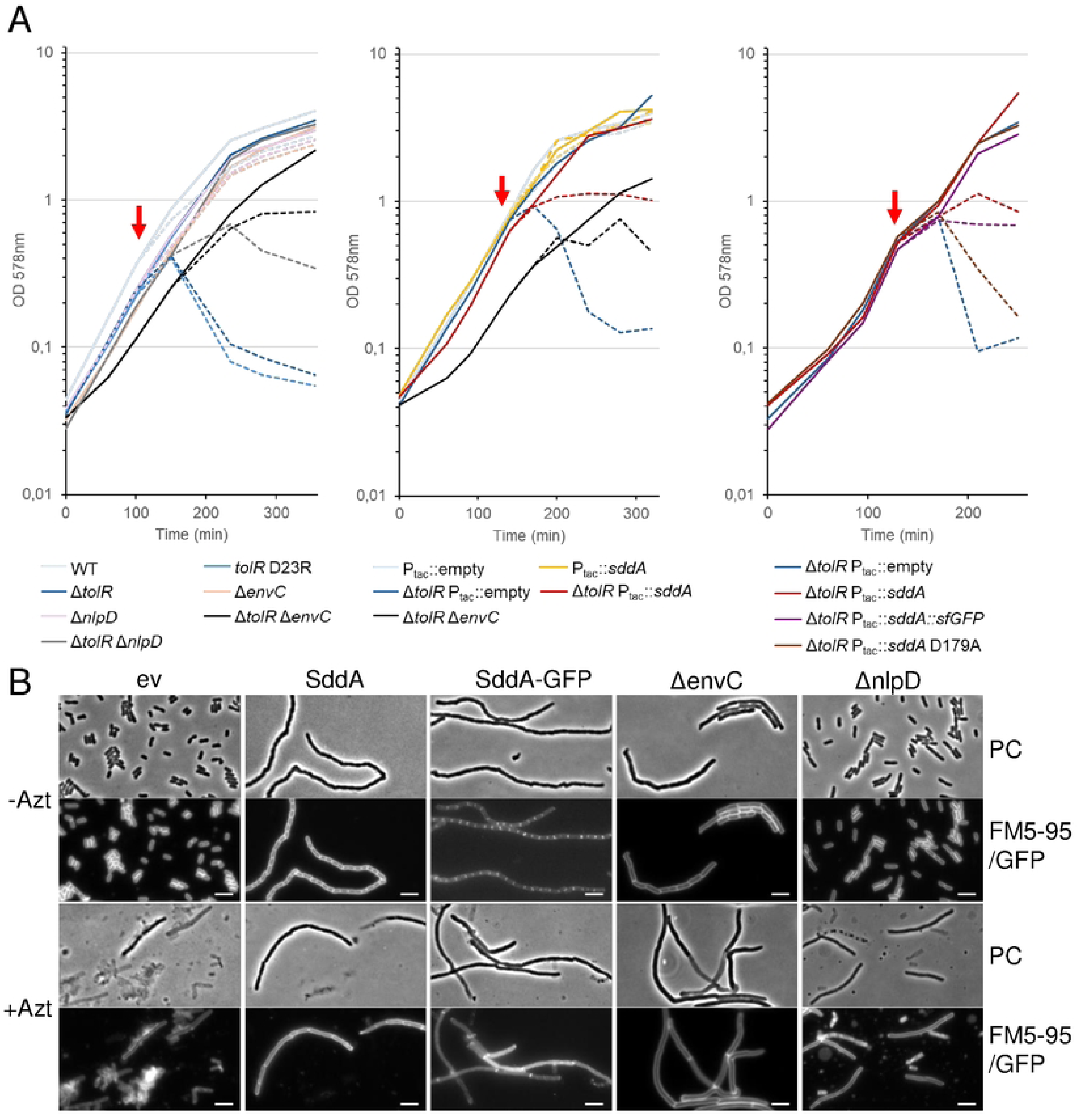
SddA phenocopies the absence of *envC* during the lysis of *tol* defective cells upon septal PG synthesis inhibition. **(A)** BW25113 (WT), BW25113 Δ*tolR*, BW25113 Δ*nlpD*, and BW25113 Δ*tolR* were grown at 37°C in LB with 20 µg ml^-1^ chloramphenicol. When indicated by a red arrow the cultures were split in two halves and 1 µg ml^-1^ aztreonam was added to one of them (dashed line). On the middle and right panels, overexpression of the plasmid-encoded genes was induced by addition of 0.5 mM IPTG when OD578 reached 0.1. **(B)** Representative phase contrast (PC) and fluorescence microscopy (FM5-95, or GFP in case of SddA-GFP) micrographs of samples taken 110 min after splitting cultures (red arrow) of BW25113 Δ*tolR*::FRT containing the empty vector (ev), pGS::*sddA* (SddA) or pMP108 (SddA-GFP), MPW55 (Δ*envC*) and MPW54 (Δ*nlpD*) cells in absence (-Azt) or presence of aztreonam (+Azt). Scale bar is 5 µm.

Since *sddA* is genetically linked to *envC*, and excess production of SddA over EnvC causes cell chaining, we explored potential SddA-EnvC or SddA-FtsX interactions using AlphaFold (Fig S16). AlphaFold predicted an SddA-FtsX complex (score: 0.338), where SddA binds to the periplasmic porter domain of FtsX, and an SddA-EnvC complex (score: 0.530), where SddA interacts with the coiled-coil region of EnvC. Although the FtsX-SddA model had a relatively low confidence score, both FtsX-SddA and EnvC- SddA models predicted that the SddA residues involved in the interactions are located on the same side as the predicted SddA catalytic active site. Moreover, both models predict D179 of SddA to be involved in contacts with FtsX or EnvC. To test this, we overproduced an alanine-179 variant of SddA and analysed its effects on the phenotypes. Overexpression of SddA D179A failed to rescue the lysis of Δ*tolR* in the presence of aztreonam (Fig 6), suggesting that the mutated protein cannot inhibit amidases. In addition, overexpression of SddA D179A or a SddA(D179A)-sfGFP fusion did not induce cell chaining (Fig 5 and S15 Fig). Strikingly, SddA(D179A)-sfGFP failed to localize at midcell (Fig 5 and S17AB Fig). Finally, we introduced the D179A mutation into His-SddA^24-238^ and tested the activity of the purified protein *in vitro*. This variant of SddA remained active against denuded strands (S17C Fig), indicating that the D179A mutation does not affect enzymatic activity, and that the lack of effects upon overproduction is not due to a loss of deacetylase activity.

To understand how the D179A change might disrupt the interaction of SddA with EnvC, we performed molecular dynamics simulations of both WT and D179A, which highlighted the differences in residue fluctuations between them (S18 Fig). SddA residues predicted by AlphaFold to form EnvC contacts include 32, 35–36, 55, 59–60, 85–88, 90, 123–124, 154, 177, 179–180, 182, 209 and 211. The D179A change affected the flexibility of some of those residues, with strong rigidification of residues 35, 36, 58, 59 and 177, potentially hindering adaptive binding, and flexibilization of residues 87 and 124, which may further disrupt the interaction by destabilizing the binding interface. Beyond local effects, D179A induced fluctuation changes in residues 174 and 202, suggesting an allosteric disruption of the interaction; while altered dynamics in regions outside the EnvC interface indicate broader effects on protein flexibility. Catalytic regions were unaffected, with minimal fluctuation differences in residues critical for enzymatic activity, in agreement with the fact that SddA D179A is catalytically active (S17C Fig). Overall, the D179A change may disrupt the SddA-EnvC interaction by altering the dynamics of critical contact residues and their surroundings. The combined changes in rigidity and flexibility appear to compromise conformational adaptability and emphasise the role of residue-specific dynamics in mediating these particular protein-protein interactions.

In summary, these results suggest that SddA interacts with EnvC and/or FtsX, and that the overexpression of SddA (without simultaneous overexpression of EnvC) impairs the activation of AmiA and AmiB.

### SddA also inhibits the activation of AmiC by NlpD

As our results suggest that SddA can inhibit amidase activation at the septum, we tested the effect of SddA on the activity of amidases and their activators *in vitro*. To do this, we performed activity assay with amidase/activator pairs, EnvC-AmiA, ActS-AmiC and NlpD-AmiC, in the presence or absence of His-SddA^24-238^ (Fig 7 and S19 Fig). In the case of EnvC, two versions of this activator were tested: EnvC(LytM), which only contains the AmiA-activating LytM domain, and EnvC(fl), which contains the full mature sequence. Previous work showed that only EnvC(LytM) but not EnvC(fl) activates AmiA and AmiB *in vitro* [21, 59]. In control assays, His-SddA^24-238^ showed no effect on AmiA or AmiC amidases in the absence of their activators (S18C Fig). Surprisingly, His-SddA^24-238^ showed no effect on ActS-AmiC activity or EnvC(fl)-AmiA activity, but there was a small activation of EnvC(LytM)-AmiA (S19 Fig). This could indicate that FtsEX is required for the inhibition of EnvC by SddA. In contrast, His-SddA^24-238^ significantly inhibited NlpD-AmiC activity against PG (Fig 7). Interestingly, inhibition did not depend on the activity of SddA as the catalytically inactive His-SddA^24-238^ D31A/D32A also inhibited NlpD-AmiC, albeit to lower extent than the active protein (Fig 7).

**Fig 7.**
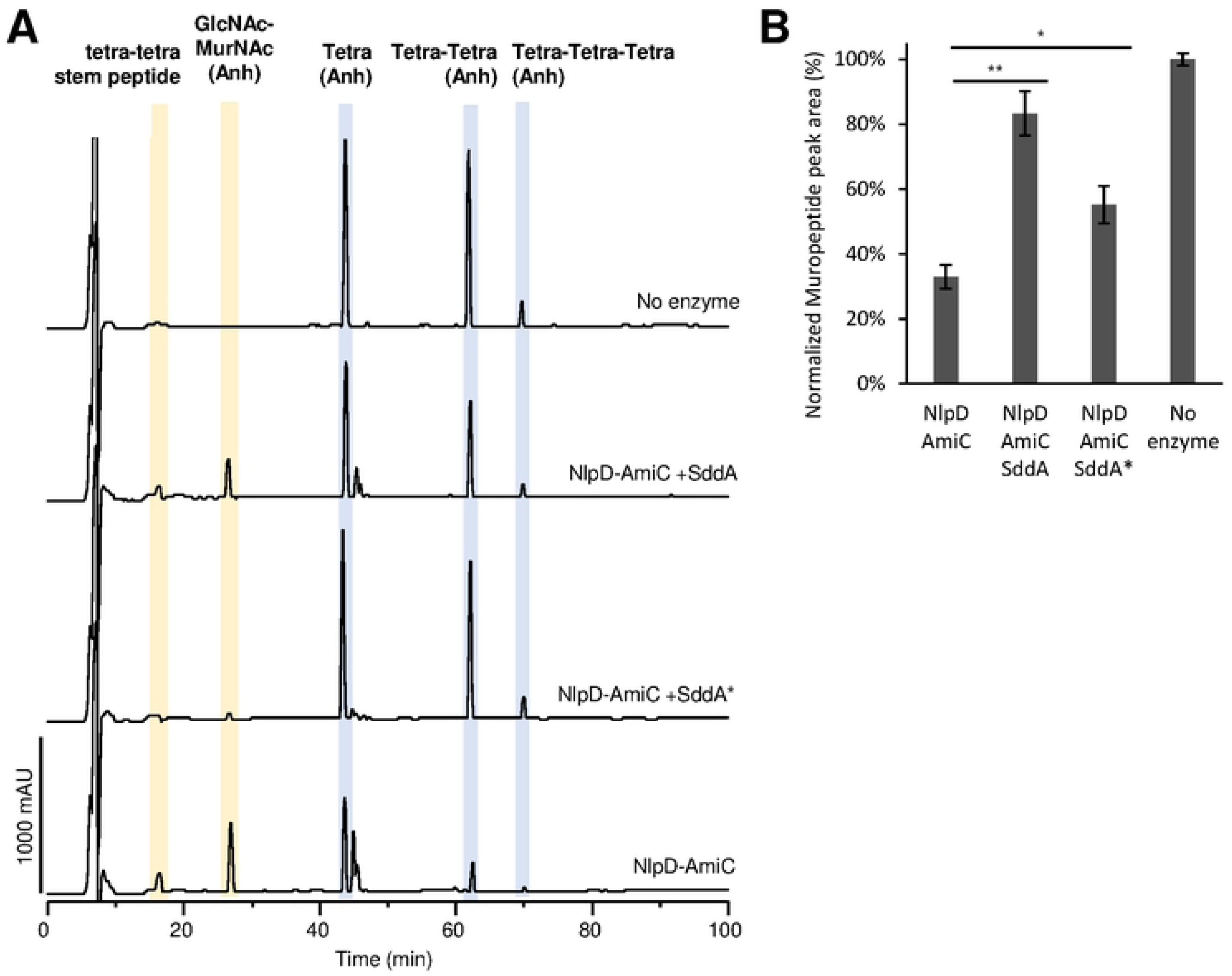
NlpD-AmiC is inhibited by SddA. **(A)** HPLC-based end point activity assay for NlpD-AmiC in the presence or absence of SddA. Sacculi were incubated with 1 µM AmiC, 1 µM NlpD and in the presence or absence of 2 µM His-SddA^24-238^ (SddA) or His-SddA^24-238^ D31A/D32A (SddA*) for 30 min at 37°C. Representative chromatograms are shown. Identity of the labelled peaks is shown above and structures are depicted on Fig S19E. **(B)** Quantification of peak areas for undigested muropeptides (peaks shaded in blue) in **(A)**. Areas are normalized to the ones in the control with no enzyme (100%). Values are average ± SD of three repeats, * p-value < 0.05 and ** p-value < 0.005.

## DISCUSSION

PgdA-type PG deacetylases have been found in many Gram-positive bacteria where they have roles in protecting the cell from exogenous PG hydrolases such as lysozyme [27, 29, 60], autolysin regulation [61], osmotic stability [62], sporulation [63] and virulence [28–30, 64]. Gram-negative bacteria have an OM that protects the PG from exogenous hydrolases, and there was no clear evidence for the presence of PgdA-like PG deacetylases. Rather, we now identified the first PG deacetylase in a Gram-negative bacterium, *E. coli* that, unlike PgdA, acts specifically on denuded glycan strands and has a role in modulating septum cleaving amidases.

SddA is the first characterized protein from the divergent polysaccharide deacetylase family (PFAM PF04748), although two X-ray crystal structures are available on the PDB database: entries 2NLY corresponding to BH1492 from *Bacillus halodurans* and 2QV5 corresponding to ATU2773 from *Agrobacterium tumefaciens*. Both structures show remarkable similarity to the polysaccharide domain of PgdA. However, only the BH1492 structure contains a Zn^2+^ ion coordinated by four residues (two histidine and two aspartic acid residues) instead of the three in PgdA (two histidine and one aspartic acid residues). These four residues are also present in *E. coli* SddA and here we show that the aspartic acid residues are required for catalytic activity. Intriguingly, Zn^2+^-coordinating residues are not present in ATU2773, but this gene still resides in the same operon as an *envC* homologue in *A. tumefaciens*.

This might indicate that a catalytically inactive ATU2773 has retained a regulatory function linked to EnvC in this organism (see below).

### Link to amidase-mediated septal PG cleavage

The cleavage of the new division septa by peptidoglycan amidases enables Gram-negative bacteria to separate the two daughter cells. The activation of these enzymes occurs from inside the cell via FtsEX-EnvC or the OM via NlpD (or ActS). Here we describe a new potential player in amidase regulation, the enzyme SddA, which we demonstrate to be a denuded glycan strand deacetylase that removes the *N*- acetyl group for Glc*N*Ac residues. In the cell, SddA localizes at the septum where denuded strands are produced during cell division by the amidases. Importantly, our data suggest that SddA induces cell chaining if its expression level exceeds that of the amidase activator EnvC, a phenotype characteristic of deficient peptidoglycan amidase activity. This provides an explanation for the conserved location of *sddA* and *envC* in the same operon in many Gram-negative bacteria. Importantly, SddA overexpression mimicked the effect of removing amidase-activators in cells deficient in the Tol-Pal system, suggesting that SddA inhibits amidases *in vivo*. This hypothesis is also supported by the *in vitro* inhibition of NlpD-AmiC by SddA. We only detected a small activation of EnvC(LytM)-AmiA but not EnvC(fl)-AmiA by SddA *in vitro*. We speculate that the effect of excessive SddA on EnvC requires the presence of FtsEX, which is ultimately responsible for EnvC activation of the amidases. More experiments will be needed to address how precisely SddA affects FtsEX-EnvC.

### Why does the cell deacetylate denuded glycan chains?

Our data are consistent with our current model in which SddA modulates a transition of amidase activation from inside the cell in early septation to amidase activation from the OM in late septation (Fig 8). We hypothesize that SddA inhibits NlpD-AmiC in early septation to allow PG hydrolysis to be solely controlled by the divisome via FtsEX-EnvC and ultimately FtsZ. In late septation, FtsZ becomes less important and even dispensable, as it leaves the division site before septation is completed, leaving PG synthesis to drive septation [65, 66]. Thus, FtsEX-EnvC might be less efficient in activating amidases at later stages of septation. In this phase, Prc might degrade SddA to allow NlpD-AmiC to be activated, shifting amidase activation control to the OM. In the absence of a phenotype of the Δ*sddA* strain, this regulation mechanism is not essential in *E. coli* but we cannot exclude that SddA is important in other species or under certain growth conditions that we have not tested. Given the widely conserved genetic link between *envC* and *sddA* (Fig 1), SddA-mediated modulation of septum cleavage could provide an advantage for Gram-negative bacteria. SddA might ensure a higher energy efficiency for the division process through confining PG hydrolysis more precisely in space and time. In support of this, the third gene of the *envC*-*sddA* operon, *gpmM*, encodes for an alternative glycolysis enzyme, establishing a link between PG hydrolysis activation and energy metabolism. More experiments will be needed to test such possible regulatory connections.

**Fig 8.**
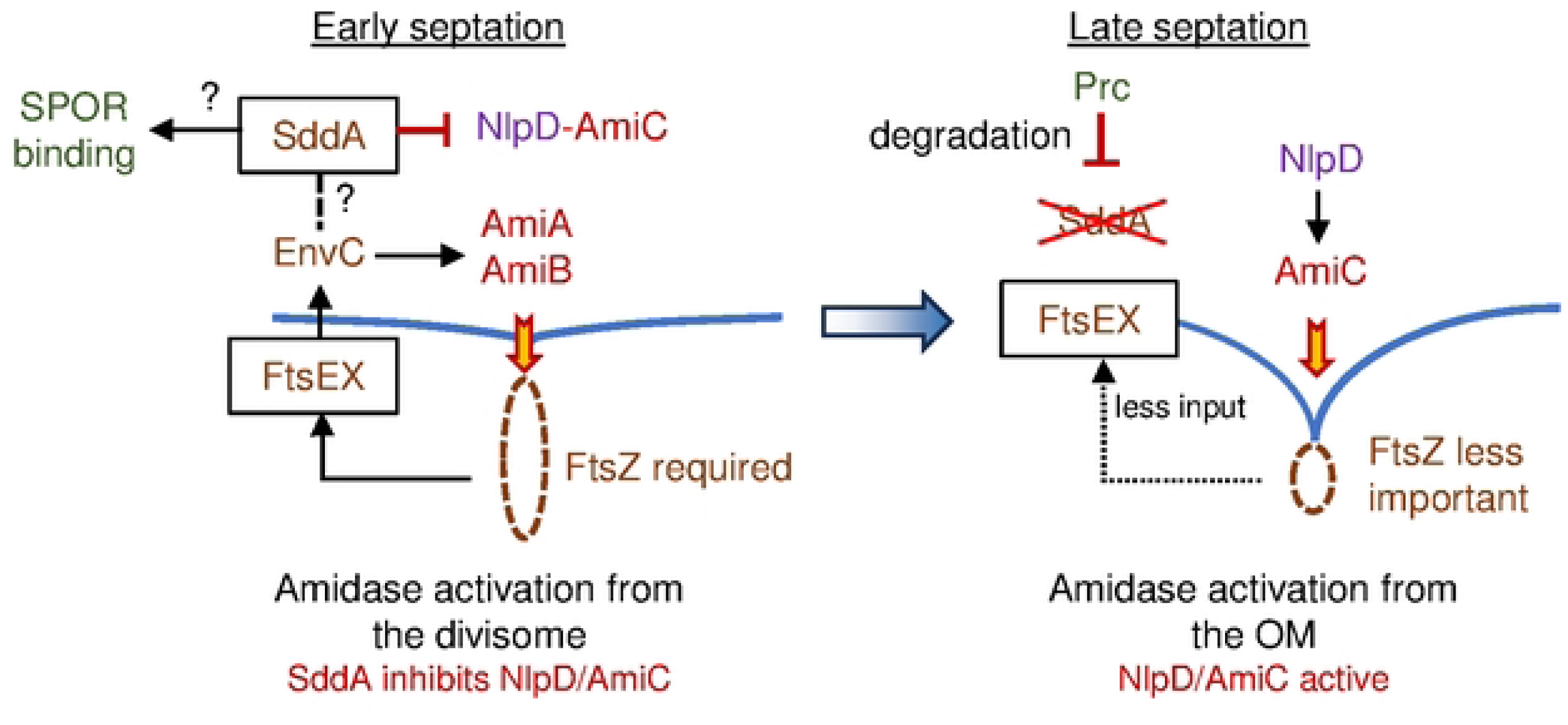
Model for the role of SddA. SddA inhibits NlpD-AmiC during early septation ensuring that PG hydrolysis is controlled by the divisome from the cytoplasmic membrane. In late septation, Prc degrades SddA to allow NlpD-AmiC to be activated and shifting amidase activation control to the OM. In addition, the deacetylation of denuded glycan strands likely modulates the binding of SPOR domain proteins and the activities of glycan strand degrading enzymes.

### SddA function beyond amidases?

SPOR domains bind Glc*N*Ac residues in denuded strands and it remains to be tested if the (partial) deacetylation of the denuded strands affects the binding and/or function of each of the SPOR proteins. Cells lacking SddA have no major growth or cell division phenotype that would be consistent with a loss of function of the essential cell division protein FtsN, hence we hypothesize that FtsN functions largely normally without SddA-mediated glycan chain deacetylation. Consistent with this view, we detected no difference in the recruitment of FtsN’s SPOR domain (iSPOR) to midcell in the absence of SddA (Fig 2E). Whether FtsN and other SPOR proteins are affected by the lack of glycan chain deacetylation under specific conditions, and whether deacetylation alters their affinities to glycan chains or their cellular dynamics, should be further tested in the future.

Although the reason is not well understood, the chaining cells of PG hydrolase gene mutants suffer from a higher permeability of their OM. Interestingly, the deletion of *sddA* in strains containing only the NlpD or EnvC amidase activators reduced the sensitivity to deoxycholate, indicating a restoration of the OM permeability barrier function. This result is consistent with a role of SddA as an inhibitor of amidase activity and/or activation. In a genetic background where amidase activation is defective, removal of an inhibitor for these enzymes is expected to improve amidase function, and hence alleviate OM defects. What it is more puzzling is the deleterious effect of the deletion of *sddA* in the Δ*envC* Δ*nlpD* background. This might suggest that SddA affects, beyond the amidases, other factors involved in cell separation, such as the lytic transglycosylase MltB or the muramidase DigH, which have been implicated in daughter cell separation in the absence of amidases [57].

In conclusion, here we discovered a new PG enzyme with an enzymatic activity that, to our knowledge, has not yet been reported, the deacetylation of denuded PG glycan strands. SddA modulates PG amidases and is widely conserved in Gram-negative bacteria. Future work will investigate the possible links of SddA function to SPOR domain proteins and other periplasmic factors.

## MATERIALS AND METHODS

### Bacterial strains and growth condition

Strains used in this study are listed in S1 Table. Bacteria were grown on LB plates or in liquid LB medium (10 g L^-1^ tryptone, 5 g L^-1^ yeast extract, 10 g L^-1^ NaCl for liquid media or 15 g L^-1^ for agar plates), or in LB-Lennox medium, liquid o agar plates (10 g L^-1^ tryptone, 5 g L^-1^ yeast extract, 5 g L^-1^ NaCl) at 37°C or 30°C when indicated. Growth of bacteria for protein expression was in LB, containing antibiotic for selection. Antibiotics were used at the following concentrations: Chloramphenicol (Cam, 20 or 25 μg mL^-1^); Kanamycin (Kan, 30 or 50 μg mL^-1^), and Ampicillin (Amp, 100 μg mL^-1^). Optical density at 578 nm (OD578) was measured with the spectrophotometer UV5 Bio (Mettler Toledo). Optical density at 600 nm (OD600) was measured with the Libra S22 spectrophotometer (Biochrom Ltd).

For cell length measuring of BW25113, Δ*sddA*, Δ*dedD* and Δ*sddA* Δ*dedD* strains (Fig 2C), strains were grown overnight in LB-Lennox at 30°C, diluted 1:100 in fresh medium and grown to an OD578 of 0.4. Cultures were diluted in fresh medium to OD578 of 0.1, and microscopy samples were taken when cultures reached OD578 of 0.3.

For SddA overexpression, BW25113 or deletion mutant derivative cells, containing either pGS100 and its derivatives, were grown overnight at 37°C, diluted 1:100 in fresh medium containing 0.5 mM IPTG and grown to stationary phase. Microscopy samples were taken at mid exponential phase (OD600 of around 0.8).

BW25113 cells containing pGS100 and derivatives were grown overnight at 37°C, diluted 1:100 in fresh medium and grown to OD578 of 0.1. Cell cultures were divided in two halves, adding to one of them aztreonam (1 µg ml^-1^). After 30 min of incubation at 37°C, overexpression of plasmid-encoded genes was induced adding 0.5 mM IPTG. Microscopy samples were taken 80 min after IPTG induction. For Western blot, samples were taken 140 min after IPTG induction.

For SddA overproduction in Δ*envC* or Δ*ftsX* cells, MPW56 derivatives were grown at 37°C in LB and MP055 ones were grown at 30°C in LB supplemented with 0.2 M sucrose. Overnight cultures were diluted (1:100, MPW56; 1:50, MP055) in fresh medium and grown to OD578 of 0.1. Overexpression of plasmid-encoded genes was induced adding 0.5 mM IPTG. Microscopy samples were taken after 140 min (MPW56) or 230 min (MP055) of incubation with IPTG.

For *tol* defective strains, overnight cell cultures grown at 37°C were diluted 1:100 (1:50, MPW55) in fresh medium and grown to OD578 of 0.1. Overexpression of plasmid encoded *sddA* (and variants) was induced by addition of 0.5 mM IPTG. After 30 min, cell cultures were divided in two halves, adding aztreonam (1 µg ml^-1^) to one of them. Microscopy samples were taken 140 min after IPTG induction

### Construction of strains and plasmids

#### Strains

MP001 (BW25113 Δ*dedD*::FRT) was obtained by removing the Kan cassette from the corresponding Keio collection single knockout mutant strain by pCP20-encoded Flp recombinase [67, 68]. MP005 (BW25113 Δ*sddA*) was obtained by conjugation [69] of recipient BW25113 carrying pACBSR with the donor MFDpir carrying pMP019. Absence of *sddA* was confirmed by PCR using the primers FwupYibQseq and Rv downYibQseq. MP006 (BW25113 Δ*dedD* Δ*sddA*) was obtained by conjugation of recipient MP001 carrying pACBSR with the donor MFDpir carrying pMP019. Absence of *sddA* was confirmed by PCR using the primers FwupYibQseq and Rv downYibQseq. MP055 (BW25113 Δ*ftsX*) was obtained by conjugation of recipient BW25113 carrying pACBSR with the donor MFDpir carrying pMP112, growing them at 30°C on LB 0.2M sucrose. Absence of *ftsX* was confirmed by PCR using two sets of primers: FwupftsX and RvdownftsX; and FwintftsX and RvintftsX. Growth defects characteristic of *ftsX* inactivation [70] were confirmed in medium with no NaCl (LB 0% NaCl), and were rescued when supplemented with sucrose (LB 0% NaCl 0.2M sucrose) or with NaCl (LB 1% NaCl). MPW54 (BW25113 Δ*tolR*::FRT Δ*nlpD::aph*) was generated by transduction of BW25113 Δ*tolR*::FRT with P1 lysate from the strain BW25113 Δ*nlpD::aph*, and absence of *nlpD* was confirmed by PCR using primers FwupnlpD and RvintnlpD. MPW55 (BW25113 Δ*tolR*::FRT Δ*envC::aph*) and MPW56 (BW25113 Δ*envC::aph*) were constructed by transduction with P1 lysate from BW25113 Δ*envC::aph* of BW25113 Δ*tolR*::FRT and BW25113 strains, respectively, and the absence of *envC* was confirmed by colony PCRS using two sets primers: FwupenvC and RvintenvC, and FwintenvC and RvdownenvC.

Other deletion strains were obtained by moving *kan*-marked alleles from the Keio *E. coli* single gene knockout library [67] by P1 phage transduction [71] or by recombineering using the pKD46 and Kan cassette amplified from pKD4 as indicated [67, 68]. Afterwards, the Kan cassette was removed by pCP20-encoded Flp recombinase to generate unmarked deletions with an FRT-site scar sequence [68]. The removal of the *kan* gene was verified by colony PCR. Strains with multiple deletions were generated by sequential P1 transduction or recombineering and *kan* cassette removal.

#### Plasmids

All template plasmids are listed on S2 Table and oligonucleotide sequences are listed on S3 Table. All mutagenesis (base changes or insertions) steps were performed using the Q5 Site-Directed Mutagenesis Kit (New England Biolabs) according to manufacturer’s instructions.

pET28a-HSddA plasmid, encoding for full length mature SddA (residues 24-319) with an N-terminal His-tag, was amplified by PCR from genomic DNA of *E. coli* BW25113 using oligonucleotides YibQ_FW_NdeI and YibQ_REV_NdeI, and cloned into pET28a(+) with the appropriate restriction enzymes. pET28a-HYibQ^24-238^ was obtained by mutagenesis using primers YibQ_N239stop_FW and YibQ_N239stop_REV, incorporating a stop codon at position 239 in pET28a-HYibQ. pET28a-HSddA^24-238^ D31A/D32A and pET28a-HSddA^24-238^ D179A were obtained by site directed mutagenesis with primers listed in S2 Table.

pVMH17 was generated by two PCR fragments: an amplified product obtained from E. coli BW25113 genomic DNA and the oligonucleotides YibQ2pBB_frag_FW and YibQ2pBB_frag_REV, and an amplified product obtained from pBB012 and the oligonucleotides YibQ2pBB_vect_FW and YibQ2pBB_vect_REV. The same volumes of each PCR fragment were mixed, heated to 98°C and cooled down to room temperature. The DNA mixed was digested with DpnI and transformed into DH5α competent cells. pVMH19 and pVMH20 were obtained by mutagenesis to insert FLAG tags at the N- or C-terminus respectively, using primers yibQ_NtFLAG_FW and yibQ_NtFLAG_REV (pVMH19) and yibQ_CtFLAG_FW and yibQ_CtFLAG_REV (pVMH20).

pGS100-*sddA* was constructed by cloning into the EcoRI-HindIII-digested pGS100 vector a PCR fragment encoding for full length SddA (residues 1-319) amplified from genomic BW25113 DNA using oligonucleotides AP907/yibQ_EcoRI_fw and AP908/yibQ_HindIII_rv.

pGS100-*envC* and pGS100-*envC*-*sddA* were constructed by cloning into the EcoRI HindIII-digested pGS100 vector a PCR fragment encoding for full length EnvC (residues 1-419) or EnvC-SddA (residues 1-419 and 1-319, respectively) amplified from genomic BW25113 DNA using oligonucleotides AP937/*envC_*EcoRI_fw and AP938/envC_HindIII_rv or AP937/*envC_*EcoRI_fw and AP908/yibQ_HindIII_rv.

pBAD24-*nlpD* was constructed by cloning into the EcoRI-HindIII-digested pBAD24 vector the PCR fragment encoding for full length NlpD (residues 1-379) amplified from genomic BW25113 DNA using oligonucleotides AP970/NlpD_EcoRI_fw and AP998/NlpD_HindIII_rev. The inserts were verified by sequencing.

pET28a-His-AmiD was generated by two PCR fragments: an amplified product obtained from pET28a and the primers FwLICHisAmiD(V) and FwLICHisAmiD(V), and an amplified product obtained from BW25113 genomic DNA and the primers FwLICHisAmiD(I) and FwLICHisAmiD(I). Same volumes of each PCR fragments were mixed, heated to 98°C and cooled down to room temperature. The DNA mixed was digested with DpnI and transformed into DH5α competent cells. pMP018 was generated by a similar procedure, using pGEC and the primers FwpGECLIC and RvpGECLIC, and an amplified product obtained from BW25113 genomic DNA and the primers FwupYibQLIC and RvdownYibQLIC. pMP107 was generated by a similar procedure, using pGEC and the primers FwpGECLIC and RvpGECLIC, and an amplified product obtained from BW25113 genomic DNA and the primers FwupFtsXLIC and RvdownFtsXLIC. pMP108 was obtained by the same procedure, using pGS100-sddA and the primers FwpGS100LIC and RvyibQLIC, and pAND101 and the primers FwyibQsfGFPLIC and RvsfGFPLIC. pMP019 was obtained by site-directed mutagenesis (SDM) over pMP018, using the primers FwYibQSDM and RvYibQSDM, and an annealing temperature of 60°C. pMP110 was obtained by site-directed mutagenesis (SDM) over pGS100-*sddA*, using the primers FwSDMyibQD179A and RvSDMyibQD179A, and an annealing temperature of 61°C. pMP112 was obtained by the same SDM procedure, using pMP107 as template and the primers FwdelftsXSDM and RvdelftsXSDM, and an annealing temperature of 62°C. pMP116 was obtained by the same SDM procedure, using pMP108 as template and the same primers used for pMP110.

### Purification of proteins

#### His-SddA variants

Versions of SddA (His-SddA, His-SddA^24-238^ or the D31A/D32A and D179A mutants) were expressed in *E. coli* BL21(DE3) from freshly transformed cells using the corresponding plasmids (S2 Table). 1L of LB containing kanamycin and auto-induction mix (0.5% glycerol, 0.05% glucose, 0.2% lactose) was inoculated with 10 mL of an overnight starting culture in LB. Cultures were incubated for 24 h at 20°C. Cells were harvested a centrifugation at 7000×g for 10 min at 4°C. Cell pellets are homogenised in lysis buffer (50 mM Tris-HCL, 1 M NaCl, 10% glycerol, pH 8.0) supplemented with EDTA-free protease inhibitor cocktail tablets (Roche), 2 mM PMSF and DNAse I. Cells were lysed by sonication on ice for 4 min at 70% power, with 15 sec pulses and 40 sec rests between pulses. The lysates were pelleted by centrifugation at 130,000×*g* for 1 h at 4°C. The supernatant was applied to 2 mL of Ni^2+^-NTA beads (Novagen) equilibrated in buffer A (25 mM Tris pH 7.5, 500 mM NaCl) supplemented with 10 mM imidazole and incubated for 1h at 4°C. Beads were washed 5 times with cold buffer A supplemented with 50 mM imidazole and the protein was eluted with 3 mL buffer A supplemented with 500 mM imidazole. The eluted protein was analysed by SDS-PAGE and the purest fractions were pooled and extensively dialysed against buffer B (25 mM Tris pH 7.5, 500 mM NaCl, 1 mM EDTA, 10% glycerol). The protein was finally concentrated using filter concentrators with a 10,000 MWCO cut-off, the concentration was measured with a BCA protein concentration kit (Thermo), aliquoted and stored at -80°C.

#### AmiD

A His-tagged version of AmiD without its signal peptide (residues 18-276) was purified using plasmid pET28a-AmiD (S2 Table). The purification protocol was adapted from reference [72]. Briefly, freshly transformed BL21(DE3) cells were used to prepare an overnight starting culture in LB with Kanamycin at 37°C. 2 L of LB with Kanamycin were inoculated with 1:100 starting culture and incubated at 37°C until OD578 reached 0.4. Protein expression was induced by addition of 1 mM IPTG for 3h at 37°C. Cells were harvested by centrifugation at 7,000×*g* for 10 min at 4°C and resuspended in 40 mL buffer A (25 mM Tris-HCl, 150 mM NaCl, pH 8.5) supplemented with EDTA-free protease inhibitor cocktail tablets (Roche), 2 mM PMSF and DNAse I. Cells were disrupted by sonication on ice for 4 min at 70% power, with 15 sec pulses and 40 sec rests between pulses, and cell debris was removed by centrifugation at 130,000×*g* for 1 h at 4°C. The supernatant was applied to 2 mL of Ni^2+^-NTA beads (Novagen) equilibrated in buffer A supplemented with 10 mM imidazole and incubated for 1h at 4°C. Beads were washed with 10 mL each of buffer A supplemented with 10, 20, 50, and 75 mM imidazole and the protein was eluted with 3 mL each of buffer A supplemented with 100, 150, 200, and 500 mM imidazole. The elution was analysed by SDS-PAGE and the best fractions were pooled and dialysed against buffer A for 2h. Then 1 unit mL^-1^ of thrombin (Novagen) was added, and dialysis against buffer A was continued overnight. Dialysis buffer was switched to buffer C (25 mM Tris-HCl pH 8.5) for 30 min and 30 additional min after a fresh buffer change. The protein was the further purified by anion exchange using a 5 mL HiTrap Q (Cytiva) column equilibrated in buffer C. The protein was eluted with a 50 mL gradient from 0 to 100% buffer D (25 mM Tris-HCl, 1 M NaCl, pH 8.5). The elution was analysed by SDS-PAGE and the best fractions were pooled. After addition of 10% glycerol, samples were aliquoted and stored at -80°C. Protein concentration was measured by UV absorbance using an extinction coefficient at 280 nm of 50,880 M^-1^cm^-1^.

#### MltA

MltA was purified from *E. coli* 122-1 bearing plasmid pMSS [73], which encodes for a soluble version of MltA without the signal peptide (residues 1-20) and the N-terminal Cys switched for Ala. The purification protocol was adapted from [73]. Briefly, 2 L of LB with Kanamycin were inoculated with 1/100 of a starting culture and incubated at 37°C until OD578 reached 0.3. Protein expression was induced by addition of 1 mM IPTG and cells were incubated for 90 min at 37°C. Cells were harvested by centrifugation at 7,000×*g* for 10 min at 4°C and resuspended in 40 mL buffer A (25 mM Na-acetate pH 5.2, 10 mM MgCl_2_) supplemented with EDTA-free protease inhibitor cocktail tablets (Roche), 2 mM PMSF and DNAse I. Cells were disrupted by sonication on ice for 4 min at 70% power, with 15 s pulses and 40 sec rests between pulses, and cell debris was removed by centrifugation at 130,000×*g* for 1 h at 4°C. The supernatant was applied to a 5 mL of HiTrap SP FF column equilibrated buffer A. The protein was eluted with a 45 mL gradient for 0 to 100% of buffer B (buffer A with 1 M NaCl). Fractions were analysed by SDS-PAGE and the best ones were pooled, concentrated, and further purified by size exclusion chromatography using a Superdex 200 16/600 pg column using buffer C (buffer A with 0.5 M NaCl). Elution was analysed by SDS-PAGE and the best fractions were pooled, concentrated, aliquoted and stored at -80C. Protein concentration was determined by UV absorbance using an extinction coefficient at 280 nm of 1.348 (g/L)^-1^.

#### Other proteins

The following *E. coli* proteins were purified following published protocols: AmiA, AmiC, NlpD, and EnvC(LytM) [59]; EnvC(fl) and ActS [21]; PBP1B S510A [74], LpoB(sol) [75].

### PG isolation and analysis

PG was isolated from *E. coli* cells and analysed by reversed-phase HPLC as described [39]. For mass-spec analysis of Glc*N*Ac-Mur*N*Ac and Glc*N*-Mur*N*Ac in *E. coli* BW25113 WT and Δ*sddA* PG, PG was digested in MS-compatible buffer (10 mM NH_4_HCO_2_ at pH 4.8) using MltA previously dialysed in the same digestion buffer. Reactions contained 15 µg of MltA per 10 µL (150 µg) of PG and were incubated for 36 h at 37°C. Samples were then boiled for 10 min, centrifuged for 15 min at 17,000×*g* and the supernatant concentrated and analysed by LC/MS.

### SddA activity assay against radiolabelled denuded glycan strands

Unlabelled lipid II and [^14^C]Glc*N*Ac-labelled lipid II were prepared as published [76, 77]. First, radiolabelled glycan strands were prepared by polymerization of ^14^C-labelled lipid II with a TPase defective mutant of PBP1B (PBP1B S510A). Reactions contained 45 µM lipid II and 0.5 µM PBP1B S510A in 50 mM Hepes pH 7.5, 150 mM NaCl, 0.05% Triton X-100, 10 mM MgCl_2_. After incubation for 90 min at 37°C, samples were boiled for 5 min and digested with NlpD + AmiC to produced denuded glycan strands. These reactions contained 2 µM AmiC, 2 µM ActS, in the same buffer as the PBP1B reactions with the addition of 1 mM ZnCl_2_, and were incubated for 2 h at 37°C and boiled for 5 min. Radiolabelled denuded strands were either incubated with His-SddA and next digested with cellosyl to break chains into smaller oligosaccharides for HPLC analysis or first digested with cellosyl producing oligosaccharides and then incubated with SddA. In the first case, strands produced from 2.25 nmol of ^14^C-labelled lipid II (15,000 dpm) were incubated with or without 10 µM His-SddA for 3 h 30 min at 37°C in the same buffer as NlpD + AmiC reactions, boiled for 5 min and then digested with 35 µg ml^-1^ cellosyl in 20 mM sodium phosphate at pH 4.8 overnight at 37°C. In the second case, the same amount of radiolabelled strands were digested with 43 µg ml^-1^ cellosyl in 20 mM sodium phosphate at pH 4.8 and then incubated with or without 10 µM His-SddA overnight at 37C in the same buffer as above. In both cases final samples were boiled for 10 min, centrifuged for 15 min at 17,000×*g* and the supernatant was reduced with sodium borohydride and analysed by HPLC as previously described [78].

### SddA activity assays against PG

For assays of SddA with PG, 10 μl (∼150 µg) of isolated PG from *E. coli* MC1061 was incubated with or without 10 µM His-SddA in the presence of 50 mM Hepes pH 7.5, 100 mM NaCl, 0.05% Triton X-100 for 16 h at 37°C and boiled for 10 min. Next, samples we digested with 88 µg ml^-1^ cellosyl in 20 mM sodium phosphate at pH 4.8 overnight at 37°C. For assays of SddA with muropeptides, 20 μl (∼150 µg) of isolated PG from *E. coli* MC1061 were first digested with 0.14 mg mL^-1^ cellosyl in 20 mM sodium phosphate at pH 4.8 overnight at 37°C. These samples were then split in two and incubated with or without 10 µM His-SddA in the same conditions as above. In both cases samples were boiled for 10 min, centrifuged for 15 min at 17,000×*g* and the supernatants were reduced with sodium borohydride and analysed by HPLC as previously described [78].

### SddA activity assays against denuded strands

#### Assays with muramidase-digested denuded strands

First, denuded strands were prepared by digesting PG with AmiC and ActS (4 µM each) in 50 mM Hepes, ∼150 mM NaCl, 0.05 % Triton, 1 mM ZnCl_2_, for 16 h at 37 °C. After boiling reactions for 10 min, denuded strands were digested with cellosyl (30 µg ml^-1^) in 20 mM sodium phosphate at pH 4.8 for 16 h at 37 °C. After boiling reactions for 10 min, the cellosyl-digested denuded strands were incubated with or without 10 µM His-SddA in the same buffer as the amidases for 24 h at 37 °C. Final reactions were boiled again for 10 min, centrifuged for 15 min at 17,000 *g* and the supernatant was analysed by HPLC without reduction (see below).

#### Assays with denuded strands followed by lytic transglycosylase digestion

Denuded strands were prepared by digesting *E. coli* MC1061 or BW25113Δ6LDT PG with AmiD. Digestions were performed with 4 µM AmiD in 20 mM Hepes pH 7.5, 0.5 mM ZnCl_2_, incubating for 16 h at 37°C. 0.2 nmol of enzyme were added per 10 µL (∼150 µg) of purified PG. Samples were boiled for at least 5 min before the next step. SddA reactions included 5 µM of purified SddA and 10 µL of digested PG (50 µL for AmiD reactions) in a total volume of 100 µL. Buffer was 20 mM Hepes pH 7.5, ∼150 mM NaCl and 0.25 mM ZnCl_2_. Reactions were incubated at 37°C for 24 h and stopped by boiling for 5 min. For analysis, samples were digested with MltA to produce anhydro-disaccharide units. 4 µM MltA and 20 mM sodium phosphate buffer at pH 4.8 were added to SddA reactions, bringing total volume to 150 µL, and incubated for 16 h at 37°C, boiled for 5 min, centrifuged for 15 min at 17,000×*g* and the supernatant was analysed by HPLC (see below).

#### HPLC analysis of SddA reaction products

HPLC analysis was done using a Prontosil 120-3-C18 AQ reversed-phase column (250 × 4.6 mm, 3 μm particle size) and a HPLC apparatus with binary pump system, UV detector and automatic fraction collector. The column temperature was set at 55°C. A 3 h gradient from 0 to 15% acetonitrile in 0.1% formic acid was use for elution. When indicated peaks were isolated by automatic fraction collection and analysed by mass spectroscopy.

### Amidase activity assays

Assays contained 10 μl (∼150 µg) of isolated PG from *E. coli* MC1061 or *E. coli* BW25113Δ6LDT as indicated. Reactions NlpD-AmiC assays contained 1 µM of each protein, while for EnvC-AmiA and ActS- AmiC reactions contained 2 µM of each protein. SddA was added as twice the concentration of the amidase and activator. Reaction buffer was 20 mM Hepes pH 7.5, 100 mM NaCl, 0.1 mM ZnCl_2_, 0.05% Triton X-100. Reactions were performed in a 100 µL volume and were incubated at 37°C for 30 min (NlpD-AmiC), 2 h (EnvC-AmiA) or 1 h (NlpD-AmiC). Reactions were stopped by boiling for 5 min. For analysis, samples were digested with MltA to produce anhydro-disaccharide units and analysed by HPLC as described above for SddA assays. At least two repeats were performed per condition. The areas of muropeptide peaks still present in reactions were quantified and analysed relative to the same peaks in control reactions with no amidases.

### Esterase assay with 4-nitrophenyl acetate

4-nitrophenyl acetate (Sigma) were freshly dissolved in DMSO. Reactions had a total volume of 100 µL and contained 10 µM His-SddA or His-SddA^24-238^ and 10 mM 4-nitrophenyl acetate in 25 mM Tris pH 7.5, 150 mM NaCl, 10% DMSO. Reactions were carried out in 96-plate wells and monitored using an Infinite F50 (Tecan) plate reader for 30 min at room temperature.

### Western blot and immunodetection of GFP fusion proteins

Indicated cell cultures (10 ml) were centrifuged at 4°C (3,200×*g*, 10 min) and the pellet was resuspended in 300 μl PBS sterile. Cells were disrupted by sonication (30 s, cycle 0.6, 60% amplitude) using a Sartorius LABSONIC M. Protein extract concentration was determined using Pierce BCA Protein Assay Kit (ThermoScientific). After addition of SDS loading buffer, the protein extracts were boiled at 95°C for 10 min. Fifteen µg of each sample were loaded on 12% polyacrylamide gels and analysed by SDS-PAGE followed by Western blot and immunodetection. Anti-GFP antibody (1:10,000) (A-6455, Invitrogen) and anti-rabbit HRP-IgG (1:5,000) (A8275, Sigma–Aldrich) were used as primary and secondary antibody, respectively. Western Blots were developed using ECL Prime Western Blotting System (GE Healthcare) and processed with Amersham™ Imager 680.

### Microscopy

#### Figures S8 and S10

Cells were grown to the indicated optical densities and in the indicated media and conditions. When indicated, 500 µL samples were taken and fixed by addition of 8% paraformaldehyde and 0.01% glutaraldehyde and incubation for 15 min at room temperature and 30 min on ice. Cells were then washed twice with 1 mL phosphate buffer saline (PBS) at 4°C and stored in PBS at 4°C until use.

Fixed cells were spotted on 1% agarose pads containing 3 µg ml^-1^ FM5-95 membrane dye (ThermoFisher) and deposited on microscope slides and sealed with a glass coverslip (VWR or Fisher Scientific). Slides were visualized using a Nikon Eclipse Ti equipped with a Nikon Plan Apochromat 100x oil objective, a Cool LED pE-4000 light source, a Photometrics BSI camera and NIS-Elements software. For FM5-95 fluorescence, a Chroma 49008 filter-set was used (EX560/40, DM585lprx, EM630/75).

#### Figures 2C, 5, 6, S12, S15, S17

1.5 ml of indicated samples were centrifuged (16,000x*g*, 2 min) and the pellet was resuspended in 1 mL sterile PBS. After another centrifugation, cell pellets were resuspended in 12.5 μl of PBS and 12.5 μl of 3% paraformaldehyde (diluted in PBS) for cell fixation and kept at 4°C until microscopy analysis. When indicated 3 µg ml^-1^ FM5-95 dye was added to the agarose pads. Cells samples were analysed in the Advanced Light Microscopy Facility at the Centro de Biología Molecular Severo Ochoa (Madrid, Spain). Images were acquired on a Zeiss Axiovert 200M widefield inverted system. A Zeiss EC Plan-Neofluar 100x 1.3 NA Ph3 oil immersion objective was used with Zeiss Inmersol 518F (n = 1.518) oil. Metamorph 7.10.5.476 was used for image acquisition, saving data as .tif formatted files. Fluorescence was sequentially excited with a Spectra-X Lumencor, with filters 470/24 and 575/25 for excitation of sfGFP fusions and FM5-95, respectively. Emission was collected on a sCMOS PCO edge 4.2 bi camera with 16-bit depth, 0.003969 µm^2^ pixel size and at 1024×1024 pixel resolution (applying a 2×2 binning). sfGFP and FM5-95 emission signals were collected through a 514/32 and 595-34 emission filters, respectively. For Fig 2C, a Zeiss Plan-Apochromat 63×1.4 Oil DIC M27 immersion objective was used with Zeiss Inmersol 518F (n = 1.518) oil. Metamorph 7.10.5.476 was used for image acquisition, saving data as .tif formatted files. Fluorescence was sequentially excited with a Spectra-X Lumencor, with filter 575/25 for excitation of FM5-95. Emission was collected on a sCMOS PCO edge 4.2 bi camera with 16-bit depth, 0.003969 µm2 pixel size and at 2048×2048 pixel resolution (applying a 1×1 binning). FM5-95 emission signals were collected through a 595-34 emission filter.

Images were analysed by segmentation with Omnipose [79] and measurement of cell parameters (length along the cell axis and averaged width along the cell axis) using custom-made scripts. Demographs for SddA(D179A)-sfGFP localization were built by plotting the normalized fluorescence intensity along the cell axis using customs scripts. Each intensity profile along the cells axis was calculated by addition of the fluorescence intensity of 5 pixels perpendicular to each position along the cell axis (*I*_*i*_). The normalized intensity (*I_i_^norm^*), and intensity profiles were normalized by:

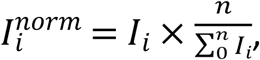

where *n* indicates the length of the profile. For Fig 2C, the violin plots, the average lengths and standard deviations were obtained by analysis of the data with CellDataAnalyser.

#### Figures 4, S11, S13

Cells were grown in LB-Lennox containing the opportune antibiotics and 0.5 M IPTG. When indicated, a total amount of cells equal to 3 OD were sampled and fixed in 2.8% formaldehyde and 0.04% glutaraldehyde. Cells were incubated for 30 min at 37°C, with shaking, washed with PBS, and resuspended in 0.5 ml of PBS stored in PBS at 4°C until use [80]. Fixed cells were stained with 5 µg ml^-1^ FM5-95 membrane dye (ThermoFisher) and 1 µg ml^-1^ DAPI DNA dye (Sigma), deposited on microscope slides and sealed with a glass coverslip (VWR or Fisher Scientific). Slides were visualized using a Zeiss LSM 900 equipped with C Plan-Apochromat63x/1.4 Oil DIC (WD=0.14mm), PALM MicroLaser Systems, AxioCAM ICc1 camera and PALM ROBO-software.

#### Figure 2D-E

BW25113 and BW25113 Δ*sddA*::FRT were transformed with pCH-ss*^dsbA^*-sfGFP-iSPOR (pAP674) to create strains AP698 and AP699, respectively. This plasmid is a derivative from pJL427 encodes the isolated SPOR domain of FtsN, tagged with sfGFP in its N-terminus, and with the signal peptide sequence from *dsbA* for periplasmic export, under control of an IPTG-inducible promoter [81]. Cells were grown in EZ rich defined medium (EZRDM) [82] containing 150 µg mL^-1^ chloramphenicol (EZRDM+Chlor) at 37°C overnight to saturation. In the morning 1 μL of culture was diluted into 3 mL of EZRDM+Chlor and 100 μM IPTG to induce expression of sfGFP-iSPOR. Cells were grown to an OD600 ∼0.25-0.40 during which time 500 μL cells were obtained and fixed using paraformaldehyde (2.8%) and glutaraldehyde (0.04%) in EZRDM for 15 min followed by washing three times in PBS by repetitive centrifugation (11,000×*g* for 5 min). Cells were then imaged on a 3% PBS agarose pad using a 100× 1.49 NA oil-immersion objective (Olympus). The light was focused onto the chip of an EMCCD camera (iXon Ultra 897, Andor Technology) with a final pixel size of 100 × 100 nm. To obtain the midcell intensity we used a 3 pixel line width to measure the intensity at the midcell (septa) and subtracted the intensity at the quarter position (background). The median midcell intensity of the WT sample (AP698) for each biological replicate was then used to normalize the midcell intensities values of AP698 and AP699 resulting in “relative midcell intensity”.

### Analysis of SddA conservation

We used the AnnoTree database [44] to study the conservation of SddA homologues within bacteria and the genetic link between EnvC and DpaA. This database contains a set of archaeal and bacterial genomes which are completely sequenced and consistently annotated with PFAM, KEGG and TIGRFAM annotations [44]. To find SddA homologues, we searched for genes annotated with either PFAM number PF04748 (divergent polysaccharide deacetylase) or KEGG number K09798 (YibQ-like, uncharacterized protein), obtaining 7174 sequences in total. EnvC analogues were identified as genes annotated with KEGG number K22719 (EnvC; murein hydrolase activator) that were not also annotated as any of the KEGG numbers for other LytM activators in *E. coli* (K19304, K06194 or K12943 for MepM, NlpD or YgeR), obtaining 17185 sequences. GpmM homologues were identified as genes annotated with TIGRFAM number TIGR01307 (phosphoglycerate mutase (2,3-diphosphoglycerate- independent)), obtaining 17651 sequences. Finally, AmiC homologues were identified as genes annotated with either KEGG number K01448 (AmiABC; *N*-acetylmuramoyl-L-alanine amidase) or PFAM number PF01520 (N-acetylmuramoyl-L-alanine amidase) that also had a PF11741 (AMIN domain) annotation, obtaining 10499 sequences. The number of proteins of each type per genome was counted using custom Python scripts, and the generated data sets were represented along a phylogenetic tree of all genomes in AnnoTree using iTOL [83]. The localization within the genome of each of the identified sequence was obtained from the downloaded Annotree SQL database and analysed with custom scripts. For each pair of homologues analysed (EnvC+SddA, EnvC+AmiC or SddA+AmiC), genomes containing both homologues were identified and the pair of genes closest together within the same chromosome was selected, counting the number of times the closest pair was on the same strand and the distance in base-pairs between them. The scripts to perform this analysis are available upon request.

### Protein structure and Protein-Protein Interaction Modeling

Protein structure models and protein-protein interaction models were generated using a local installation of AlphaFold (AlphaFold v2.3.1) [84, 85]. Amino acid sequences of the proteins of interest were submitted in FASTA format, following the server’s input guidelines. AlphaFold utilizes a deep neural network architecture trained on structural data from the Protein Data Bank (PDB) [86] and multiple sequence alignments to predict protein structures, including protein-protein interactions. For each protein pair, five models were generated, and the highest-confidence model, based on AlphaFold’s predicted *ranking_score* which is calculated from the predicted template modeling (pTM) score and the interface predicted template modeling (ipTM) score among other metrics.

The Predicted Aligned Error (PAE) matrix provides a visual representation of the predicted positional accuracy between residues in the modeled protein complex. Each cell in the matrix represents the estimated alignment error (in Å) between pairs of residues across the protein structure, as predicted by AlphaFold. The matrix is color-coded, where darker shades indicate lower predicted alignment errors and higher model confidence, and lighter shades denote areas of greater uncertainty in alignment.

In our analysis, the PAE matrix was used to assess the reliability of inter-protein and intra-protein contacts within the predicted complex. Low error regions, typically appearing along the diagonal of the matrix, correspond to structurally stable segments of each protein. In contrast, off-diagonal regions with low PAE values are indicative of high-confidence inter-protein interactions, suggesting specific interaction sites or binding interfaces. Regions with high PAE values highlight areas of structural flexibility or uncertainty, which may correspond to disordered regions or low-confidence interactions.

The prediction of the Prc-SddA complex was performed using the AlphaFold3 server (https://golgi.sandbox.google.com/, accessed on October 22^nd^, 2024) [87]. All five models generated show an interaction between Prc and the C-terminal region of SddA. The best model (pTM=0.66) is shown in S7D Fig. The analysis of this prediction and the image generation was performed using UCSF ChimeraX [88].

### Molecular Dynamics Simulations

Molecular dynamics (MD) simulations were performed for the wild-type SddA protein (WT) and its D179A mutant using five independent replicates for each system. Modeled structures were obtained using AlphaFold. The protonation states of ionizable residues were determined prior to molecular dynamics simulations using the H++ server [89] at pH 7.4. This tool computes residue protonation based on pKa predictions, considering the influence of the surrounding environment on each residue’s ionizable groups. The calculated protonation states were subsequently used to ensure an accurate representation of the protein’s charge distribution under the simulation conditions. The system was prepared using the *tleap* module of AmberTools [90]. The protein structure was parameterized using the ff19SB force field [91], and the system was solvated in a triclinic box of OPC water [92] molecules, ensuring a minimum distance of 10 Å between any protein atom and the edge of the box. Sodium and chloride ions were added at random positions to achieve a neutral system and an ionic concentration of 0.15 M. To enable the use of a 4 fs integration timestep during the production phase, the hydrogen mass repartitioning (HMR) [93] technique was applied using the *parmed* module of AmberTools [94]. This method redistributes part of the hydrogen atom masses to their bonded heavy atoms, effectively increasing the hydrogen mass and allowing for a longer integration timestep without compromising simulation stability. The modified topology files were used in all subsequent simulations.

The MD protocol consisted of four phases: energy minimization, heating, equilibration, and production. (i) **Energy Minimization:** This was done in three stages. First, only water molecules were minimized; second, water and ions (excluding hydrogens) were minimized; and third, the full system was minimized. Steepest descent was used for the initial 500-1,000 steps, followed by conjugate gradient minimization. Periodic boundary conditions with an isotropic barostat were applied throughout. (ii) **Heating:** The system’s temperature was gradually increased from 100 K to 300 K over 200 ps, using a timestep of 1 fs. A Langevin thermostat [17, 95] with a collision frequency of 10 ps^−1^ controlled the temperature. The simulation was at constant volume, with harmonic restraints applied to the backbone and beta-carbon atoms to stabilize the solute. (iii) **Equilibration:** This phase involved 10 steps of 100 ps each using a time step of 1 fs, under constant pressure and temperature. The temperature was kept at 300 K using a Langevin thermostat, and pressure was controlled by a Monte Carlo barostat [96]. A harmonic restraint was applied to alpha-carbon atoms with decreasing force constants. Nonbonded interactions were treated with a 9 Å cutoff, and electrostatics were handled using the particle mesh Ewald (PME) method [97]. (iv) **Production**: The simulation ran for 60 ns with a 4 fs timestep. The temperature was maintained at 300 K using a Langevin thermostat, and pressure was controlled by a Monte Carlo barostat. Electrostatics were treated with PME, and nonbonded interactions had an 11 Å cutoff. Hydrogen mass repartitioning (HMR) enabled the 4 fs timestep. Center-of-mass translation and rotation were removed every 25 ps, and energy, trajectory, and restart files were recorded at 25 ps intervals to ensure accurate sampling during the production phase.

Trajectory analysis was performed using **cpptraj** [98], from AmberTools suite of programs, focusing on root mean square deviation (RMSD) across the simulation and per-residue root mean square fluctuation (RMSF). Data were visualized using R to evaluate the structural stability and flexibility of both systems.

## ACKNOWLDEDGEMENTS

This work received funding from the UK Biotechnology and Biological Sciences Research Council (BB/W005557/1; BB/W013630/1; to W.V.), from the National Institutes of Health (NIH R35 GM136436 to J.X. and NIH F32 GM150262 to A.J.P.), from MICIU/AEI/https://doi.org/10.13039/501100011033 and ERDF/EU (PID2022-140818OA-I00; to M.P.), the Comunidad de Madrid Atracción de Talento M1 program (2020-T1/BMD-19970, to M.P.). A.B. acknowledges a PhD fellowship from Fundación Ramón Areces. D.B. acknowledges a PhD fellowship (PRE2022-101502) associated to the “Centre of Excellence Severo Ochoa” grant to the CBM (CEX2021-001154-S) funded by MICIU/AEI /https://doi.org/10.13039/501100011033. CBM receives institutional grants from Fundación Ramón Areces. B.I.I. is supported by a grant from the French National Research Agency (ANR) through the PPR Antibioresistance program (ANR-20-PAMR-0010).

## SUPPLEMENTARY INFORMATION CAPTIONS

**S1 Fig.**
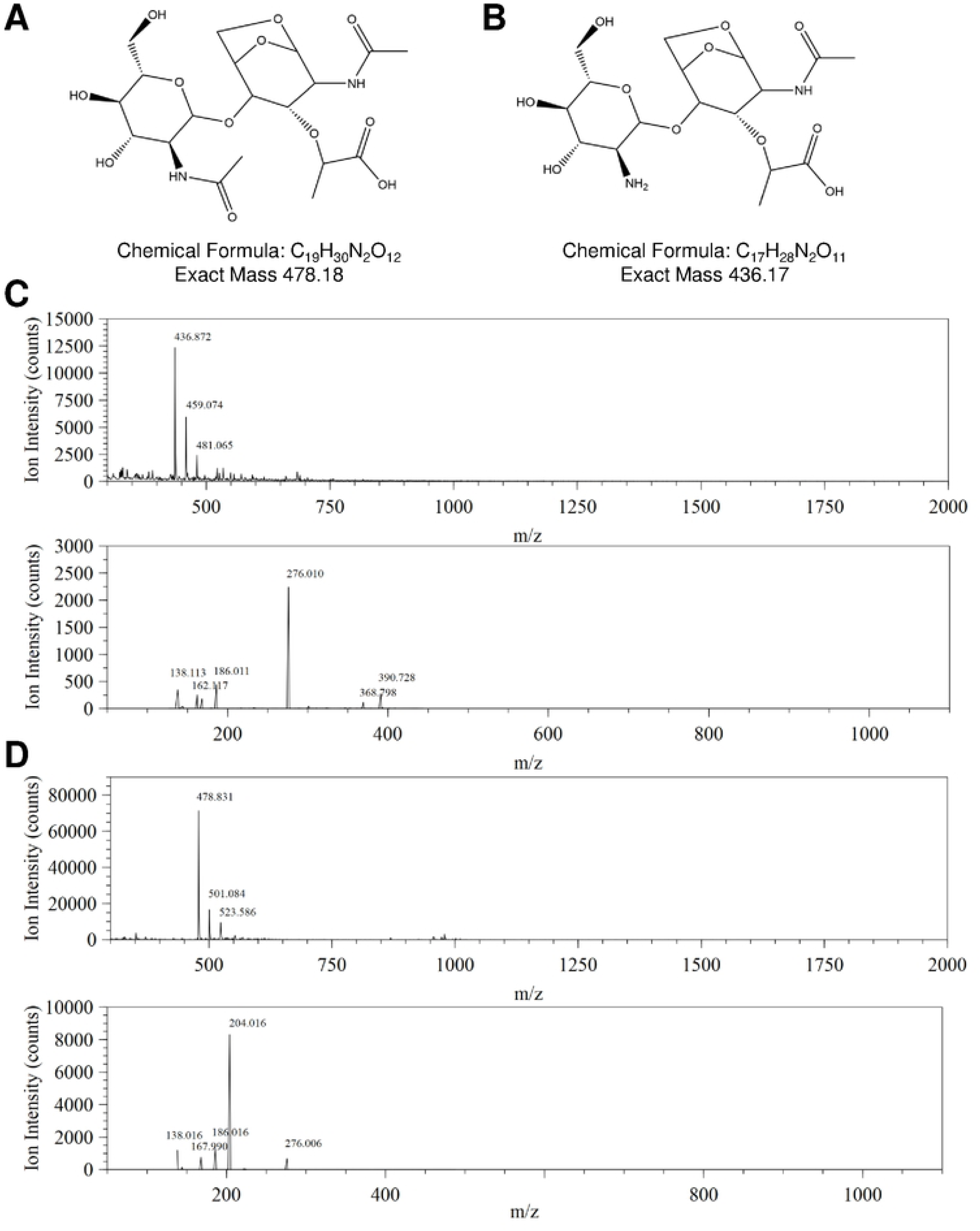
Verification by MS/MS of the identity of peaks for Glc*N*Ac-Mur*N*AcAnh and Glc*N*Ac- Mur*N*AcAnh in the MltA-digested PG samples. (A, **B)** Chemical structures of Glc*N*Ac-Mur*N*AcAnh and Glc*N*-Mur*N*AcAnh, respectively. **(C)** Verification using MS/MS (bottom) that the peak with m/z 437 (top) corresponds to Glc*N*-Mur*N*AcAnh. Fragmentation of Glc*N*-Mur*N*AcAnh yields Glc*N*(-H_2_O)+H^+^ (m/z 162), and not Glc*N*Ac(-H_2_O)+H^+^ (m/z 204), and Mur*N*AcAnh+H^+^ (m/z 276). **(D)** Verification using MS/MS (bottom) that peak with m/z 479 (top) corresponds to Glc*N*Ac-Mur*N*AcAnh. Fragmentation of Glc*N*Ac-Mur*N*AcAnh yields the Glc*N*Ac(-H_2_O)+H^+^ ion (m/z 204) and MurNAcAnh+H^+^ (m/z 276).

**S2 Fig.**
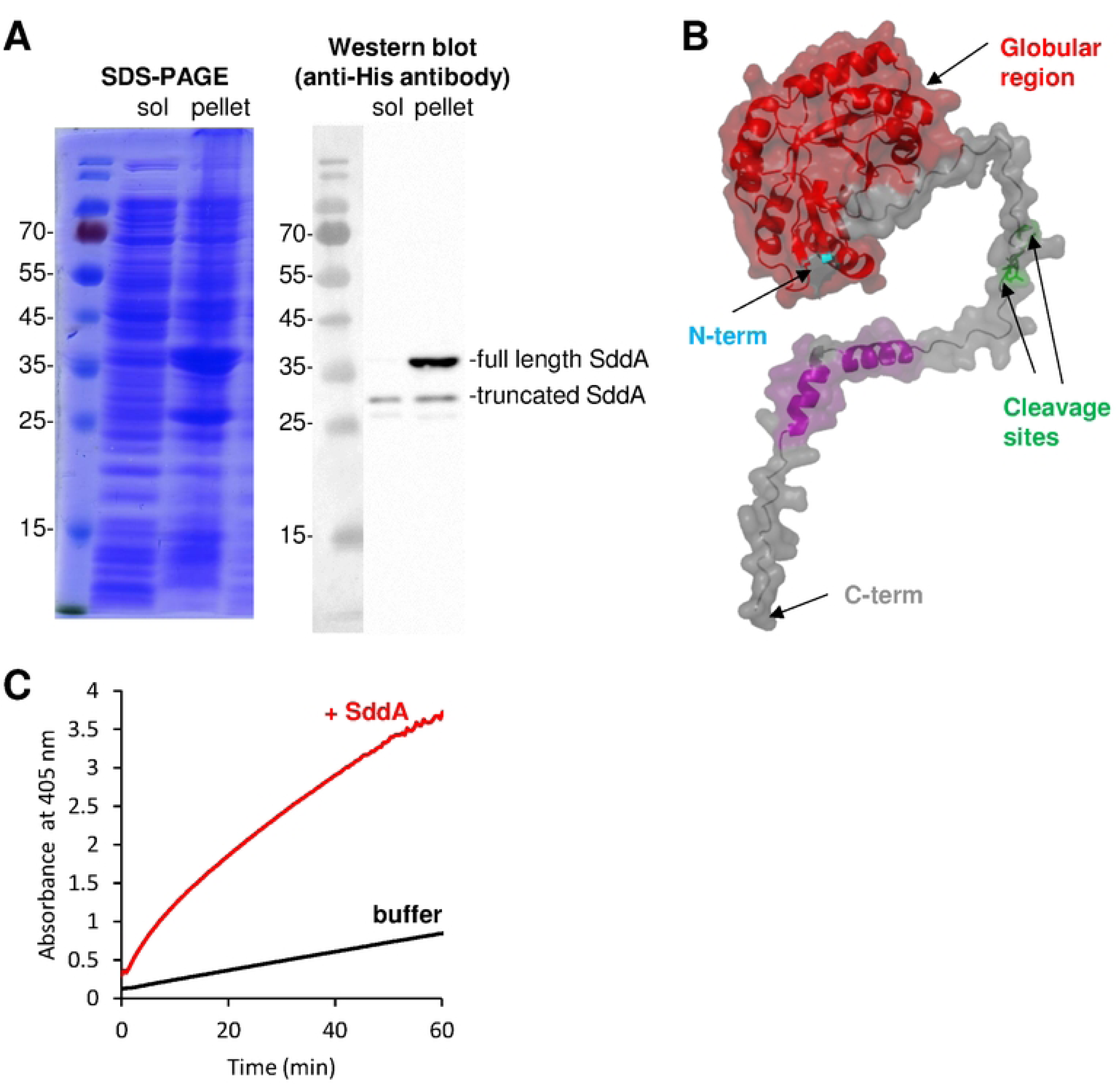
Overexpression of full length SddA in the cytoplasm produces a truncated protein. **(A)** SDS-PAGE and Western-Blot analysis of soluble (*sol*) and *pellet* fractions from sonicated cell extracts of BL21(DE3) pET28a-HisSddA induced with IPTG. The His-tag in the N-terminus His-SddA was detected using anti-His-tag antibody. **(B)** AlphaFold model of *E. coli* SddA showing the predicted globular (red) and unfolded C-terminal (grey) regions, plus the location of cleavage sites in His-SddA purified from soluble extracts as the one analysed in A. The identified cleavage sites are P258, V262, K263 and L264. **(C)** Representative 4-nitrophenyl acetate (4NA) esterase assays with His-SddA. Reactions contained 10 mM 4NA in the presence or absence of 8.8 µM His-SddA (red and black lines, respectively) and were incubated at room temperature.

**S3 Fig.**
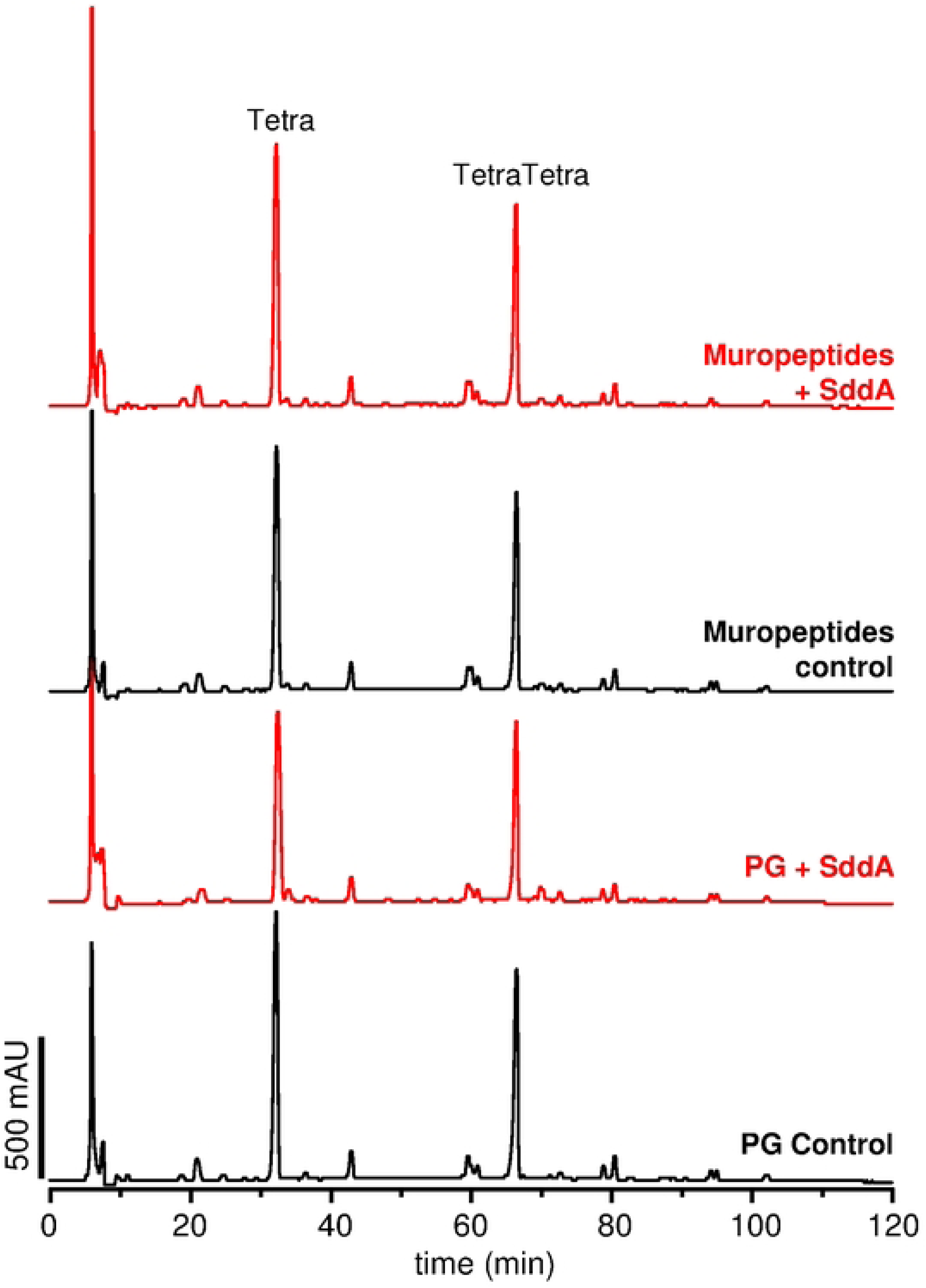
SddA is unable to digest peptidoglycan or muropeptides. PG was incubated first with SddA or buffer and then with the muramidase cellosyl (chromatograms labelled “PG”) or first with cellosyl and then with SddA or buffer (chromatograms labelled “muropeptides”), to test the activity of SddA on PG or muropeptides, respectively. SddA was added at 10 µM in both cases. SddA treatment did not introduce any changes in the resulting chromatograms.

**S4 Fig.**
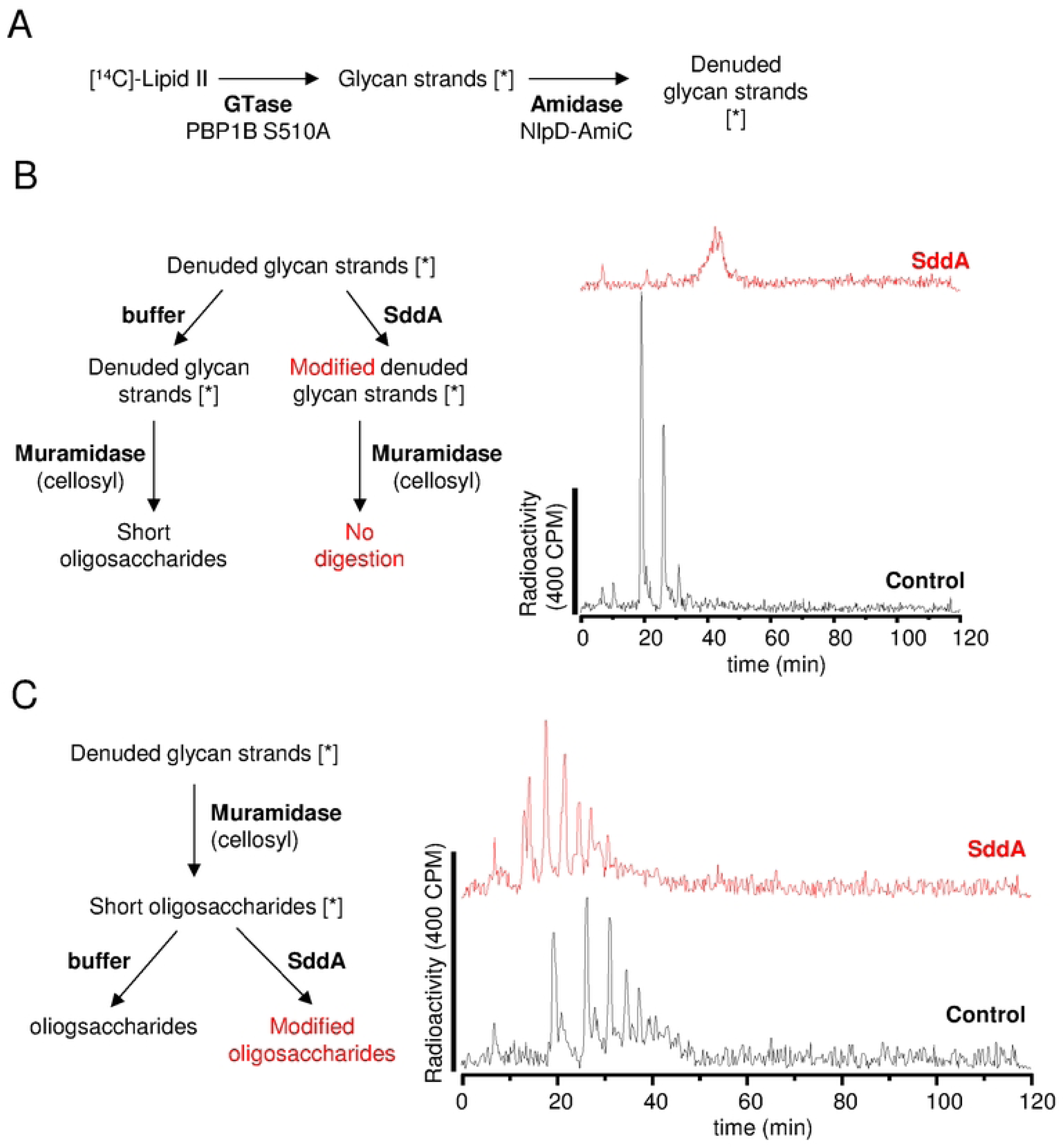
SddA can modify denuded glycan strands. **(A)** Radiolabelled denuded glycan strands were prepared from radiolabelled lipid II ([^14^C]-lipid II) using transpeptidase-defective PBP1B (S510A), and the amidase AmiC plus its activator NlpD. **(B)** Scheme depicting the preparation of samples (left side) and their analysis (chromatograms on the right side). Muramidase was unable to digest the denuded glycan strands treated with SddA. **(C)** Scheme depicting the preparation of samples (left side) and their analysis (chromatograms on the right side). SddA modified the short oligosaccharides obtained by digesting radiolabelled denuded glycan strands with a muramidase.

**S5 Fig.**
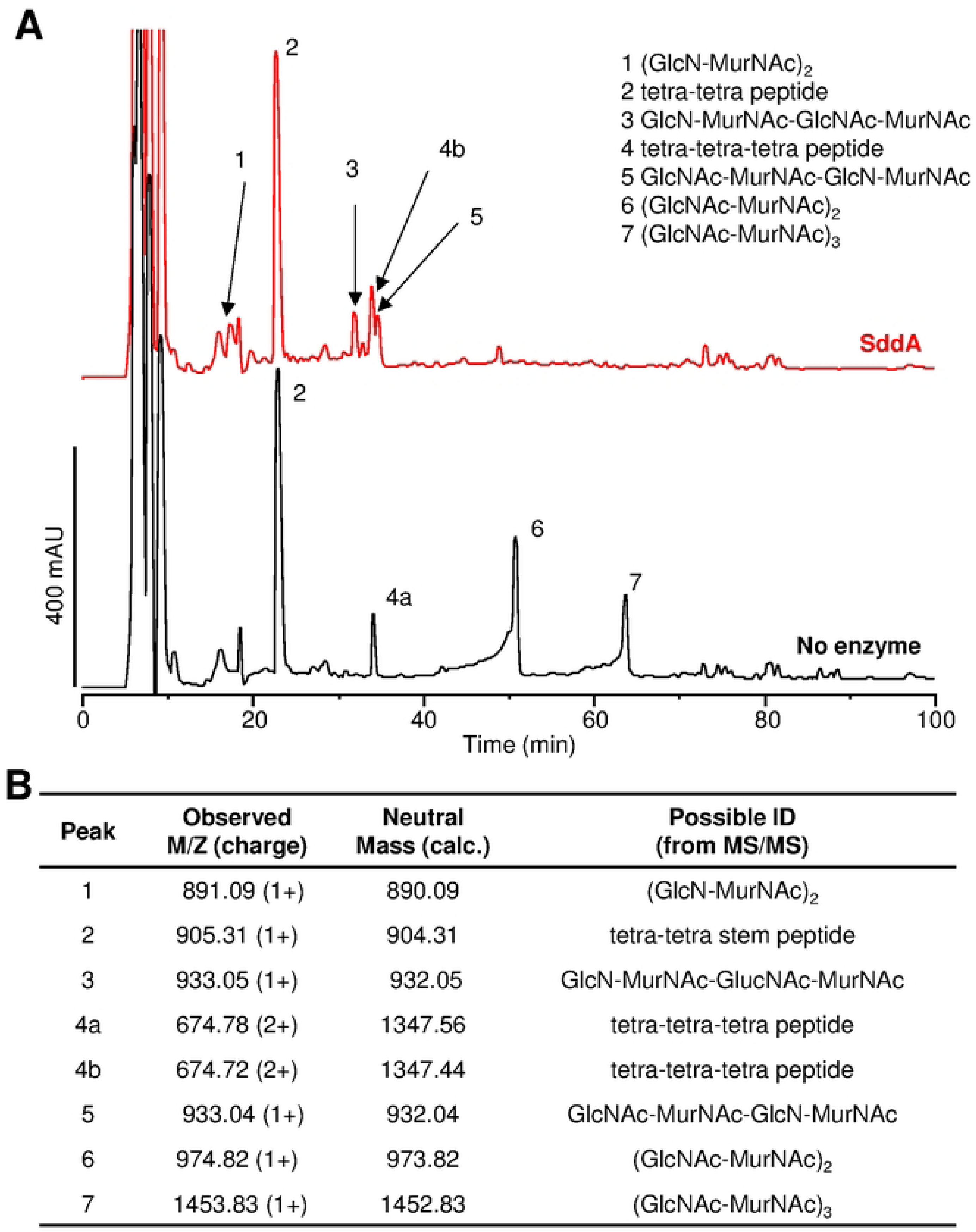
SddA deacetylates Glc*N*Ac residues in denuded glycan strands. **(A)** Chromatograms of muramidase-digested denuded glycan strands, treated with SddA (red) or buffer (black) after digestion with the muramidase cellosyl. **(B)** Results of MS and MS/MS analysis of the labelled peaks in A. All muropeptide peaks correspond to the non-reduced species.

**S6 Fig.**
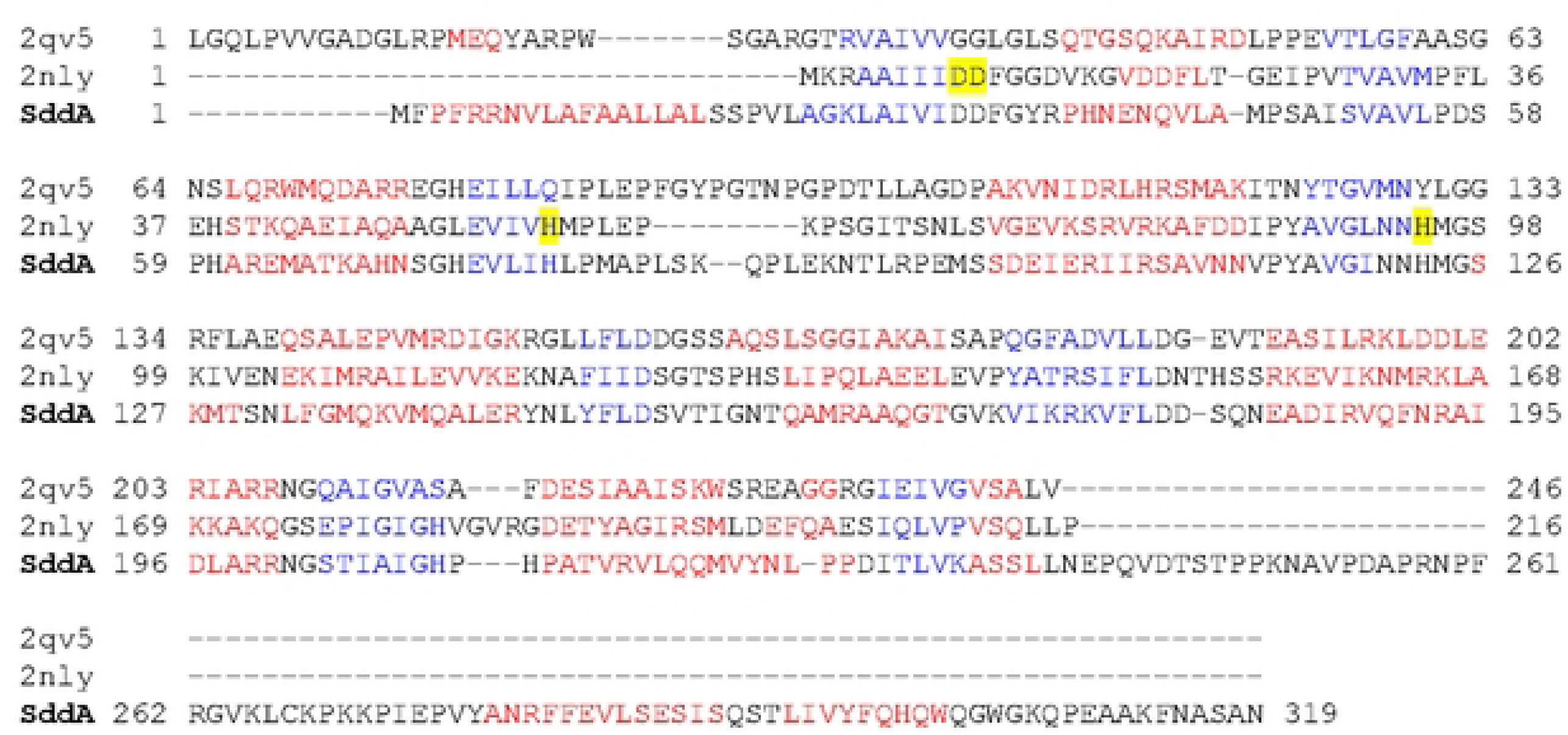
*E. coli* SddA contains conserved residues for Zn^2+^ binding. Sequence alignment of *E. coli* SddA with the sequences of the two proteins with a divergent polysaccharide deacetylase domain (PF04748) whose crystal structure is available. These two proteins are BH1492 from *Bacillus halodurans* (PDB 2NLY), which contains a Zn^2+^ in the crystal structure, and ATU2773 from *Agrobacterium tumefaciens* (PDB 2QV5), which does not contain a Zn^2+^ ion in the crystal structure. The sequences are coloured by secondary structure (red indicating alpha helix and blue beta strand). The residues coordinating Zn^2+^ in BH1492 are highlighted in yellow.

**S7 Fig.**
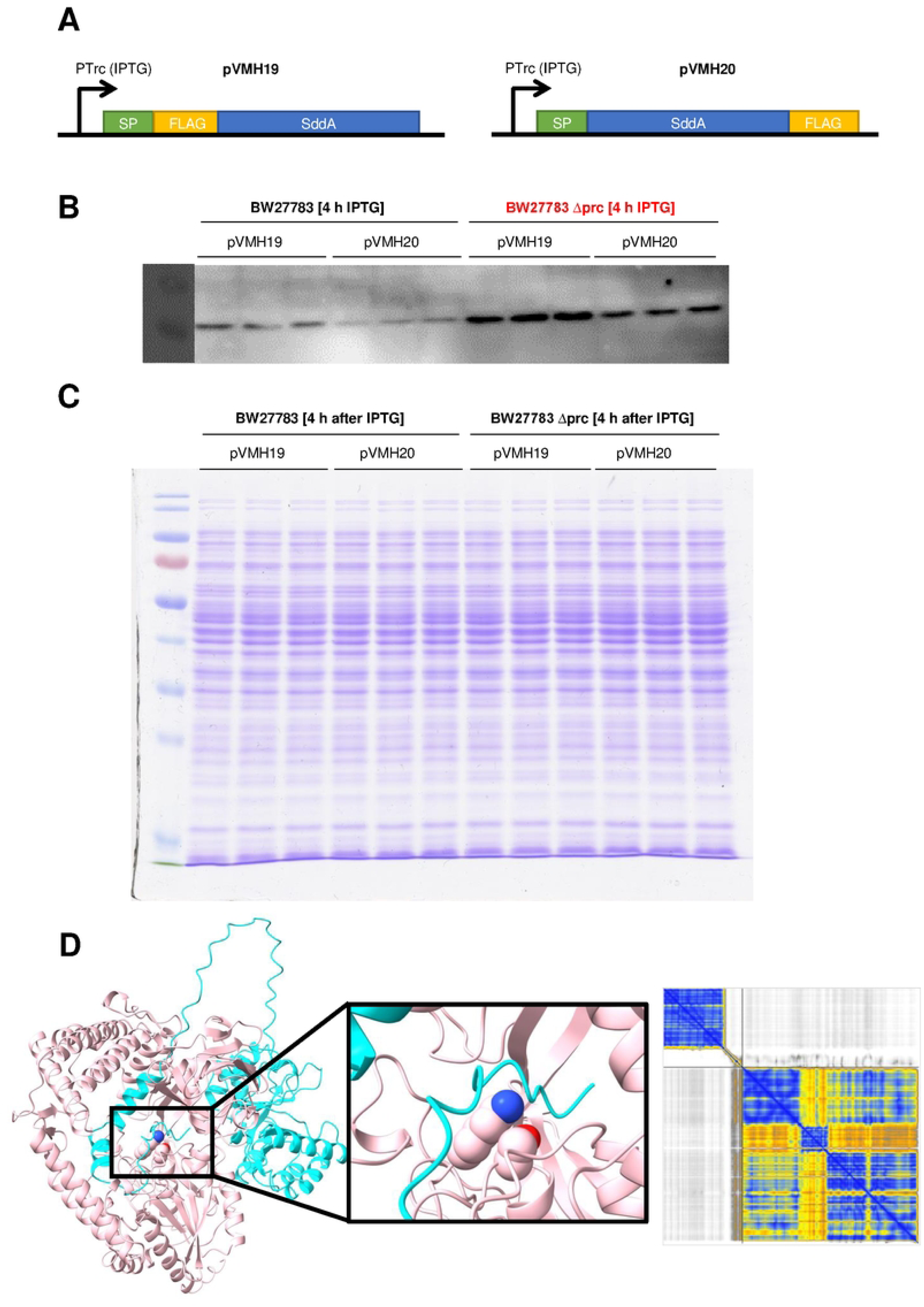
Protein level of ectopically expressed SddA is higher in a Δ*prc* background. **(A)** Scheme depicting the cloning of *sddA* with a FLAG tag and its native signal peptide (SP) into pVMH19 and pVMH20 plasmids. (**B)** Expression of SddA from pVMH19 or pVMH20 in BW27883 or BW27883 Δ*prc* was monitored by Western blotting using an anti-FLAG antibody. **(C)** Analysis of the cell extracts used for Western blot in **(B)** using SDS-PAGE and Coomassie staining, showing a uniform level of proteins for all samples. In **(B)** and **(C)**, the expression of FLAG-tagged SddA was induced by incubation with 1mM IPTG for 4 h. **(D)** Left: AlphaFold3 prediction showing an interaction between Prc (colored in pink) and the C-terminal region of SddA (colored in cyan); Middle: focus on the active site residues Ser452 and Lys477 of Prc (CPK representation)[48] interacting with the C-terminal residues KFNASAN of SddA; Right: Predicted Aligned Error (PAE), with SddA top-left and Prc bottom-right (blue, orange and white coding for high, medium and low prediction confidence, respectively).

**S8 Fig.**
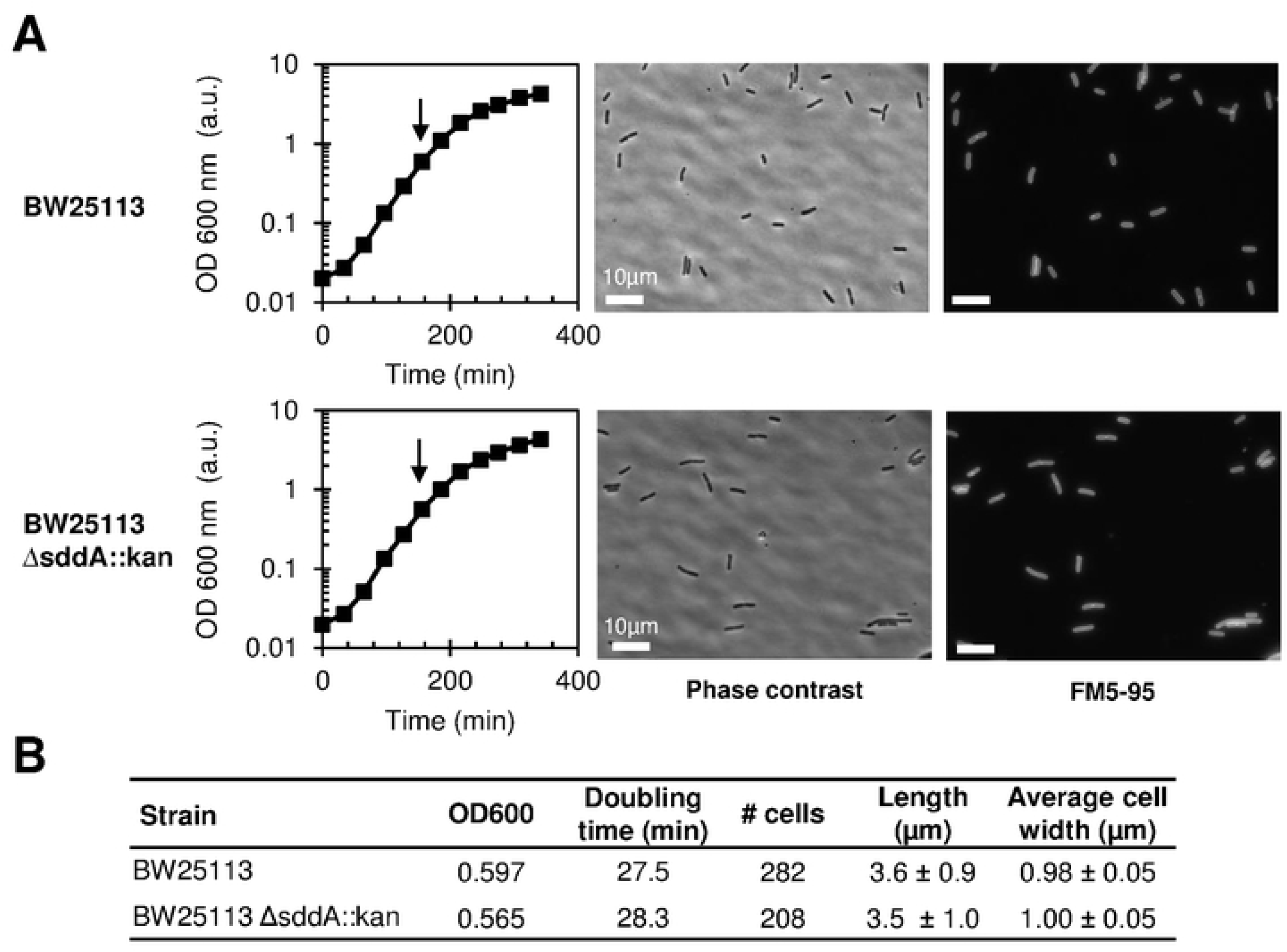
Deletion of *sddA* does not cause morphological changes when growing in LB. **(A)** BW25113 or BW25513 Δ*ssdA*::kan were grown in LB at 37°C. Samples were collected (arrows), fixed, stained with FM5-95 (cell membrane), immobilized and imaged by phase contrast and epifluorescence microscopy. Representative images are shown. Scale bar is 10 µm. **(B)** morphological measurements and growth doubling time of the cells and growth curves shown in A.

**S9 Fig.**
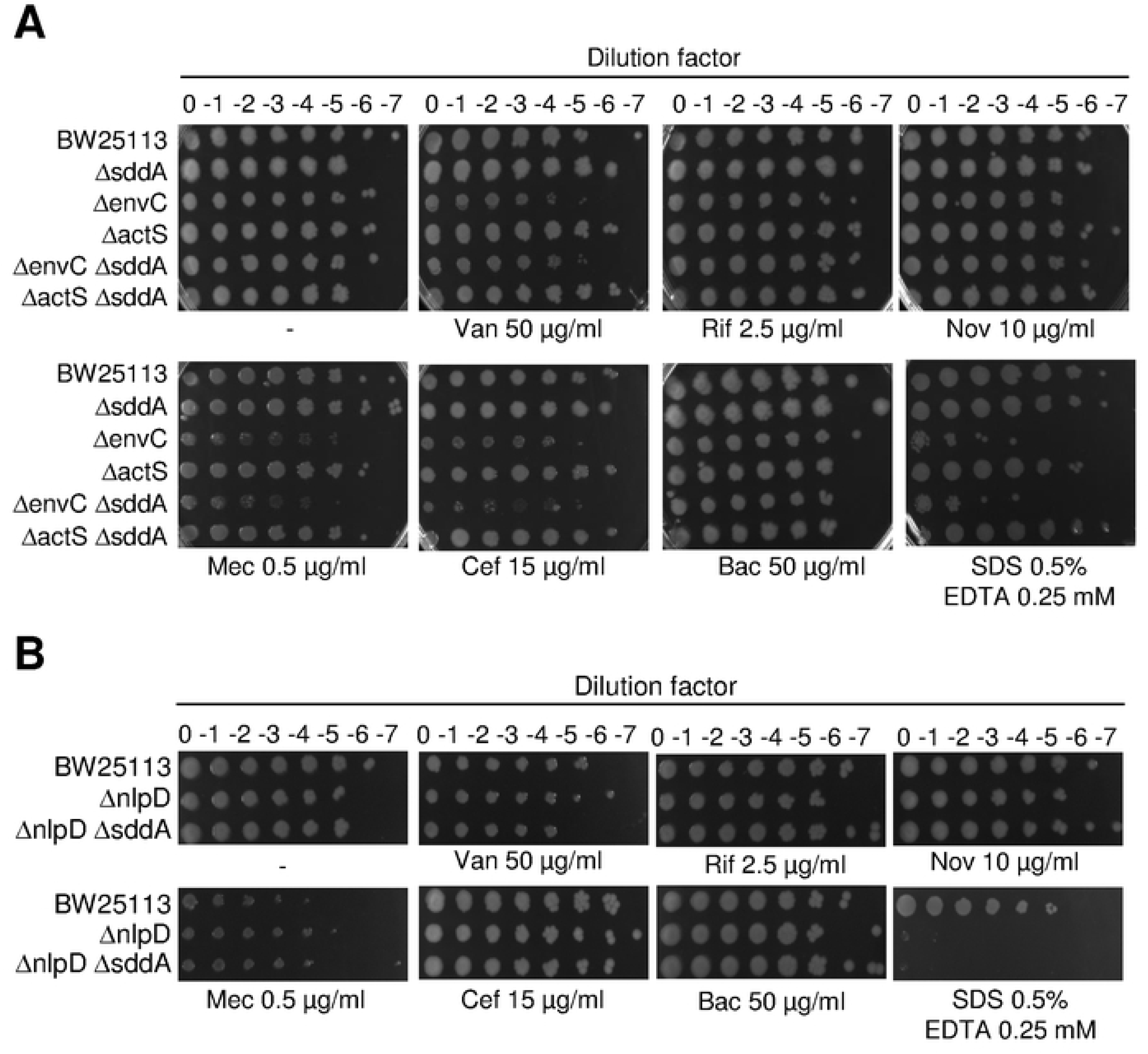
Deletion of *actS*, *nlpD* or *envC* does not affect OM permeability of cells lacking SddA. **(A)** Overnight cultures of BW25113 and isogenic Δ*sddA,* Δ*envC,* Δ*actS,* Δ*envC* Δ*sddA,* Δ*actS* Δ*sddA* mutants and **(B)** of BW25113 Δ*nlpD* and isogenic Δ*nlpD* Δ*sddA* were serially diluted and spotted on LB with 5% NaCl plates containing vancomycin (Van), rifampicin (Rif), novobiocin (Nov), mecillinam (Mec), cefsulodin (Cefs), bacitracin (Bac) or SDS/EDTA at the indicated concentrations. Plates were incubated at 37°C for 24 h.

**S10 Fig.**
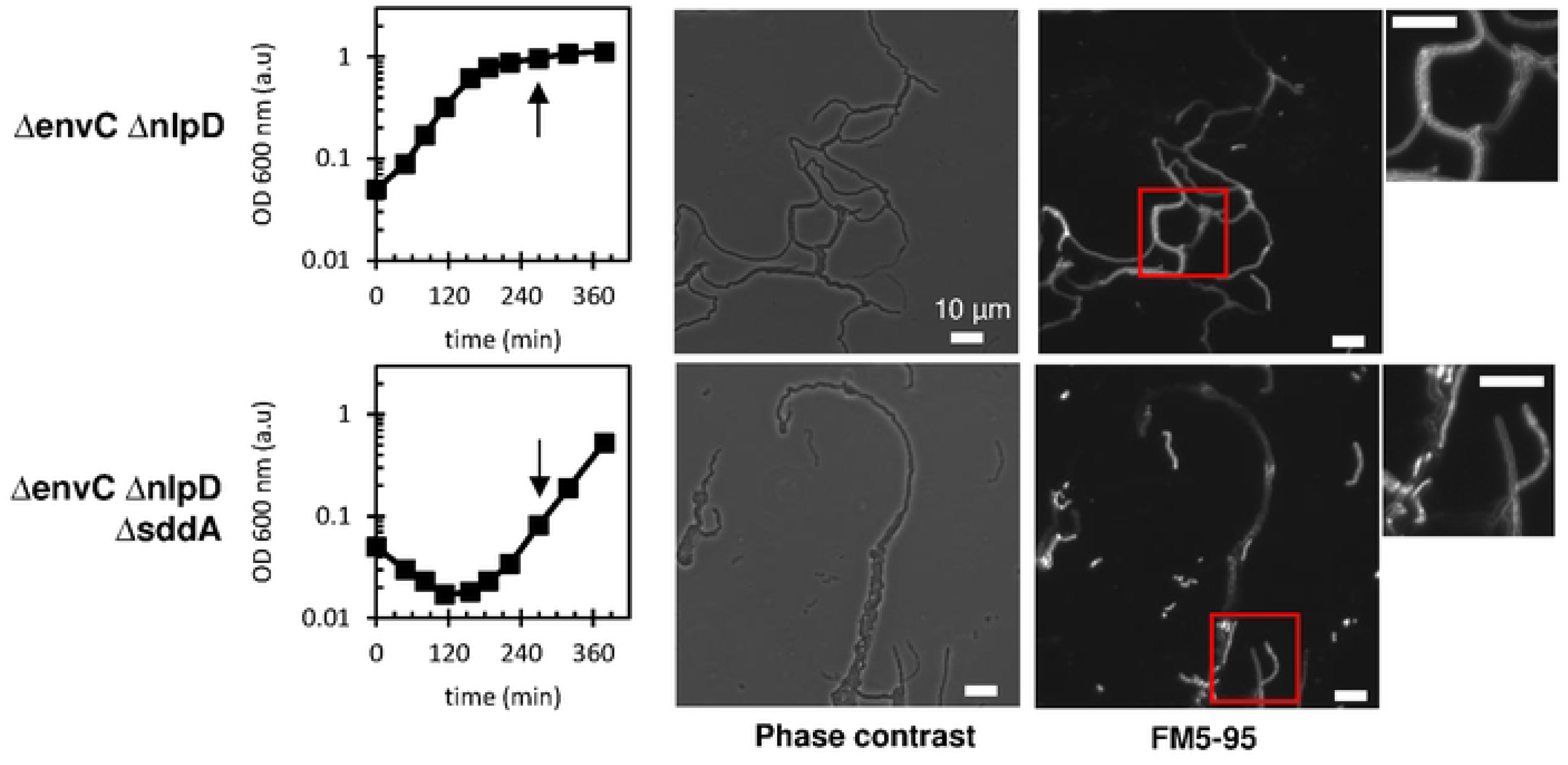
SddA deletion increases the sick phenotype of Δ*envC* Δ*nlpD*. BW25113 Δ*envC* Δ*nlpD* and BW25113 Δ*envC* Δ*nlpD* Δ*sddA* were grown in LB at 37°C. At indicated times, samples were taken and imaged on agarose pads containing FM5-95 membrane stain. All scale bars represent 10 µM.

**S11 Fig.**
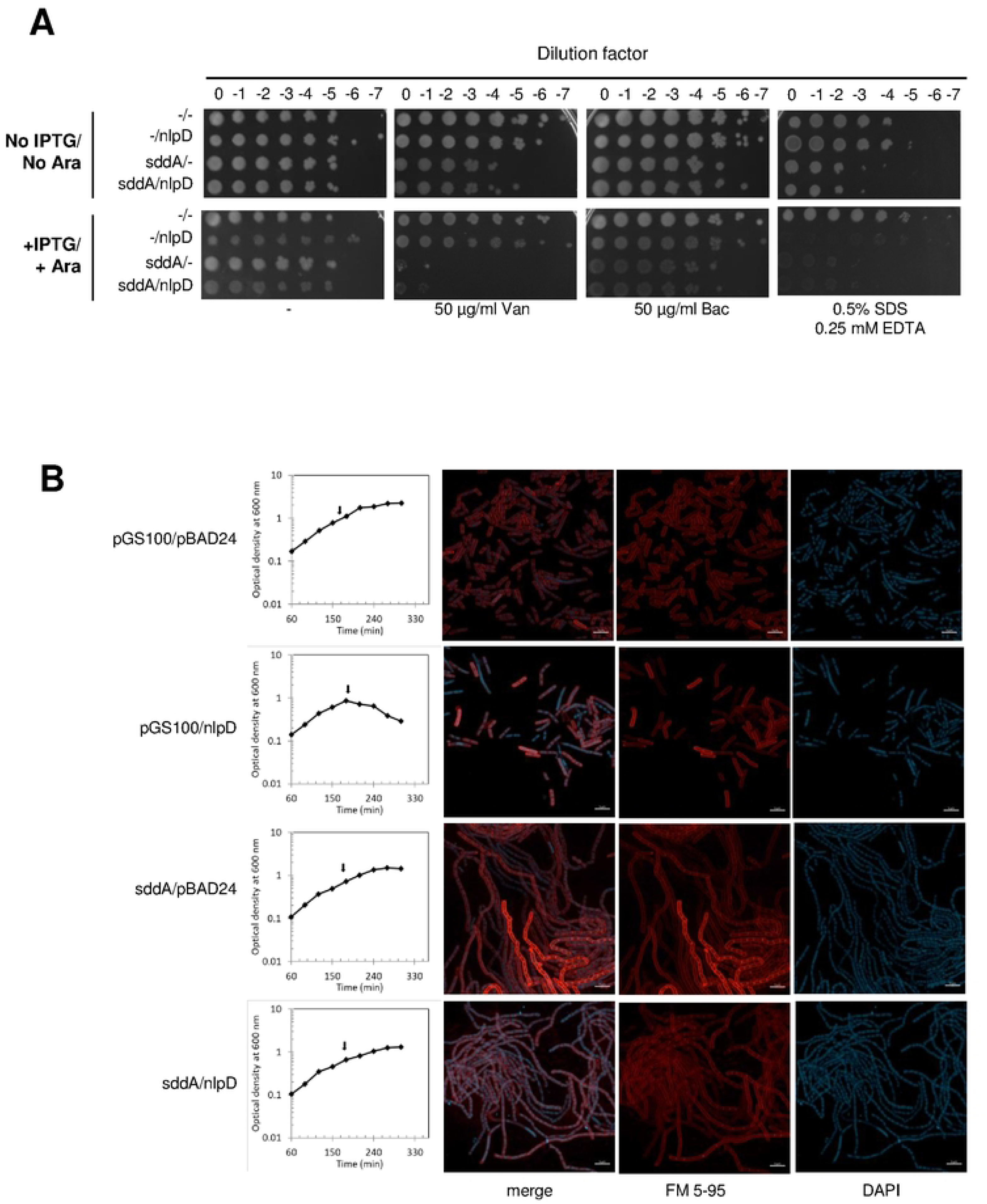
Co-expression of SddA and NlpD does not rescue OM defects and cell chaining. **(A)** Overnight cultures of BW25113 harbouring empty pGS100 or pBAD24 plasmids (-/-) or pGS100 with *sddA* (*sddA*/-) or pBAD24 with *nlpD* (-/*nlpD*) or both (*ssdA*/*nlpD*) were serially diluted and spotted onto LB-Lennox supplemented with 25 μg ml^-1^ chloramphenicol and 100 μg ml^-1^ ampicillin containing vancomycin (Van), bacitracin (Bac) or SDS/EDTA at the indicated concentrations. 0.2% arabinose and 0.5 mM IPTG were used to induce *nlpD* and *sddA* expression, respectively. **(B)** BW25113 cells harbouring pGS100 and pBAD24 or pBAD24 with *nlpD* (pGS100/*nlpD*), pGS100 with sddA (*sddA*/pBAD24) or both (*ssdA*/*nlpD*) were grown in LB with 5% NaCl supplemented with 25 μg ml^-1^ chloramphenicol and 100 μg ml^-1^ ampicillin. IPTG (0.5 mM) and arabinose (0.2%) were used to induce the expression of *sddA* and *nlpD*, respectively. Samples were collected at the exponential growth phase (arrows) stained with FM5-95 (red, cell membrane) and DAPI (blue, nucleoid), immobilized and imaged by confocal fluorescence microscopy. Representative images are shown. Scale bar is 5 µm.

**S12 Fig.**
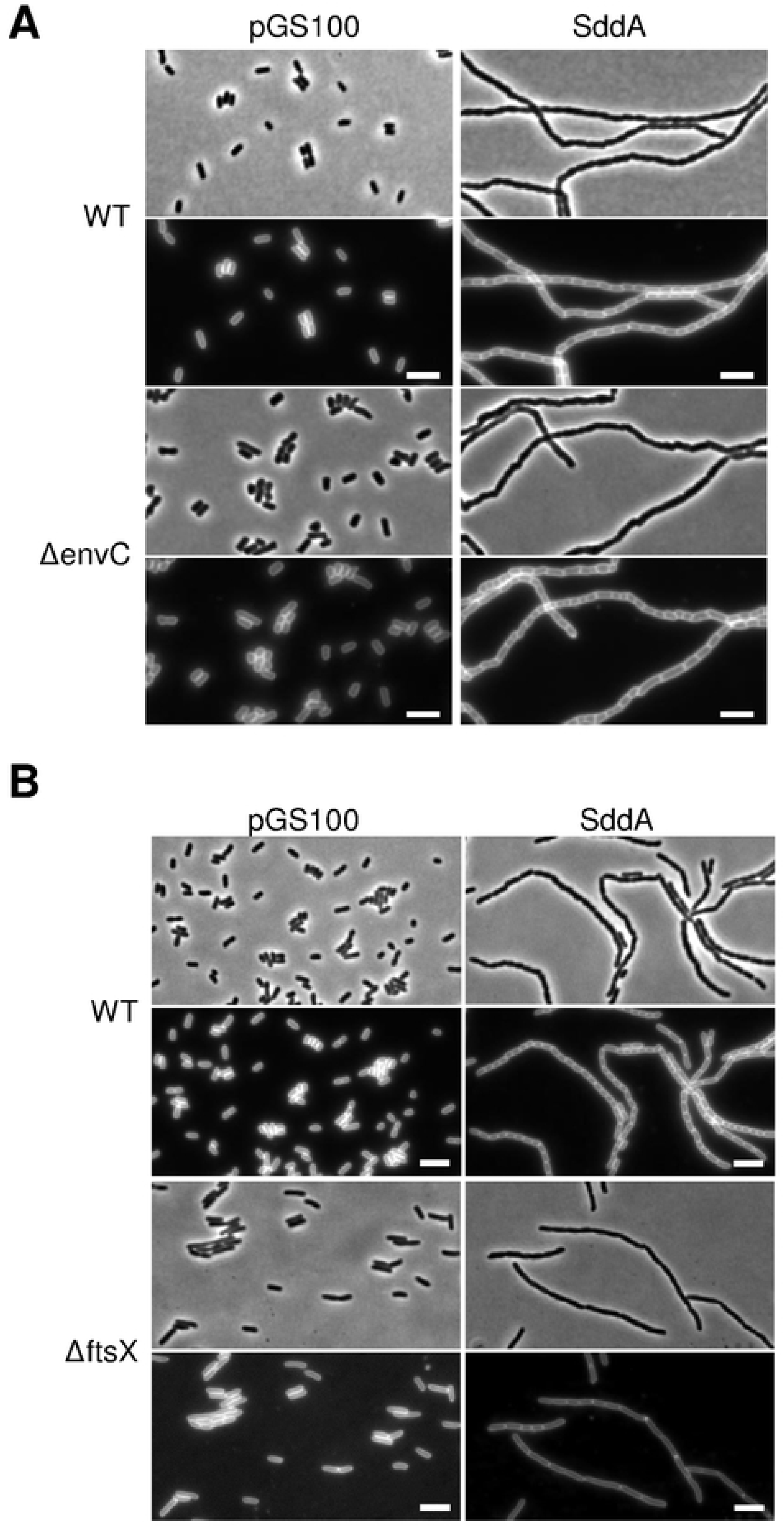
Cell chaining phenotype caused by SddA overproduction is independent of EnvC or FtsX. **(A)**BW25113 (WT) and BW25113 Δ*envC* (Δ*envC*) cells harbouring pGS100 (ev) or pGS100 expressing *sddA* (SddA) were grown at 37°C in LB with 20 µg ml^-1^ chloramphenicol and expression was induced with 0.5 mM IPTG for 140 min. Samples were imaged by phase contrast and fluorescence microscopy (FM5- 95). **(B)** BW25113 (WT) and BW25113 Δ*ftsX* (Δ*ftsX*) cells harbouring pGS100 (ev) or pGS100 expressing *sddA* (SddA) were grown at 30°C in LB with 20 µg ml^-1^ chloramphenicol supplemented with 0.2 M sucrose and expression was induced with 0.5 mM IPTG for 230 min. Samples were imaged by phase contrast and fluorescence microscopy (FM5-95). Representative images are shown. Scale bar is 5 µm.

**S13 Fig.**
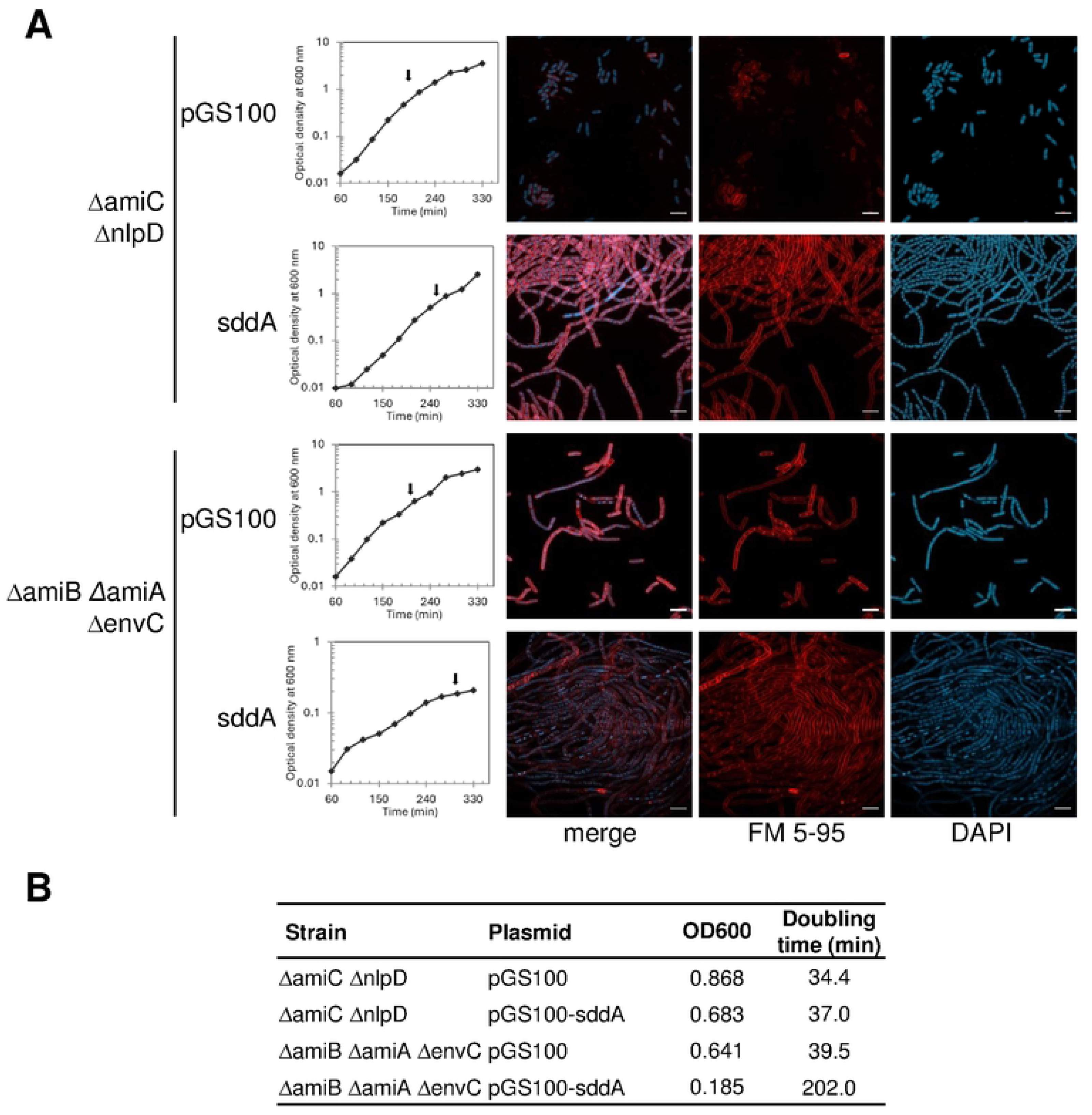
High *sddA* expression in mutants defective in amidase activation aggravates their cell chaining phenotype. **(A)** Δ*amiC* Δ*nlpD* and Δ*amiB ΔamiA* Δ*envC* cells harbouring pGS100 or pGS100 expressing *sddA* were grown in LB with 5% NaCl supplemented with chloramphenicol at 25 µg ml^-1^. Samples were collected (arrows) stained with FM5-95 (red, cell membrane) and DAPI (blue, nucleoid), immobilized and imaged by confocal fluorescence microscopy. Representative images are shown. Scale bar is 5 µm. **(B)** Doubling time of cultures shown in panel A.

**S14 Fig.**
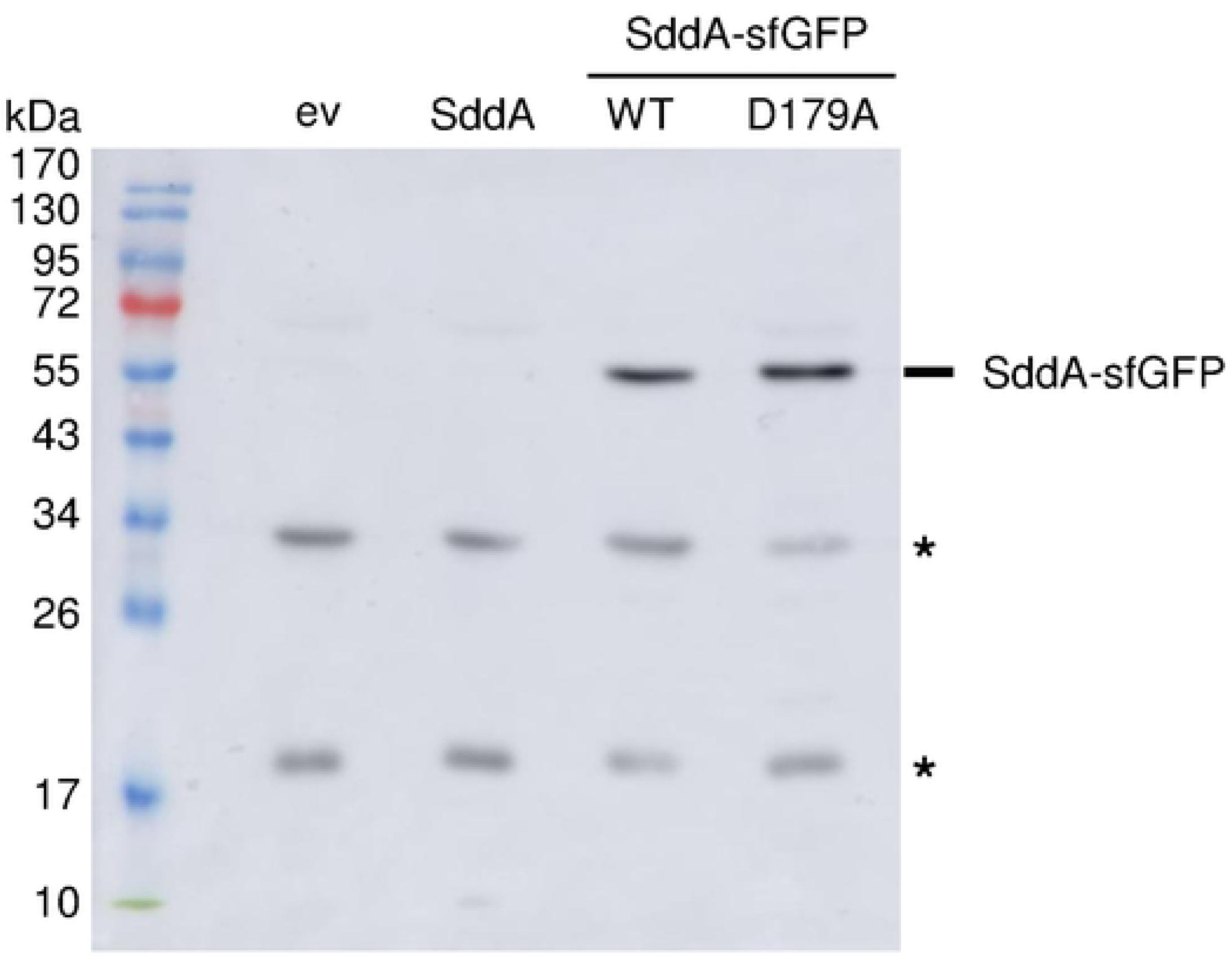
Stability of overproduced SddA-sfGFP constructs. BW25113 cells harbouring pGS100 (ev) or pGS100 expressing *sddA*, *sddA::sfgfp* (WT) or *sddA D179A::sfgfp* (D179A) were grown in LB with 20 µg ml^-1^ chloramphenicol at 37°C and expression of the fluorescent constructs was induced with 0.5 mM IPTG for 140 min. Samples were pelleted, protein extracts were obtained by sonication and quantified by BCA. 15 µg of each extract was loaded per lane, separated by SDS-PAGE and GFP was immunodetected by Western blot using α-GFP antibody. (*, unspecific bands)

**S15 Fig.**
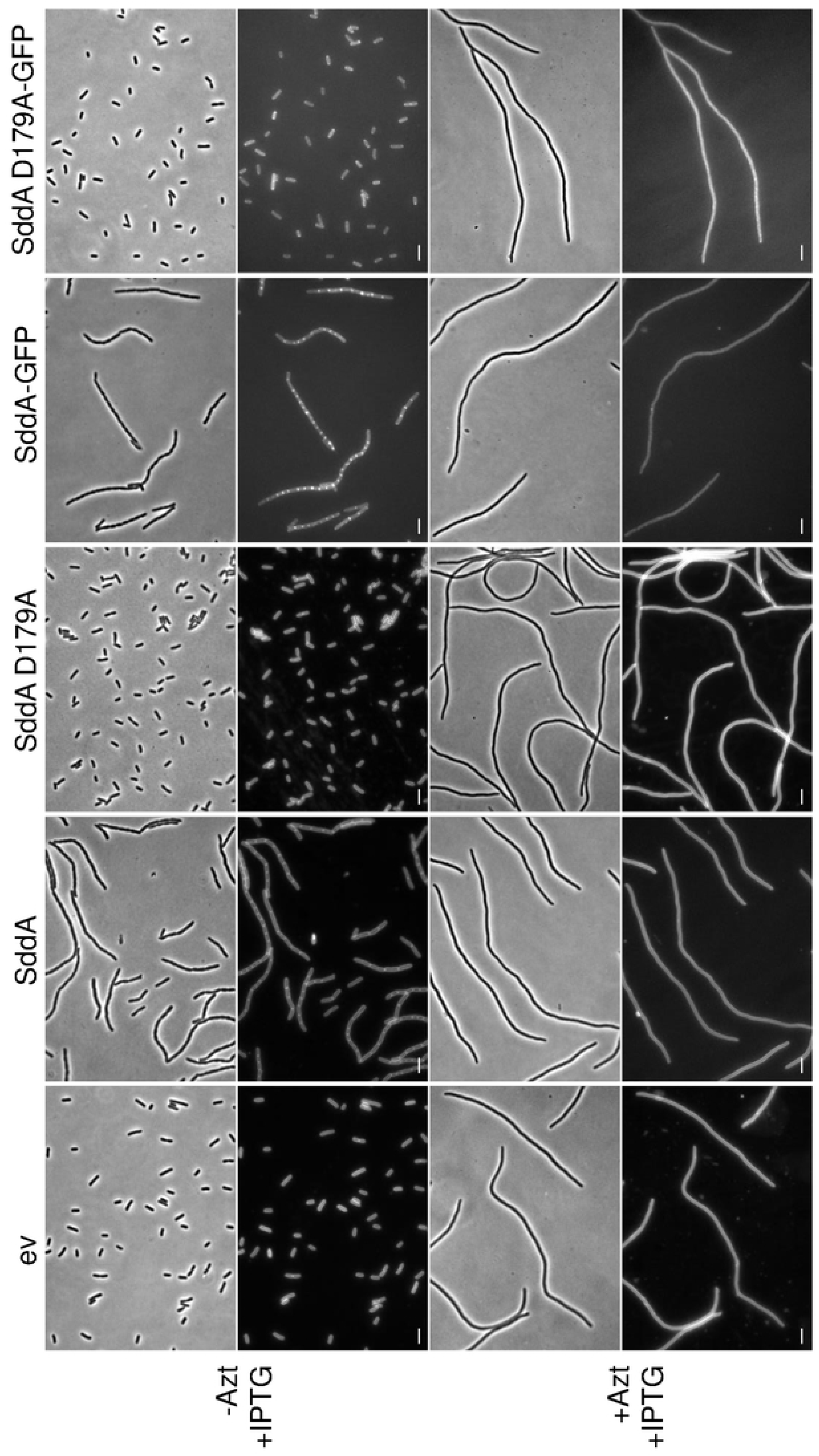
Sdd*A* localization is dependent on septal peptidoglycan synthesis. BW25113 cells harbouring pGS100 (ev), pGS100 encoding *sddA, sddA D179A, sddA::sfgfp,* or *sddA D179A::sfgfp* were grown in LB with 20 µg ml^-1^ of chloramphenicol at 37°C. When indicated, 1 µg ml^-1^ of aztreonam was added. Expression of the plasmid-encoded constructs was induced 30 min later by addition of 0.5 mM IPTG for 80 min. Samples were imaged by phase contrast and fluorescence microscopy (FM5-95 or GFP). Representative images are shown. Scale bar is 5 µm.

**S16 Fig.**
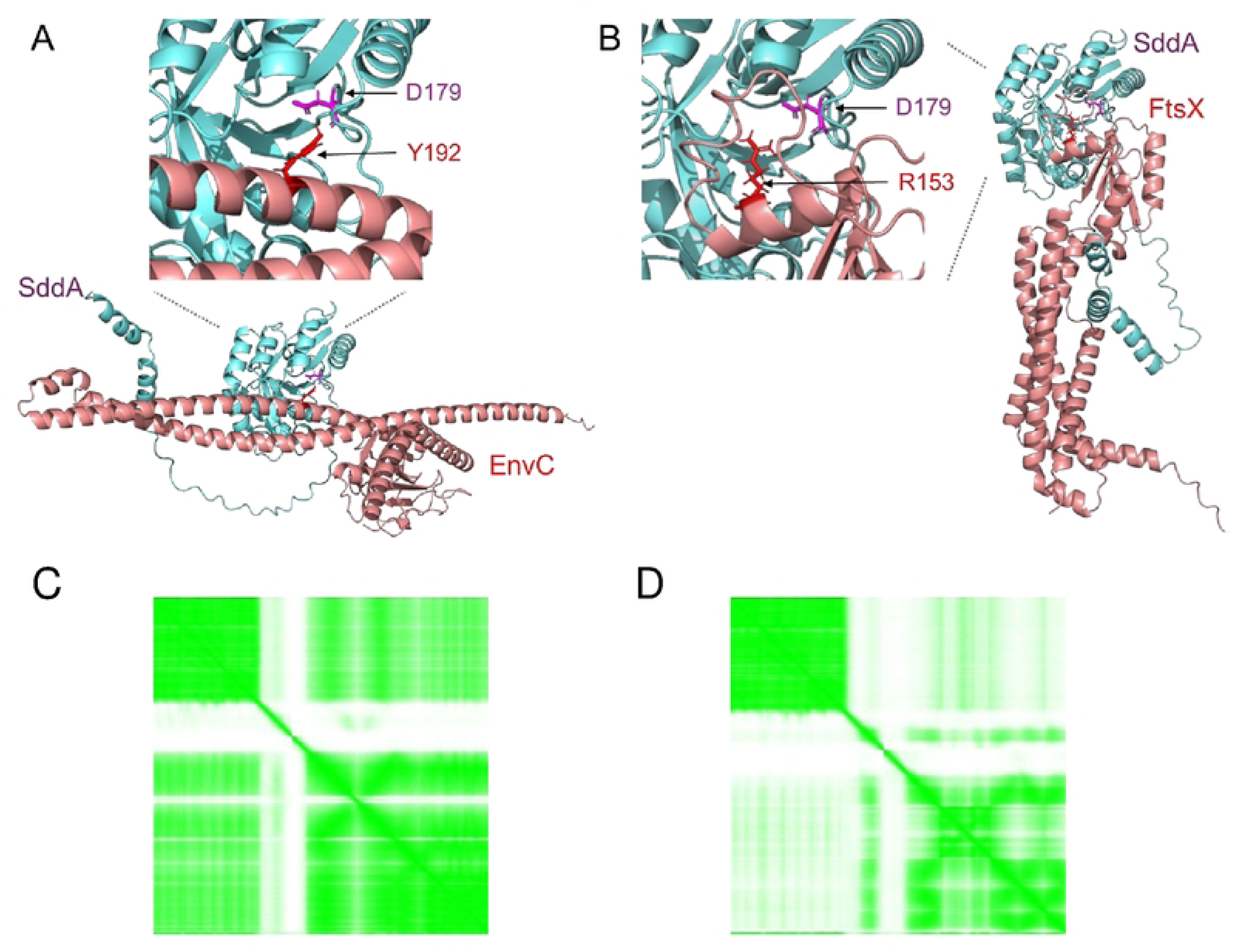
Alphafold-predicted interaction between SddA and EnvC or SddA and FtsX. Best models obtained by Alphafold of hypothetical SddA-EnvC, **(A)**, and SddA-FtsX complexes, **(B)**; and their corresponding Predicted Aligned Error (PAE) matrices **(C)** and **(D)**, respectively. Each cell in the PAE matrices represents the estimate alignment error (in Å) between pairs of residues across the predicted protein structure, with darker shades indicating lower PAE and higher model confidence.

**S17 Fig.**
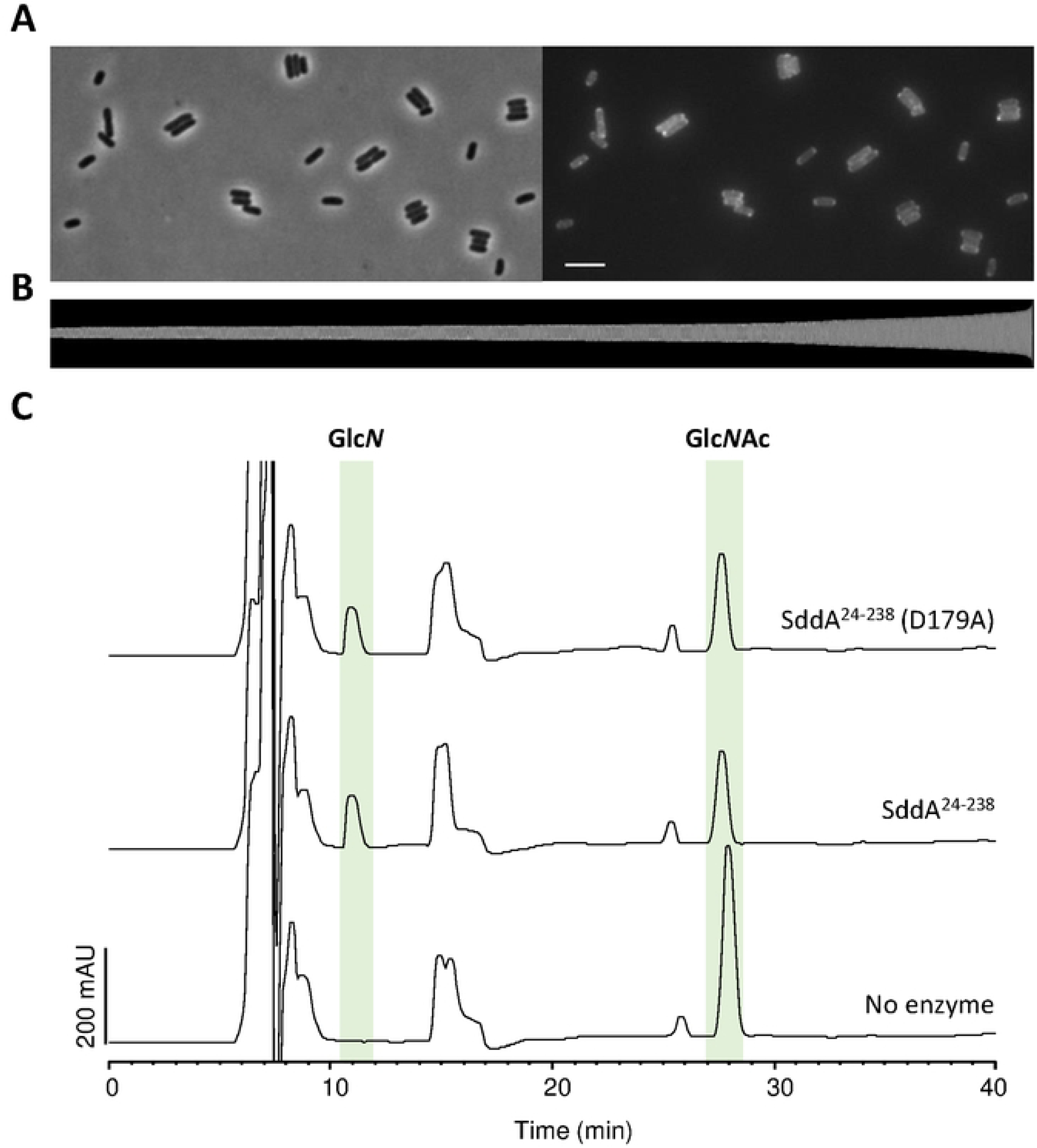
SddA D179A is active against denuded strands and its sfGFP-fusion does not localise to mid-cell and does not cause a cell chaining phenotype. **(A)** Phase contrast and fluorescence microscopy images of BW25113 harbouring pMP116 (pGS100 *sddA D179A::sfgfp*). Scale bar equals 5 µm. **(B)** Demograpth of SddA D179A-sfGFP localization in cells sorted according to their cell length (from 1.43 µm to 6.4 µm, n = 1555). The fluorescence signal does not show an increase of the intensity at midcell. **(C)** Chromatograms of the analysis of denuded glycan strands treated first with buffer, His-SddA^24-238^ or His-SddA^24-238^ D179A, and then with the lytic transglycosylase MltA. MltA fully digested denuded strands producing anhydro sugars. Reactions contained 2 µM of enzyme and were incubated at 37°C for 24 h. Peak labelled Glc*N* corresponds to Glc*N*-Mur*N*AcAnh and peak labelled Glc*N*Ac corresponds to Glc*N*Ac-Mur*N*AcAnh.

**S18 Fig.**
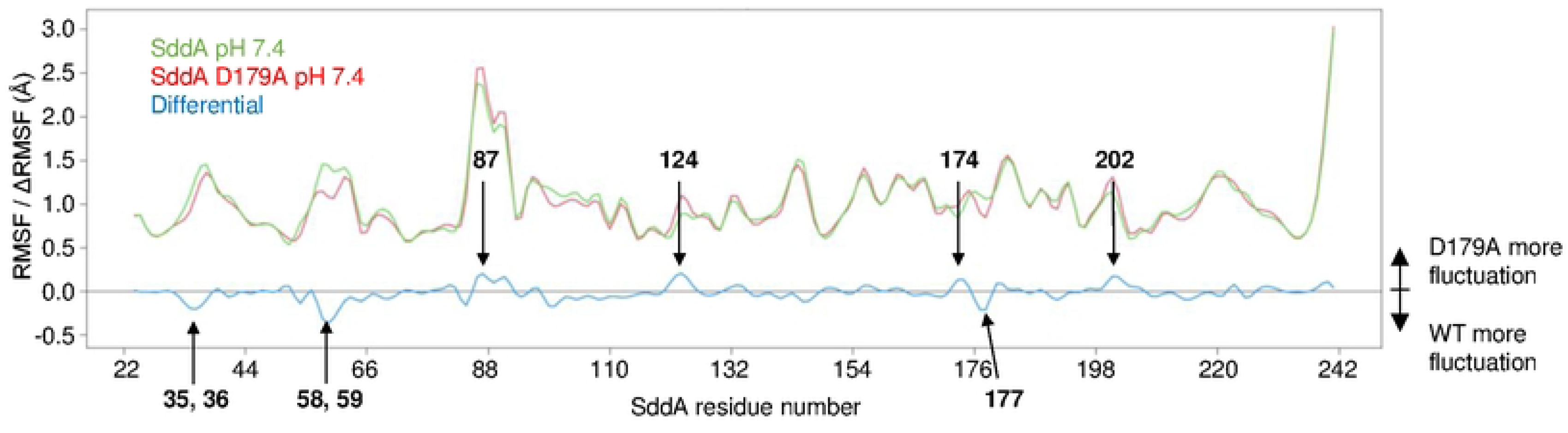
The mutation D179A induces localized and global changes in SddA dynamics, particularly in regions critical for binding to EnvC. Residue-resolved root mean square fluctuations (RMSF) of wild-type (WT, green) and D179A mutant (red) SddA, calculated from molecular dynamics simulations. The blue curve represents the difference in fluctuations (ΔRMSF) between WT and mutant (WT - mutant). Negative ΔRMSF values indicate greater rigidity in the mutant, while positive values reflect increased flexibility. Residues involved in interaction with EnvC, as predicted by AlphaFold, are highlighted in the analysis.

**S19 Fig.**
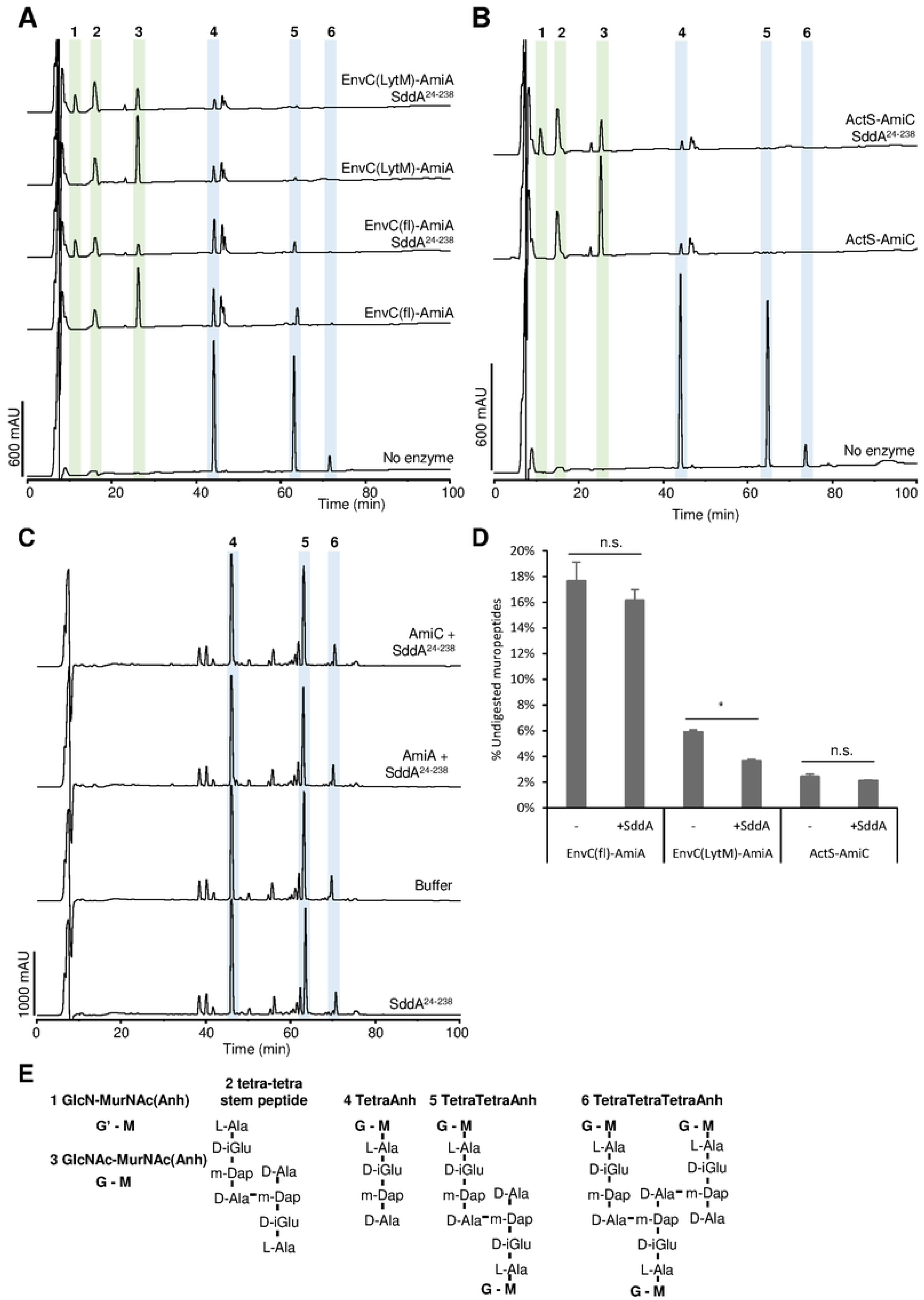
SddA does not significantly inhibit ActS-AmiC or EnvC-AmiA. (A, **B)** HPLC-based end-point activity assays for EnvC-AmiA or ActS-AmiC amidase-activator pairs, in the presence or absence of SddA^24-238^. Sacculi from *E. coli* BW25113Δ6LDT were incubated with 2 µM of the indicated amidase and activator, in the presence or absence of 4 µM His-SddA^24-238^ (SddA^24-238^). EnvC(fl) indicates the EnvC construct containing the LytM and coiled-coiled domain whereas EnvC(LytM) indicates the construct with only the LytM domain. Reactions were incubated for 2 h **(A)** or 1 h **(B)** at 37°C. Representative chromatograms are shown. **(C)** Chromatograms for the control reactions showing no activation of AmiA or AmiC by SddA^24-238^. Reactions contained 2 µM of each protein and *E. coli* MC1061 PG and were incubated for 2 h at 37°C. **(D)** Quantification of muropeptides peak areas (peaks 4, 5 and 6) in reactions depicted in A and B, normalized against the areas of those peaks in the controls with no enzyme. Values are average +/- variation of 2 reactions. There was a slight activation of EnvC(LytM)- AmiA activity by SddA. **(E)** Identity and structures of the peaks labelled in A-C and Fig 6A. G stands for Glc*N*Ac, M for Mur*N*AcAnh and G’ for Glc*N*.

**S1 Table. Strains used in this work.**

**S2 Table. Plasmids used in this work.**

**S3 Table. Oligonucleotides used in this work.**

**S1 References.**

**Supplementary references.**

## REFERENCES

1. Vollmer W, Blanot D, de Pedro MA. Peptidoglycan structure and architecture. FEMS Microbiol Rev. 2008;32(2):149–67. Epub 20080108. doi: 10.1111/j.1574-6976.2007.00094.x. PubMed PMID: 18194336.

2. Egan AJF, Errington J, Vollmer W. Regulation of peptidoglycan synthesis and remodelling. Nat Rev Microbiol. 2020;18(8):446–60. Epub 20200518. doi: 10.1038/s41579-020-0366-3. PubMed PMID: 32424210.

3. Typas A, Banzhaf M, Gross CA, Vollmer W. From the regulation of peptidoglycan synthesis to bacterial growth and morphology. Nat Rev Microbiol. 2011;10(2):123–36. Epub 20111228. doi: 10.1038/nrmicro2677. PubMed PMID: 22203377; PubMed Central PMCID: PMCPMC5433867.

4. Gray AN, Egan AJ, Van’t Veer IL, Verheul J, Colavin A, Koumoutsi A, et al. Coordination of peptidoglycan synthesis and outer membrane constriction during *Escherichia coli* cell division. eLife. 2015;4:e07118. Epub 2015/05/08. doi: 10.7554/eLife.07118. PubMed PMID: 25951518; PubMed Central PMCID: PMCPMC4458516.

5. Mamou G, Corona F, Cohen-Khait R, Housden NG, Yeung V, Sun D, et al. Peptidoglycan maturation controls outer membrane protein assembly. Nature. 2022;606(7916):953-9. doi: 10.1038/s41586-022-04834-7.

6. Bisson-Filho AW, Hsu Y-P, Squyres GR, Kuru E, Wu F, Jukes C, et al. Treadmilling by FtsZ filaments drives peptidoglycan synthesis and bacterial cell division. Science. 2017;355(6326):739-43. doi: 10.1126/science.aak9973.

7. Yang X, Lyu Z, Miguel A, McQuillen R, Huang KC, Xiao J. GTPase activity–coupled treadmilling of the bacterial tubulin FtsZ organizes septal cell wall synthesis. Science. 2017;355(6326):744-7. doi: 10.1126/science.aak9995.

8. Monteiro JM, Pereira AR, Reichmann NT, Saraiva BM, Fernandes PB, Veiga H, et al. Peptidoglycan synthesis drives an FtsZ-treadmilling-independent step of cytokinesis. Nature. 2018;554(7693):528-32. doi: 10.1038/nature25506.

9. Mahone CR, Goley ED. Bacterial cell division at a glance. J Cell Sci. 2020;133(7). Epub 20200408. doi: 10.1242/jcs.237057. PubMed PMID: 32269092; PubMed Central PMCID: PMCPMC7157936.

10. Pazos M, Vollmer W. Regulation and function of class A Penicillin-binding proteins. Current Opinion in Microbiology. 2021;60:80–7. doi: 10.1016/j.mib.2021.01.008.

11. Pazos M, Peters K, Casanova M, Palacios P, VanNieuwenhze M, Breukink E, et al. Z-ring membrane anchors associate with cell wall synthases to initiate bacterial cell division. Nat Commun. 2018;9(1):5090. Epub 20181130. doi: 10.1038/s41467-018-07559-2. PubMed PMID: 30504892; PubMed Central PMCID: PMCPMC6269477.

12. Pazos M, Peters K, Boes A, Safaei Y, Kenward C, Caveney NA, et al. SPOR proteins are required for functionality of class A penicillin-binding proteins in *Escherichia coli*. mBio. 2020;11(6). doi: 10.1128/mBio.02796-20.

13. Pazos M, Peters K, Vollmer W. Robust peptidoglycan growth by dynamic and variable multi- protein complexes. Curr Opin Microbiol. 2017;36:55–61. Epub 20170215. doi: 10.1016/j.mib.2017.01.006. PubMed PMID: 28214390.

14. Heidrich C, Templin MF, Ursinus A, Merdanovic M, Berger J, Schwarz H, et al. Involvement of N-acetylmuramyl-L-alanine amidases in cell separation and antibiotic-induced autolysis of *Escherichia coli*. Mol Microbiol. 2001;41(1):167–78. doi: 10.1046/j.1365-2958.2001.02499.x. PubMed PMID: 11454209.

15. Priyadarshini R, de Pedro MA, Young KD. Role of peptidoglycan amidases in the development and morphology of the division septum in *Escherichia coli*. J Bacteriol. 2007;189(14):5334–47. Epub 20070504. doi: 10.1128/JB.00415-07. PubMed PMID: 17483214; PubMed Central PMCID: PMCPMC1951850.

16. Uehara T, Dinh T, Bernhardt TG. LytM-domain factors are required for daughter cell separation and rapid ampicillin-induced lysis in *Escherichia coli*. J Bacteriol. 2009;191(16):5094–107. Epub 20090612. doi: 10.1128/JB.00505-09. PubMed PMID: 19525345; PubMed Central PMCID: PMCPMC2725582.

17. Chen Y, Gu J, Yang B, Yang L, Pang J, Luo Q, et al. Structure and activity of the septal peptidoglycan hydrolysis machinery crucial for bacterial cell division. PLoS Biol. 2024;22(5):e3002628. Epub 20240530. doi: 10.1371/journal.pbio.3002628. PubMed PMID: 38814940; PubMed Central PMCID: PMCPMC11139282.

18. Hao A, Suo Y, Lee SY. Structural insights into the FtsEX-EnvC complex regulation on septal peptidoglycan hydrolysis in *Vibrio cholerae*. Structure. 2024;32(2):188–99 e5. Epub 20231208. doi: 10.1016/j.str.2023.11.007. PubMed PMID: 38070498.

19. Xu X, Li J, Chua WZ, Pages MA, Shi J, Hermoso JA, et al. Mechanistic insights into the regulation of cell wall hydrolysis by FtsEX and EnvC at the bacterial division site. Proc Natl Acad Sci U S A. 2023;120(21):e2301897120. Epub 20230515. doi: 10.1073/pnas.2301897120. PubMed PMID: 37186861; PubMed Central PMCID: PMCPMC10214136.

20. Cook J, Baverstock TC, McAndrew MBL, Roper DI, Stansfeld PJ, Crow A. Activator-induced conformational changes regulate division-associated peptidoglycan amidases. Proc Natl Acad Sci U S A. 2023;120(24):e2302580120. Epub 20230605. doi: 10.1073/pnas.2302580120. PubMed PMID: 37276423; PubMed Central PMCID: PMCPMC10268282.

21. Gurnani Serrano CK, Winkle M, Martorana AM, Biboy J, More N, Moynihan P, et al. ActS activates peptidoglycan amidases during outer membrane stress in *Escherichia coli*. Mol Microbiol. 2021;116(1):329–42. Epub 20210323. doi: 10.1111/mmi.14712. PubMed PMID: 33660879; PubMed Central PMCID: PMCPMC8360153.

22. Mueller EA, Iken AG, Ali Ozturk M, Winkle M, Schmitz M, Vollmer W, et al. The active repertoire of *Escherichia coli* peptidoglycan amidases varies with physiochemical environment. Mol Microbiol. 2021;116(1):311–28. Epub 20210403. doi: 10.1111/mmi.14711. PubMed PMID: 33666292; PubMed Central PMCID: PMCPMC8295211.

23. Tsang MJ, Yakhnina AA, Bernhardt TG. NlpD links cell wall remodeling and outer membrane invagination during cytokinesis in *Escherichia coli*. PLoS Genet. 2017;13(7):e1006888. Epub 20170714. doi: 10.1371/journal.pgen.1006888. PubMed PMID: 28708841; PubMed Central PMCID: PMCPMC5533458.

24. Boelter G, Bryant JA, Doherty H, Wotherspoon P, Alodaini D, Ma X, et al. The lipoprotein DolP affects cell separation in *Escherichia coli*, but not as an upstream regulator of NlpD. Microbiology (Reading). 2022;168(5). doi: 10.1099/mic.0.001197. PubMed PMID: 35604759.

25. Szczepaniak J, Press C, Kleanthous C. The multifarious roles of Tol-Pal in Gram-negative bacteria. FEMS Microbiol Rev. 2020;44(4):490–506. doi: 10.1093/femsre/fuaa018. PubMed PMID: 32472934; PubMed Central PMCID: PMCPMC7391070.

26. Vollmer W. Structural variation in the glycan strands of bacterial peptidoglycan. FEMS Microbiol Rev. 2008;32(2):287–306. Epub 20071205. doi: 10.1111/j.1574-6976.2007.00088.x. PubMed PMID: 18070068.

27. Vollmer W, Tomasz A. The *pgdA* gene encodes for a peptidoglycan *N*-acetylglucosamine deacetylase in *Streptococcus pneumoniae*. J Biol Chem. 2000;275(27):20496–501. doi: 10.1074/jbc.M910189199. PubMed PMID: 10781617.

28. Vollmer W, Tomasz A. Peptidoglycan *N*-acetylglucosamine deacetylase, a putative virulence factor in *Streptococcus pneumoniae*. Infect Immun. 2002;70(12):7176–8. doi: 10.1128/IAI.70.12.7176-7178.2002. PubMed PMID: 12438406; PubMed Central PMCID: PMCPMC133073.

29. Boneca IG, Dussurget O, Cabanes D, Nahori MA, Sousa S, Lecuit M, et al. A critical role for peptidoglycan *N*-deacetylation in *Listeria* evasion from the host innate immune system. Proc Natl Acad Sci U S A. 2007;104(3):997–1002. Epub 20070110. doi: 10.1073/pnas.0609672104. PubMed PMID: 17215377; PubMed Central PMCID: PMCPMC1766339.

30. Fittipaldi N, Sekizaki T, Takamatsu D, de la Cruz Dominguez-Punaro M, Harel J, Bui NK, et al. Significant contribution of the *pgdA* gene to the virulence of *Streptococcus suis*. Mol Microbiol. 2008;70(5):1120–35. doi: 10.1111/j.1365-2958.2008.06463.x. PubMed PMID: 18990186.

31. Blair DE, Schuttelkopf AW, MacRae JI, van Aalten DM. Structure and metal-dependent mechanism of peptidoglycan deacetylase, a streptococcal virulence factor. Proc Natl Acad Sci U S A. 2005;102(43):15429–34. Epub 20051012. doi: 10.1073/pnas.0504339102. PubMed PMID: 16221761; PubMed Central PMCID: PMCPMC1252587.

32. Wang G, Maier SE, Lo LF, Maier G, Dosi S, Maier RJ. Peptidoglycan deacetylation in *Helicobacter pylori* contributes to bacterial survival by mitigating host immune responses. Infect Immun. 2010;78(11):4660–6. Epub 20100830. doi: 10.1128/IAI.00307-10. PubMed PMID: 20805339; PubMed Central PMCID: PMCPMC2976313.

33. Shaik MM, Cendron L, Percudani R, Zanotti G. The structure of *Helicobacter pylori* HP0310 reveals an atypical peptidoglycan deacetylase. PLoS One. 2011;6(4):e19207. Epub 20110429. doi: 10.1371/journal.pone.0019207. PubMed PMID: 21559431; PubMed Central PMCID: PMCPMC3084791.

34. Sycuro LK, Pincus Z, Gutierrez KD, Biboy J, Stern CA, Vollmer W, et al. Peptidoglycan crosslinking relaxation promotes *Helicobacter pylori*’s helical shape and stomach colonization. Cell. 2010;141(5):822–33. doi: 10.1016/j.cell.2010.03.046. PubMed PMID: 20510929; PubMed Central PMCID: PMCPMC2920535.

35. Sycuro LK, Wyckoff TJ, Biboy J, Born P, Pincus Z, Vollmer W, et al. Multiple peptidoglycan modification networks modulate *Helicobacter pylori*’s cell shape, motility, and colonization potential. PLoS Pathog. 2012;8(3):e1002603. Epub 20120322. doi: 10.1371/journal.ppat.1002603. PubMed PMID: 22457625; PubMed Central PMCID: PMCPMC3310797.

36. Chaput C, Ecobichon C, Cayet N, Girardin SE, Werts C, Guadagnini S, et al. Role of AmiA in the morphological transition of *Helicobacter pylori* and in immune escape. PLoS Pathog. 2006;2(9):e97. doi: 10.1371/journal.ppat.0020097. PubMed PMID: 17002496; PubMed Central PMCID: PMCPMC1574363.

37. Chaput C, Labigne A, Boneca IG. Characterization of *Helicobacter pylori* lytic transglycosylases Slt and MltD. J Bacteriol. 2007;189(2):422–9. Epub 20061103. doi: 10.1128/JB.01270-06. PubMed PMID: 17085576; PubMed Central PMCID: PMCPMC1797392.

38. Patel AV, Turner RD, Rifflet A, Acosta-Martin AE, Nichols A, Awad MM, et al. PGFinder, a novel analysis pipeline for the consistent, reproducible, and high-resolution structural analysis of bacterial peptidoglycans. Elife. 2021;10. Epub 20210928. doi: 10.7554/eLife.70597. PubMed PMID: 34579805; PubMed Central PMCID: PMCPMC8478412.

39. Glauner B, Höltje JV, Schwarz U. The composition of the murein of Escherichia coli. J Biol Chem. 1988;263(21):10088–95. doi: 10.1016/S0021-9258(19)81481-3.

40. Glauner B. Separation and quantification of muropeptides with high-performance liquid chromatography. Anal Biochem. 1988;172(2):451–64. doi: 10.1016/0003-2697(88)90468-x. PubMed PMID: 3056100.

41. Itoh Y, Rice JD, Goller C, Pannuri A, Taylor J, Meisner J, et al. Roles of *pgaABCD* genes in synthesis, modification, and export of the *Escherichia coli* biofilm adhesin poly-beta-1,6-N-acetyl-D- glucosamine. J Bacteriol. 2008;190(10):3670–80. Epub 20080321. doi: 10.1128/JB.01920-07. PubMed PMID: 18359807; PubMed Central PMCID: PMCPMC2394981.

42. Wang X, Preston JF, 3rd, Romeo T. The *pgaABCD* locus of *Escherichia col*i promotes the synthesis of a polysaccharide adhesin required for biofilm formation. J Bacteriol. 2004;186(9):2724–34. doi: 10.1128/JB.186.9.2724-2734.2004. PubMed PMID: 15090514; PubMed Central PMCID: PMCPMC387819.

43. Hu P, Janga SC, Babu M, Diaz-Mejia JJ, Butland G, Yang W, et al. Global functional atlas of *Escherichia coli* encompassing previously uncharacterized proteins. PLoS Biol. 2009;7(4):e96. doi: 10.1371/journal.pbio.1000096. PubMed PMID: 19402753; PubMed Central PMCID: PMCPMC2672614.

44. Mendler K, Chen H, Parks DH, Lobb B, Hug LA, Doxey AC. AnnoTree: visualization and exploration of a functionally annotated microbial tree of life. Nucleic Acids Res. 2019;47(9):4442–8. doi: 10.1093/nar/gkz246. PubMed PMID: 31081040; PubMed Central PMCID: PMCPMC6511854.

45. Bui NK, Turk S, Buckenmaier S, Stevenson-Jones F, Zeuch B, Gobec S, et al. Development of screening assays and discovery of initial inhibitors of pneumococcal peptidoglycan deacetylase PgdA. Biochem Pharmacol. 2011;82(1):43–52. Epub 20110408. doi: 10.1016/j.bcp.2011.03.028. PubMed PMID: 21501597.

46. Lommatzsch J, Templin MF, Kraft AR, Vollmer W, Holtje JV. Outer membrane localization of murein hydrolases: MltA, a third lipoprotein lytic transglycosylase in *Escherichia coli*. J Bacteriol. 1997;179(17):5465–70. doi: 10.1128/jb.179.17.5465-5470.1997. PubMed PMID: 9287002; PubMed Central PMCID: PMCPMC179418.

47. Liu X, den Blaauwen T. NlpI-Prc proteolytic complex mediates peptidoglycan synthesis and degradation via regulation of hydrolases and synthases in *Escherichia coli*. Int J Mol Sci. 2023;24(22). Epub 20231115. doi: 10.3390/ijms242216355. PubMed PMID: 38003545; PubMed Central PMCID: PMCPMC10671308.

48. Singh SK, Parveen S, SaiSree L, Reddy M. Regulated proteolysis of a cross-link-specific peptidoglycan hydrolase contributes to bacterial morphogenesis. Proc Natl Acad Sci U S A. 2015;112(35):10956–61. Epub 20150817. doi: 10.1073/pnas.1507760112. PubMed PMID: 26283368; PubMed Central PMCID: PMCPMC4568209.

49. Yahashiri A, Jorgenson MA, Weiss DS. Bacterial SPOR domains are recruited to septal peptidoglycan by binding to glycan strands that lack stem peptides. Proc Natl Acad Sci U S A. 2015;112(36):11347–52. Epub 20150824. doi: 10.1073/pnas.1508536112. PubMed PMID: 26305949; PubMed Central PMCID: PMCPMC4568695.

50. Ursinus A, van den Ent F, Brechtel S, de Pedro M, Holtje JV, Lowe J, et al. Murein (peptidoglycan) binding property of the essential cell division protein FtsN from *Escherichia coli*. J Bacteriol. 2004;186(20):6728–37. doi: 10.1128/JB.186.20.6728-6737.2004. PubMed PMID: 15466024; PubMed Central PMCID: PMCPMC522186.

51. Yahashiri A, Jorgenson MA, Weiss DS. The SPOR domain, a widely conserved peptidoglycan binding domain that targets proteins to the site of cell division. J Bacteriol. 2017;199(14). Epub 20170627. doi: 10.1128/JB.00118-17. PubMed PMID: 28396350; PubMed Central PMCID: PMCPMC5494741.

52. Müller P, Ewers C, Bertsche U, Anstett M, Kallis T, Breukink E, et al. The essential cell division protein FtsN interacts with the murein (peptidoglycan) synthase PBP1B in *Escherichia coli*. J Biol Chem. 2007;282(50):36394–402. Epub 20071015. doi: 10.1074/jbc.M706390200. PubMed PMID: 17938168.

53. Arends SJ, Williams K, Scott RJ, Rolong S, Popham DL, Weiss DS. Discovery and characterization of three new *Escherichia coli* septal ring proteins that contain a SPOR domain: DamX, DedD, and RlpA. J Bacteriol. 2010;192(1):242–55. doi: 10.1128/JB.01244-09. PubMed PMID: 19880599; PubMed Central PMCID: PMCPMC2798263.

54. Heidrich C, Ursinus A, Berger J, Schwarz H, Holtje JV. Effects of multiple deletions of murein hydrolases on viability, septum cleavage, and sensitivity to large toxic molecules in *Escherichia coli*. J Bacteriol. 2002;184(22):6093–9. doi: 10.1128/JB.184.22.6093-6099.2002. PubMed PMID: 12399477; PubMed Central PMCID: PMCPMC151956.

55. Georgopapadakou NH, Smith SA, Sykes RB. Mode of action of azthreonam. Antimicrob Agents Chemother. 1982;21(6):950–6. doi: 10.1128/AAC.21.6.950. PubMed PMID: 6180685; PubMed Central PMCID: PMCPMC182051.

56. de Pedro MA, Quintela JC, Holtje JV, Schwarz H. Murein segregation in *Escherichia coli*. J Bacteriol. 1997;179(9):2823–34. doi: 10.1128/jb.179.9.2823-2834.1997. PubMed PMID: 9139895; PubMed Central PMCID: PMCPMC179041.

57. Yakhnina AA, Bernhardt TG. The Tol-Pal system is required for peptidoglycan-cleaving enzymes to complete bacterial cell division. Proc Natl Acad Sci U S A. 2020;117(12):6777–83. Epub 20200309. doi: 10.1073/pnas.1919267117. PubMed PMID: 32152098; PubMed Central PMCID: PMCPMC7104345.

58. Cascales E, Lloubès R, Sturgis JN. The TolQ–TolR proteins energize TolA and share homologies with the flagellar motor proteins MotA–MotB. Molecular microbiology. 2001;42(3):795–807. doi: 10.1046/j.1365-2958.2001.02673.x.

59. Uehara T, Parzych KR, Dinh T, Bernhardt TG. Daughter cell separation is controlled by cytokinetic ring-activated cell wall hydrolysis. EMBO J. 2010;29(8):1412–22. Epub 20100318. doi: 10.1038/emboj.2010.36. PubMed PMID: 20300061; PubMed Central PMCID: PMCPMC2868575.

60. Kaus GM, Snyder LF, Muh U, Flores MJ, Popham DL, Ellermeier CD. Lysozyme resistance in *Clostridioides difficile* is dependent on two peptidoglycan deacetylases. J Bacteriol. 2020;202(22). Epub 20201022. doi: 10.1128/JB.00421-20. PubMed PMID: 32868404; PubMed Central PMCID: PMCPMC7585060.

61. Dobihal GS, Flores-Kim J, Roney IJ, Wang X, Rudner DZ. The WalR-WalK signaling pathway modulates the activities of both CwlO and LytE through control of the peptidoglycan deacetylase PdaC in *Bacillus subtilis*. J Bacteriol. 2022;204(2):e0053321. Epub 20211206. doi: 10.1128/JB.00533-21. PubMed PMID: 34871030; PubMed Central PMCID: PMCPMC8846395.

62. Arnaouteli S, Giastas P, Andreou A, Tzanodaskalaki M, Aldridge C, Tzartos SJ, et al. Two putative polysaccharide deacetylases are required for osmotic stability and cell shape maintenance in *Bacillus anthracis*. J Biol Chem. 2015;290(21):13465–78. Epub 20150330. doi: 10.1074/jbc.M115.640029. PubMed PMID: 25825488; PubMed Central PMCID: PMCPMC4505593.

63. Coullon H, Rifflet A, Wheeler R, Janoir C, Boneca IG, Candela T. N-Deacetylases required for muramic-delta-lactam production are involved in *Clostridium difficile* sporulation, germination, and heat resistance. J Biol Chem. 2018;293(47):18040–54. Epub 20180928. doi: 10.1074/jbc.RA118.004273. PubMed PMID: 30266804; PubMed Central PMCID: PMCPMC6254358.

64. Benachour A, Ladjouzi R, Le Jeune A, Hebert L, Thorpe S, Courtin P, et al. The lysozyme-induced peptidoglycan N-acetylglucosamine deacetylase PgdA (EF1843) is required for *Enterococcus faecalis* virulence. J Bacteriol. 2012;194(22):6066–73. Epub 20120907. doi: 10.1128/JB.00981-12. PubMed PMID: 22961856; PubMed Central PMCID: PMCPMC3486378.

65. Corbin Goodman LC, Erickson HP. FtsZ at mid-cell is essential in *Escherichia coli* until the late stage of constriction. Microbiology (Reading). 2022;168(6). doi: 10.1099/mic.0.001194. PubMed PMID: 35679326.

66. Silber N, Mayer C, Matos de Opitz CL, Sass P. Progression of the late-stage divisome is unaffected by the depletion of the cytoplasmic FtsZ pool. Commun Biol. 2021;4(1):270. Epub 20210301. doi: 10.1038/s42003-021-01789-9. PubMed PMID: 33649500; PubMed Central PMCID: PMCPMC7921118.

67. Baba T, Ara T, Hasegawa M, Takai Y, Okumura Y, Baba M, et al. Construction of *Escherichia coli* K-12 in-frame, single-gene knockout mutants: the Keio collection. Mol Syst Biol. 2006;2:2006 0008. Epub 20060221. doi: 10.1038/msb4100050. PubMed PMID: 16738554; PubMed Central PMCID: PMCPMC1681482.

68. Datsenko KA, Wanner BL. One-step inactivation of chromosomal genes in *Escherichia coli* K- 12 using PCR products. Proc Natl Acad Sci U S A. 2000;97(12):6640–5. doi: 10.1073/pnas.120163297. PubMed PMID: 10829079; PubMed Central PMCID: PMCPMC18686.

69. Seco EM, Fernandez LA. Efficient markerless integration of genes in the chromosome of probiotic *E. coli* Nissle 1917 by bacterial conjugation. Microb Biotechnol. 2022;15(5):1374–91. Epub 20211109. doi: 10.1111/1751-7915.13967. PubMed PMID: 34755474; PubMed Central PMCID: PMCPMC9049610.

70. Reddy M. Role of FtsEX in cell division of *Escherichia coli*: viability of *ftsEX* mutants is dependent on functional SufI or high osmotic strength. J Bacteriol. 2007;189(1):98–108. Epub 20061027. doi: 10.1128/JB.01347-06. PubMed PMID: 17071757; PubMed Central PMCID: PMCPMC1797223.

71. Silhavy TJ, Berman ML, Enquist LW, Laboratory CSH. Experiments with gene fusions: Cold Spring Harbor Laboratory; 1984.

72. Pennartz A, Genereux C, Parquet C, Mengin-Lecreulx D, Joris B. Substrate-induced inactivation of the *Escherichia coli* AmiD N-acetylmuramoyl-L-alanine amidase highlights a new strategy to inhibit this class of enzyme. Antimicrob Agents Chemother. 2009;53(7):2991–7. Epub 20090223. doi: 10.1128/AAC.01520-07. PubMed PMID: 19237650; PubMed Central PMCID: PMCPMC2704666.

73. Van Straaten KE, Dijkstra BW, Thunnissen AM. Purification, crystallization and preliminary X- ray analysis of the lytic transglycosylase MltA from *Escherichia coli*. Acta Crystallogr D Biol Crystallogr. 2004;60(Pt 4):758–60. Epub 20040323. doi: 10.1107/S0907444904002574. PubMed PMID: 15039577.

74. Bertsche U, Kast T, Wolf B, Fraipont C, Aarsman ME, Kannenberg K, et al. Interaction between two murein (peptidoglycan) synthases, PBP3 and PBP1B, in *Escherichia coli*. Molecular microbiology. 2006;61(3):675–90. Epub 2006/06/29. doi: 10.1111/j.1365-2958.2006.05280.x. PubMed PMID: 16803586.

75. Egan AJF, Jean NL, Koumoutsi A, Bougault CM, Biboy J, Sassine J, et al. Outer-membrane lipoprotein LpoB spans the periplasm to stimulate the peptidoglycan synthase PBP1B. Proceedings of the National Academy of Sciences of the United States of America. 2014;111(22):8197–202. doi: 10.1073/pnas.1400376111. PubMed PMID: PMC4050580.

76. Breukink E, van Heusden HE, Vollmerhaus PJ, Swiezewska E, Brunner L, Walker S, et al. Lipid II is an intrinsic component of the pore induced by nisin in bacterial membranes. J Biol Chem. 2003;278(22):19898–903. Epub 2003/03/29. doi: 10.1074/jbc.M301463200. PubMed PMID: 12663672.

77. Bertsche U, Breukink E, Kast T, Vollmer W. In vitro murein (peptidoglycan) synthesis by dimers of the bifunctional transglycosylase-transpeptidase PBP1B from *Escherichia coli*. J Biol Chem. 2005;280(45):38096–101. Epub 20050909. doi: 10.1074/jbc.M508646200. PubMed PMID: 16154998.

78. Biboy J, Bui NK, Vollmer W. *In vitro* peptidoglycan synthesis assay with lipid II substrate. In: Delcour AH, editor. Bacterial Cell Surfaces: Methods and Protocols. Totowa, NJ: Humana Press; 2013. p. 273-88.

79. Cutler KJ, Stringer C, Lo TW, Rappez L, Stroustrup N, Brook Peterson S, et al. Omnipose: a high- precision morphology-independent solution for bacterial cell segmentation. Nat Methods. 2022;19(11):1438–48. Epub 20221017. doi: 10.1038/s41592-022-01639-4. PubMed PMID: 36253643; PubMed Central PMCID: PMCPMC9636021.

80. Morè N, Martorana AM, Biboy J, Otten C, Winkle M, Serrano CKG, et al. Peptidoglycan remodeling enables *Escherichia coli* to survive severe outer membrane assembly defect. mBio. 2019;10(1). Epub 20190205. doi: 10.1128/mBio.02729-18. PubMed PMID: 30723128; PubMed Central PMCID: PMCPMC6428754.

81. Lyu Z, Yahashiri A, Yang X, McCausland JW, Kaus GM, McQuillen R, et al. FtsN maintains active septal cell wall synthesis by forming a processive complex with the septum-specific peptidoglycan synthases in *E. coli*. Nat Commun. 2022;13(1):5751. Epub 20220930. doi: 10.1038/s41467-022-33404-8. PubMed PMID: 36180460; PubMed Central PMCID: PMCPMC9525312.

82. Neidhardt FC, Bloch PL, Smith DF. Culture medium for enterobacteria. J Bacteriol. 1974;119(3):736–47. doi: 10.1128/jb.119.3.736-747.1974. PubMed PMID: 4604283; PubMed Central PMCID: PMCPMC245675.

83. Letunic I, Bork P. Interactive Tree of Life (iTOL) v6: recent updates to the phylogenetic tree display and annotation tool. Nucleic Acids Res. 2024;52(W1):W78–W82. doi: 10.1093/nar/gkae268. PubMed PMID: 38613393; PubMed Central PMCID: PMCPMC11223838.

84. Jumper J, Evans R, Pritzel A, Green T, Figurnov M, Ronneberger O, et al. Highly accurate protein structure prediction with AlphaFold. Nature. 2021;596(7873):583-9. Epub 20210715. doi: 10.1038/s41586-021-03819-2. PubMed PMID: 34265844; PubMed Central PMCID: PMCPMC8371605.

85. Evans R, O’Neill M, Pritzel A, Antropova N, Senior A, Green T, et al. Protein complex prediction with AlphaFold-Multimer. bioRxiv. 2022. doi: 10.1101/2021.10.04.463034.

86. Berman HM, Westbrook J, Feng Z, Gilliland G, Bhat TN, Weissig H, et al. The Protein Data Bank. Nucleic Acids Res. 2000;28(1):235–42. doi: 10.1093/nar/28.1.235. PubMed PMID: 10592235; PubMed Central PMCID: PMCPMC102472.

87. Abramson J, Adler J, Dunger J, Evans R, Green T, Pritzel A, et al. Accurate structure prediction of biomolecular interactions with AlphaFold 3. Nature. 2024;630(8016):493-500. Epub 20240508. doi: 10.1038/s41586-024-07487-w. PubMed PMID: 38718835; PubMed Central PMCID: PMCPMC11168924.

88. Goddard TD, Huang CC, Meng EC, Pettersen EF, Couch GS, Morris JH, et al. UCSF ChimeraX: Meeting modern challenges in visualization and analysis. Protein Sci. 2018;27(1):14–25. Epub 20170906. doi: 10.1002/pro.3235. PubMed PMID: 28710774; PubMed Central PMCID: PMCPMC5734306.

89. Gordon JC, Myers JB, Folta T, Shoja V, Heath LS, Onufriev A. H++: a server for estimating pKas and adding missing hydrogens to macromolecules. Nucleic Acids Res. 2005;33(Web Server issue):W368–71. doi: 10.1093/nar/gki464. PubMed PMID: 15980491; PubMed Central PMCID: PMCPMC1160225.

90. Case DA, Aktulga HM, Belfon K, Cerutti DS, Cisneros GA, Cruzeiro VWD, et al. AmberTools. J Chem Inf Model. 2023;63(20):6183–91. doi: 10.1021/ACS.JCIM.3C01153.

91. Tian C, Kasavajhala K, Belfon KAA, Raguette L, Huang H, Migues AN, et al. ff19SB: amino-acid- specific protein backbone parameters trained against quantum mechanics energy surfaces in solution. J Chem Theory Comput. 2020;16(1):528–52. Epub 20191203. doi: 10.1021/acs.jctc.9b00591. PubMed PMID: 31714766.

92. Izadi S, Anandakrishnan R, Onufriev AV. Building water models: a different approach. J Phys Chem Lett. 2014;5(21):3863–71. Epub 20141016. doi: 10.1021/jz501780a. PubMed PMID: 25400877; PubMed Central PMCID: PMCPMC4226301.

93. Hopkins CW, Le Grand S, Walker RC, Roitberg AE. Long-time-step molecular dynamics through hydrogen mass repartitioning. J Chem Theory Comput. 2015;11(4):1864–74. Epub 20150330. doi: 10.1021/ct5010406. PubMed PMID: 26574392.

94. Shirts MR, Klein C, Swails JM, Yin J, Gilson MK, Mobley DL, et al. Lessons learned from comparing molecular dynamics engines on the SAMPL5 dataset. J Comput Aided Mol Des. 2017;31(1):147–61. Epub 20161027. doi: 10.1007/s10822-016-9977-1. PubMed PMID: 27787702; PubMed Central PMCID: PMCPMC5581938.

95. Loncharich RJ, Brooks BR, Pastor RW. Langevin dynamics of peptides: the frictional dependence of isomerization rates of N-acetylalanyl-N’-methylamide. Biopolymers. 1992;32(5):523–35. doi: 10.1002/bip.360320508. PubMed PMID: 1515543.

96. Åqvist J, Wennerström P, Nervall M, Bjelic S, Brandsdal BO. Molecular dynamics simulations of water and biomolecules with a Monte Carlo constant pressure algorithm. Chemical Physics Letters. 2004;384(4-6):288–94. doi: 10.1016/j.cplett.2003.12.039.

97. Darden T, York D, Pedersen L. Particle mesh Ewald: An N⋅log(N) method for Ewald sums in large systems. The Journal of Chemical Physics. 1993;98(12):10089–92. doi: 10.1063/1.464397.

98. Roe DR, Cheatham TE, 3rd. PTRAJ and CPPTRAJ: Software for processing and analysis of molecular dynamics trajectory data. J Chem Theory Comput. 2013;9(7):3084–95. Epub 20130625. doi: 10.1021/ct400341p. PubMed PMID: 26583988.

